# A Comprehensive Atlas of Perineuronal Net Distribution and Colocalization with Parvalbumin in the Adult Mouse Brain

**DOI:** 10.1101/2023.01.24.525313

**Authors:** Leonardo Lupori, Valentino Totaro, Sara Cornuti, Luca Ciampi, Fabio Carrara, Edda Grilli, Aurelia Viglione, Francesca Tozzi, Elena Putignano, Raffaele Mazziotti, Giuseppe Amato, Claudio Gennaro, Paola Tognini, Tommaso Pizzorusso

**Affiliations:** BIO@SNS lab, Scuola Normale Superiore, 56126 Pisa, Italy; Institute of Information Science and Technologies (ISTI-CNR), 56124 Pisa, Italy; Department of Biology, University of Pisa, 56126 Pisa, Italy; Institute of Neuroscience (IN-CNR), 56124 Pisa, Italy; Department of Translational Research and New Technologies in Medicine and Surgery, University of Pisa, 56126 Pisa, Italy

**Author notes:** These authors contributed equally to this work (Leonardo Lupori, Valentino Totaro).

## Abstract

Perineuronal nets (PNNs) surround specific neurons in the brain and are involved in various forms of plasticity and clinical conditions. However, our understanding of the PNN role in these phenomena is limited by the lack of highly quantitative maps of PNN distribution and association with specific cell types. Here, we present the first comprehensive atlas of PNN distribution (in Allen Brain Atlas coordinates) and colocalization with parvalbumin (PV) cells for over 600 regions of the adult mouse brain. Data analysis showed that PV expression is a good predictor of PNN aggregation. In the cortex, PNNs are dramatically enriched in layer 4 of all primary sensory areas in correlation with thalamocortical input density, and their distribution mirrors intracortical connectivity patterns. Gene expression analysis identified many PNN correlated genes. Strikingly, PNN anticorrelated transcripts were enriched in synaptic plasticity genes, generalizing PNN role as circuit stability factors. Overall, this atlas offers novel resources for understanding the organizational principles of the brain extracellular matrix.

## Introduction

Perineuronal Nets (PNNs) are specialized reticular structures of the extracellular matrix (ECM) that ensheath neurons in the entire mouse and human brain (Galtrey et al., 2008; Hendry et al., 1988; Seeger et al., 1994; Köppe et al., 1997). These structures aggregate progressively during postnatal development, in parallel with the closure of critical periods for developmental plasticity (Pizzorusso et al., 2002; Boggio et al., 2019; Reichelt et al., 2019; Ye et al., 2013). Although their precise composition may vary between different brain regions, PNNs are known to share three essential molecular constituents: hyaluronic acid, glycosylated proteins called chondroitin-sulfate proteoglycans (CSPGs), and link proteins such as hyaluronan and proteoglycan link protein 1 (HAPLN1) and Tenascin-R (Carulli et al., 2010; Dauth et al., 2016; Kwok et al., 2010). The sugars present on CSPGs are also the binding target of the lectin Wisteria floribunda agglutinin (WFA), the most widely used marker to visualize PNNs in histological analyses (Fawcett et al., 2019; Härtig et al., 1999).

The precise contribution of PNNs in regulating brain function is a strongly active area of research. Many roles have been proposed, but a key overarching theme is that PNNs tightly control the plasticity and stability of neuronal circuits (Fawcett et al., 2022; Nabel et al., 2013). This function has been studied throughout many cortical and subcortical regions of the brain. For example, PNNs are known to control ocular dominance plasticity in the visual cortex (Pizzorusso et al., 2002; Carulli et al., 2010; Miyata et al., 2012; Rowlands et al., 2018; Beurdeley et al., 2012), fear memory extinction in the amygdala (Gogolla et al., 2009), spatial representation stability of grid cells in the entorhinal cortex (Christensen et al., 2021), associative motor learning in the cerebellum (Carulli et al., 2020), and social memory in the hippocampus (Cope et al., 2021; Domínguez et al., 2019). Enzymatic digestion of PNNs has been shown to promote plasticity and improve recovery after damage to the central nervous system (Bradbury et al., 2002). Additionally, PNNs are thought to stabilize neuronal circuitry by protecting fast-spiking neurons against oxidative stress (Cabungcal et al., 2013) a risk factor for psychiatric diseases. Abnormalities in PNNs that make PV cells more susceptible to oxidative damage have been reported in schizophrenic patients (Pantazopoulos et al., 2010).

Despite these general features, PNNs also show a remarkable degree of variability between different brain regions both in terms of structure and function (Ueno et al., 2018). In the isocortex, several studies showed that PNNs primarily surround parvalbumin-expressing (PV) fast-spiking GABAergic interneurons. However, in the hippocampal CA2 and in other areas, they also ensheath excitatory pyramidal neurons, suggesting a different biological function in these regions (Carstens et al., 2016). At the functional level, the enzymatic removal of PNNs can have different effects (Wingert et al., 2021). For example, it enhances LTD in the perirhinal cortex (Romberg et al., 2013), while it impairs both early-phase LTP and LTD in the hippocampus (Bukalo et al., 2007). The lack of understanding of the principles of PNN organization throughout the brain hinders our comprehension of their functional role and possible therapeutic implications. Furthermore, the extent to which PNNs are linked to PV cells across brain areas has not been systematically studied.

Here, we present a systematic brain-wide analysis of PNNs and PV neurons in the mouse brain. We provide multiple quantitative measurements for PNNs, PV cells, and their interaction for more than 600 different brain areas. We also release two deep learning models, pre-trained on a dataset of approximately 0.8 million manually annotated PNNs and PV cells, for their automatic detection. Finally, we demonstrate that, thanks to our dataset, it is possible to detect connectivity and gene expression patterns that correlate with the presence of PNNs. We believe that these resources will have a significant impact on facilitating research on PNNs.

## Results

### PNN and PV cells quantification in the mouse brain

We performed immuno-/lectin histochemistry on serially collected whole-brain coronal slices of seven adult mice, staining sections with both WFA and an anti-PV antibody (***Figure 1***A). We then acquired fluorescence images and registered them to the Allen Institute CCFv3.

**Figure 1.**
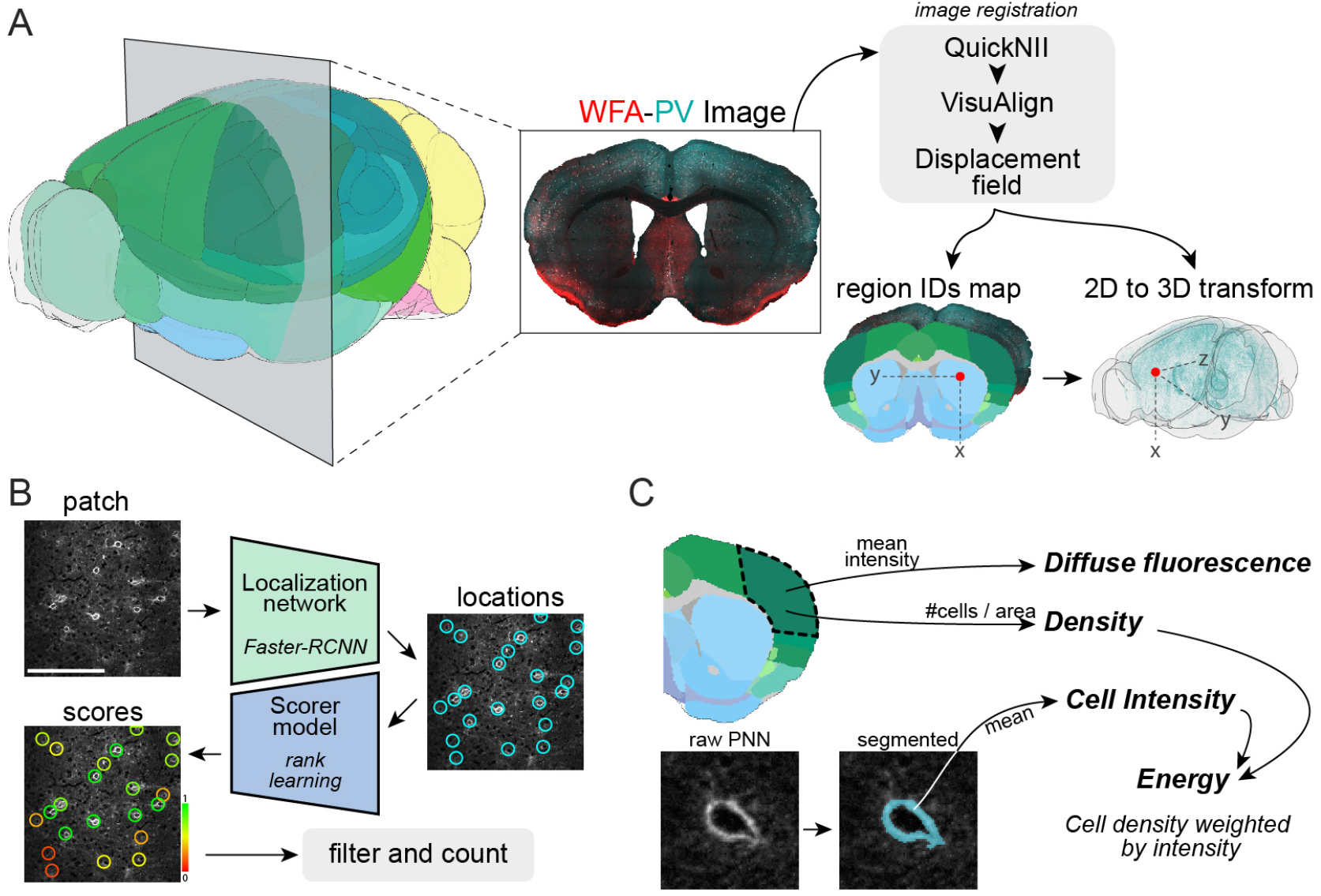
Image registration and analysis pipeline. (**A**) Schematic of the pipeline for slice registration to the Allen Institute CCFv3 reference volume. (**A**) Schematic of the strategy for cell counting. Two different modules were used, a larger convolutional neural network for localization and a smaller one for scoring. Scale bar: 200μm. (**A**) Diagram showing a graphical explanation of the four metrics used to quantify PNN and PV staining.

To automatically detect the (x,y) coordinates of PNNs and PV cells, we trained two deep convolutional neural networks with a dataset comprising roughly 0.67 million manually annotated PNNs and 0.16 million PV cells (***Figure 1***B). While manually counting non-trivial structures on a large scale, an experimenter can be influenced by illumination conditions, fatigue, or different judgements, spanning from conservative to liberal. As a result, the training dataset can inherit annotation biases. To address this issue, we implemented a second stage whereby we assigned a confidence score to each object detected by the two deep neural networks. This scorer module consisted of other two deep-learning models trained on two smaller datasets (4,727 PNNs and 5,738 PV cells) labeled by seven independent expert raters. The aim was to produce scores for each putative object that maximally correlate with the raters’ agreement. A detailed description of this method is available in Ciampi et al., 2022.

In our multi-rater dataset, the average agreement(Jaccard index) between pairs of expert raters was 64% for PNNs and 72% for PV cells, demonstrating relevant individual differences in counting strategies (***Figure S1***A). Our scoring models produced detection scores that strongly correlated with the number of raters that detected each object (***Figure S1***B, C). Overall, when tested on objects located by at least three raters, our models proved to be reliable in the detection of PNNs and PV cells (see Ciampi et al., 2022 and section Deep learning models for cell counting in Methods & Materials). We release the pre-trained four models used in this study ([link](https://github.com/ciampluca/counting_perineuronal_nets/releases)) to allow performing predictions on new images or to fine-tune them based on different experimental setups.

To quantify PV and WFA staining, we defined a set of metrics describing either “general” or “cellular” aspects of the staining signal (***Figure 1***C). To quantify general staining intensity in a region, we defined *diffuse fluorescence* as the average pixel intensity value in that region. This measure includes the signal coming from both interstitial CSPGs diffusely present in the ECM, and from CSPGs aggregated in PNNs. To quantify “cellular” aspects (either single PV cells or aggregated, cell-ensheathing, PNNs), we first defined *density*, corresponding to the number of objects per unit of surface area. We then measured the intensity of each individual PNN and PV cell by averaging the values of the pixels belonging to the object, segmented from a small (80×80 pixels) patch centered on its (x,y) coordinates. Based on this measurement, we defined *cell intensity*, expressing the average staining intensity of individual PNNs or PV cells in a region. Finally, we reasoned that the functional relevance of PNNs or PV cells might be better represented by a single metric that integrates both the density and the intensity of cells. We thus defined *energy*, as the density multiplied by the average cell intensity, a metric analogous to the one used by the Allen Institute in Lein et al., 2007 (***Figure 1***C, see section Staining metrics definitions in Methods & Materials for details). As a result, a region with more and brighter PNNs would have increased PNN energy. *Diffuse fluorescence* and *energy* were normalized within each mouse by dividing them by their respective value calculated on the entire brain. As a result, a value of 1 equals the brain’s average and, importantly, the two metrics have the same scale. In the rest of the paper, we will use the metrics *diffuse fluorescence* and *energy* respectively as a “general” and “cellular” measurement.

### Distribution of PNNs across the mouse brain

To describe the distribution of PNNs in the entire brain, we first aggregated data in 12 major brain subdivisions (***Figure 2***A). These regions had highly different values for both WFA diffuse fluorescence and PNN energy with particular enrichment in the cortex and in posterior areas of the brain (***Figure 2**A, B*, see ***Table 1*** for statistical comparisons). We then analyzed PNN energy, representing aggregated PNNs in a region. Using this metric, the differences between the studied areas were more pronounced than those observed in measurements of diffuse fluorescence (***Figure 2***A, B). These data indicate that there is a non homogeneous expression of diffuse WFA staining and PNNs in the brain that is already evident at this macroscopic level of analysis.

**Figure 2.**
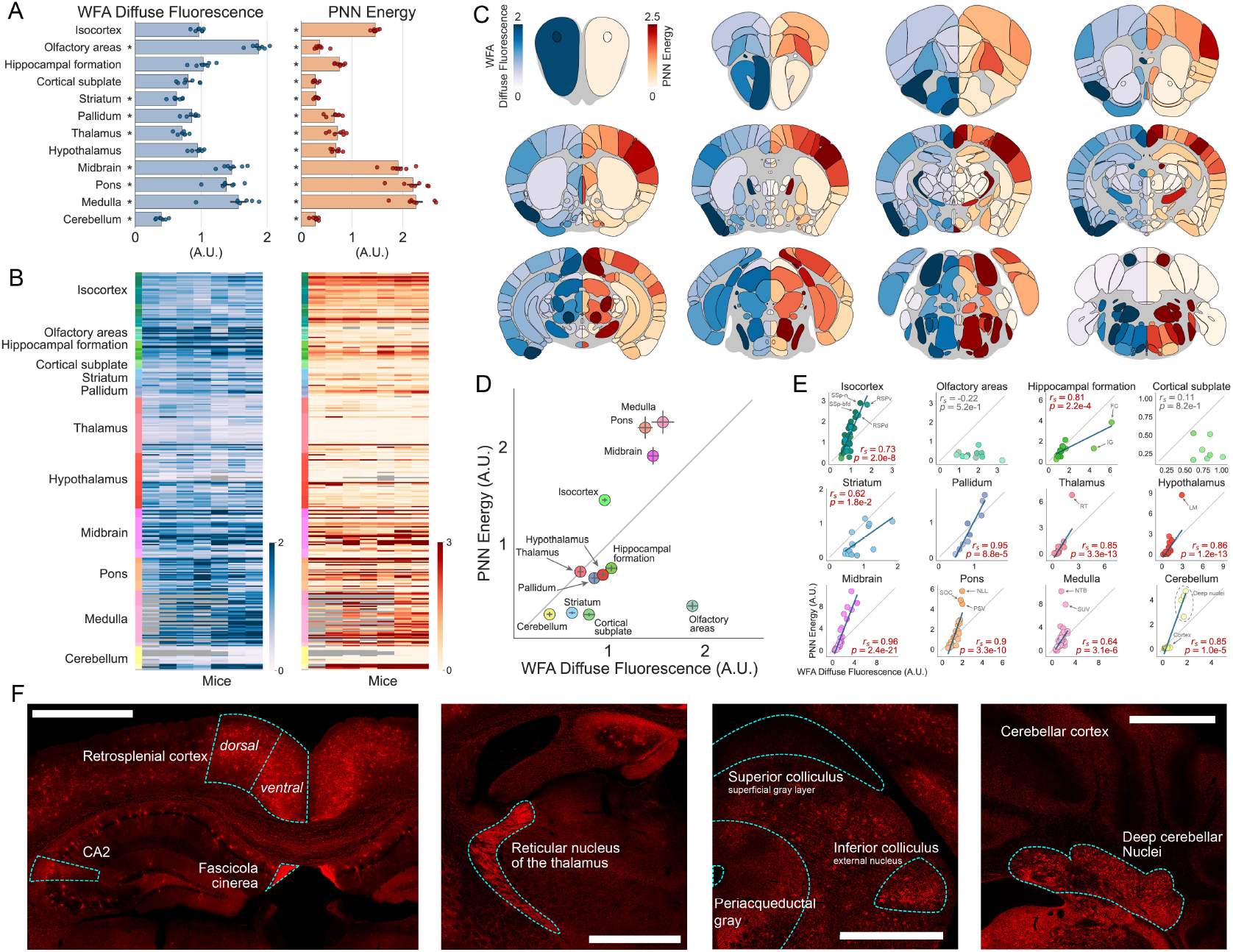
Distribution of WFA-positive PNNs throughout the entire mouse brain. (**A**) Quantification of diffuse fluorescence and PNN energy for 12 major brain subdivisions. Asterisks indicate subdivisions significantly different from the brain average (value of 1. See ***Table 1*** for statistical comparisons). (**B**) Heatmaps showing staining metrics for mid-ontology brain regions in individual mice. Grayed-out cells represent regions where data are not available due to no sampling of that region. (**C**) Heatmaps showing coronal sections of the brain, sliced at different anteroposterior locations. On the left hemisphere (blue colormap) is displayed average diffuse WFA fluorescence, while on the right hemisphere (red colormap) is displayed average PNN energy for each brain region. (**D**) Plots of PNN energy versus WFA diffuse fluorescence for each of the 12 major brain subdivisions. (**E**) Same as in D but data is split in each brain region of the 12 major brain subdivisions. (**F**) Representative WFA staining in a selection of brain areas. Scalebar: 1mm. Error bars in A and D represent SEM across mice. Dots in A represent mice, in D and E, represent brain regions. In E, text insets indicate the Spearman correlation coefficient (*r_s_*) and the corresponding p-value, the gray line indicates the X-Y bisector, and, for significant correlations highlighted in red, the blue line shows the best linear fit.

**Table 1.** Statistical comparisons.

| Fig | Description | Test | N (units) | Results |
| --- | --- | --- | --- | --- |
| 2A | Diffuse fluorescence differences between major brain subdivision | one-way RM ANOVA | 7 mice per brain region | $F(11,66)=62.45$, $P<0.0001$ |
| 2A | PNN energy differences between major brain subdivision | one-way RM ANOVA | 7 mice per brain region | $F(11,66)=143.1$, $P<0.0001$ |
| 2A | WFA Diffuse fluorescence significantly different from 1 |  |  |  |
| | Isocortex | one sample t-test | 7 (mice) | $t(6)=1.40$, $P=0.21$ |
| | Olfactory areas | one sample t-test | 7 (mice) | $t(6)=17.87$, $P<0.001$ |
| | Hippocampal formation | one sample t-test | 7 (mice) | $t(6)=0.59$, $P=0.58$ |
| | Cortical subplate | one sample t-test | 7 (mice) | $t(6)=3.56$, $P=0.01$ |
| | Striatum | one sample t-test | 7 (mice) | $t(6)=11.49$, $P<0.001$ |
| | Pallidum | one sample t-test | 7 (mice) | $t(6)=3.84$, $P=0.008$ |
| | Thalamus | one sample t-test | 7 (mice) | $t(6)=9.28$, $P<0.001$ |
| | Hypothalamus | one sample t-test | 7 (mice) | $t(6)=1.66$, $P=0.15$ |
| | Midbrain | one sample t-test | 7 (mice) | $t(6)=7.52$, $P<0.001$ |
| | Pons | one sample t-test | 7 (mice) | $t(6)=4.97$, $P=0.003$ |
| | Medulla | one sample t-test | 7 (mice) | $t(6)=4.84$, $P=0.003$ |
| | Cerebellum | one sample t-test | 7 (mice) | $t(6)=21.60$, $P<0.001$ |
| 2A | PNN energy significantly different from 1 |  |  |  |
| | Isocortex | one sample t-test | 7 (mice) | $t(6)=20.67$, $P<0.001$ |
| | Olfactory areas | one sample t-test | 7 (mice) | $t(6)=16.48$, $P<0.001$ |
| | Hippocampal formation | one sample t-test | 7 (mice) | $t(6)=8.19$, $P<0.001$ |
| | Cortical subplate | one sample t-test | 7 (mice) | $t(6)=39.88$, $P<0.001$ |
| | Striatum | one sample t-test | 7 (mice) | $t(6)=51.13$, $P<0.001$ |
| | Pallidum | one sample t-test | 7 (mice) | $t(6)=6.77$, $P<0.001$ |
| | Thalamus | one sample t-test | 7 (mice) | $t(6)=4.90$, $P=0.003$ |
| | Hypothalamus | one sample t-test | 7 (mice) | $t(6)=9.34$, $P<0.001$ |
| | Midbrain | one sample t-test | 7 (mice) | $t(6)=10.12$, $P<0.001$ |
| | Pons | one sample t-test | 7 (mice) | $t(6)=10.63$, $P<0.001$ |
| | Medulla | one sample t-test | 7 (mice) | $t(6)=10.46$, $P<0.001$ |
| | Cerebellum | one sample t-test | 7 (mice) | $t(6)=29.42$, $P<0.001$ |
| 4C | Probability of having a PNN (whole brain) | one way RM ANOVA | 7 per class (mice) | see figure inset |
| 4D | Probability of having a PNN (major brain subdivisions) | one way RM ANOVA | 7 per class (mice) | see figure inset |
| 5C | Visual areas. Diffuse fluorescence comparison | Paired t-test | 7 (mice) per group | $t(6)=4.72$, $P=0.003$ |
| | Visual areas. PNN energy comparison | Paired t-test | 7 (mice) per group | $t(6)=8.60$, $P<0.001$ |
| 5D | Visual areas. Comparison by layer | two-way RM ANOVA | 7 (mice) per group | Interaction<br>layer*areaHierarchy<br>$F(4,24)=92.50$, $P<0.0001$ |
| | L1 - Primary vs Associative | Paired T-test, Sidak | 7 (mice) per group | $t(6)=0.42$, $P=0.997$ |
| | L2/3 - Primary vs Associative | Paired T-test, Sidak | 7 (mice) per group | $t(6)=2.58$, $P=0.193$ |
| | L4 - Primary vs Associative | Paired T-test, Sidak | 7 (mice) per group | $t(6)=12.80$, $P<0.001$ |
| | L5 - Primary vs Associative | Paired T-test, Sidak | 7 (mice) per group | $t(6)=10.74$, $P<0.001$ |
| | L6 - Primary vs Associative | Paired T-test, Sidak | 7 (mice) per group | $t(6)=5.05$, $P=0.012$ |
| 5E | Auditory areas. Diffuse fluorescence comparison | Paired t-test | 7 (mice) per group | $t(6)=5.526$, $P=0.002$ |
| | Auditory areas. PNN energy comparison | Paired t-test | 7 (mice) per group | $t(6)=11.33$, $P<0.001$ |
| 5F | Auditory areas. Comparison by layer | two-way RM ANOVA | 7 (mice) per group | Interaction<br>layer*areaHierarchy<br>$F(4,24)=41.13$, $P<0.0001$ |
| | L1 - Primary vs Associative | Paired T-test, Sidak | 7 (mice) per group | $t(6)=1.51$, $P=0.63$ |
| | L2/3 - Primary vs Associative | Paired T-test, Sidak | 7 (mice) per group | $t(6)=4.33$, $P=0.024$ |
| | L4 - Primary vs Associative | Paired T-test, Sidak | 7 (mice) per group | $t(6)=10.45$, $P<0.001$ |
| | L5 - Primary vs Associative | Paired T-test, Sidak | 7 (mice) per group | $t(6)=8.04$, $P=0.001$ |
| | L6 - Primary vs Associative | Paired T-test, Sidak | 7 (mice) per group | $t(6)=2.48$, $P=0.216$ |
| 5G | Somatosensory areas. Diffuse fluorescence comparison | Paired t-test | 7 (mice) per group | $t(6)=0.922$, $P=0.392$ |
|  | Somatosensory areas. PNN energy comparison | Paired t-test | 7 (mice) per group | t(6)=8.42, P<0.001 |
| 5H | Somatosensory areas. Comparison by layer | two-way RM ANOVA | 7 (mice) per group | Interaction layer*areaHierarchy F(4,24) = 15.65, P<0.0001 |
|  | L1 - Primary vs Associative | Paired T-test, Sidak | 7 (mice) per group | t(6)=2.76, P=0.154 |
|  | L2/3 - Primary vs Associative | Paired T-test, Sidak | 7 (mice) per group | t(6)=2.47, P=0.220 |
|  | L4 - Primary vs Associative | Paired T-test, Sidak | 7 (mice) per group | t(6)=5.23, P=0.009 |
|  | L5 - Primary vs Associative | Paired T-test, Sidak | 7 (mice) per group | t(6)=5.37, P=0.008 |
|  | L6 - Primary vs Associative | Paired T-test, Sidak | 7 (mice) per group | t(6)=8.05, P=0.001 |
| 5L | (Inset) - Thalamic input strength different between high- and low-WFA cortical regions | two sample t-test | 25 vs 11 (regions) | t(34)=0.19, P=0.85 |
| 5M | Silhouette score comparison | One-way ANOVA | 7 (mice) per group | F(2)=83.37, P<0.001 |
|  | Low-high-WFA vs Cortical Subnetworks | T-test, Holm-Sidak | 7 (mice) per group | t(6)=9.61, P<0.001 |
|  | Low-high-WFA vs Shuffle | T-test, Holm-Sidak | 7 (mice) per group | t(6)=10.00, P<0.001 |
|  | Cortical Subnetworks vs Shuffle | T-test, Holm-Sidak | 7 (mice) per group | t(6)=2.00, P=0.068 |

We then grouped data in a set of 316 mid-ontology brain regions (***Figure 2***B, ***Figure S2***, for individual areas, see ***Table ST4*** for area acronyms). The profile of both metrics was consistent across individual mice and it showed that individual brain areas have remarkably diverse values for both diffuse fluorescence and PNN energy even within the same major subdivision (***Figure 2***C, F). To visualize the results at this granularity, we plotted the average of both metrics across mice in a series of brain heatmaps coronally sliced at 12 anteroposterior locations (***Figure 2***C).

Intriguingly, both the diffuse and the cellular measurements of PNNs often varied together. However, some areas showed striking differences between the two metrics (***Figure 2***C). Thus, we asked whether the presence of PNNs in an area is always associated with a high level of diffuse WFA staining in all brain regions. To answer this question, we plotted WFA diffuse fluorescence versus PNN energy for all the major brain subdivisions (***Figure2***D). Isocortex, midbrain, pons, and medulla were skewed towards the top-left side of the plot, indicating that they are characterized by strong individual aggregated PNNs, but relatively weak diffuse CSPG signal. Conversely, all the other brain subdivisions showed the opposite effect. Notably, for the olfactory areas, we measured the highest difference between the two metrics, with a strong level of diffuse fluorescence but almost absent aggregated PNNs. We then split these subdivisions into mid-ontology regions and explored the relationship between the two metrics within each group of brain areas (***Figure 2***E). We found that WFA diffuse fluorescence and PNN energy were significantly correlated in all subdivisions except for olfactory areas and the cortical subplate, although the strength of such correlation was not uniform. Striatum had the lowest correlation (*r_s_*=0.62), while Midbrain and Pallidum showed the highest correlation between metrics (*r_s_*=0.96 and 0.95 respectively). These results demonstrate that PNN abundance is not defined at the macrostructure level and that diffuse WFA staining is not necessarily correlated with numerous and strongly labeled PNNs.

Overall, these data represent the first systematic and highly quantitative description of the distribution of WFA-positive PNNs in the entire mouse brain. Raw measurements for individual mice at three levels of anatomical granularity are available in supplementary data SD1.

### Brain-wide analysis of the colocalization between PNNs and PV cells

In the same brain slices used for PNN analysis, we also quantified PV-positive inhibitory interneurons (***Figure 3***A) using the same procedures and metrics used for PNNs (***Figure S3***, data for PV staining in all brain areas are available in supplementary data SD2). PV distribution has been analyzed in previous studies and our results show an overall analogous profile despite methodological differences (Kim et al., 2017; Bjerke et al., 2021). To explore the relationship between PNNs and PV cells in the entire brain, we quantified their colocalization as the percentage of PNNs containing a PV cell (PV^+^ PNNs) or as the percentage of PV cells that are surrounded by a PNN (WFA^+^ PV cells). On average, in the entire brain, 59.1 ±1.0% of PNNs were located around a PV cell, while about one-third of all PV cells in the brain (30.4±1.4%) were surrounded by a PNN. After splitting the data into 12 brain subdivisions, we found that the relationship between PNNs and PV cells was highly heterogeneous (***Figure 3***B). In the isocortex, PNNs surrounded PV cells in more than 70% of the cases, reaching, for example, 81.1 ±0.7% in the retrosplenial cortex (RSPv), 80.8±0.4% in layer 4 of the primary visual cortex (VISp4), and 77.4±0.3% in the anterior cingulate area (ACAv). In all the other major subdivisions, this was the case for at least one-third of the PNNs.

**Figure 3.**
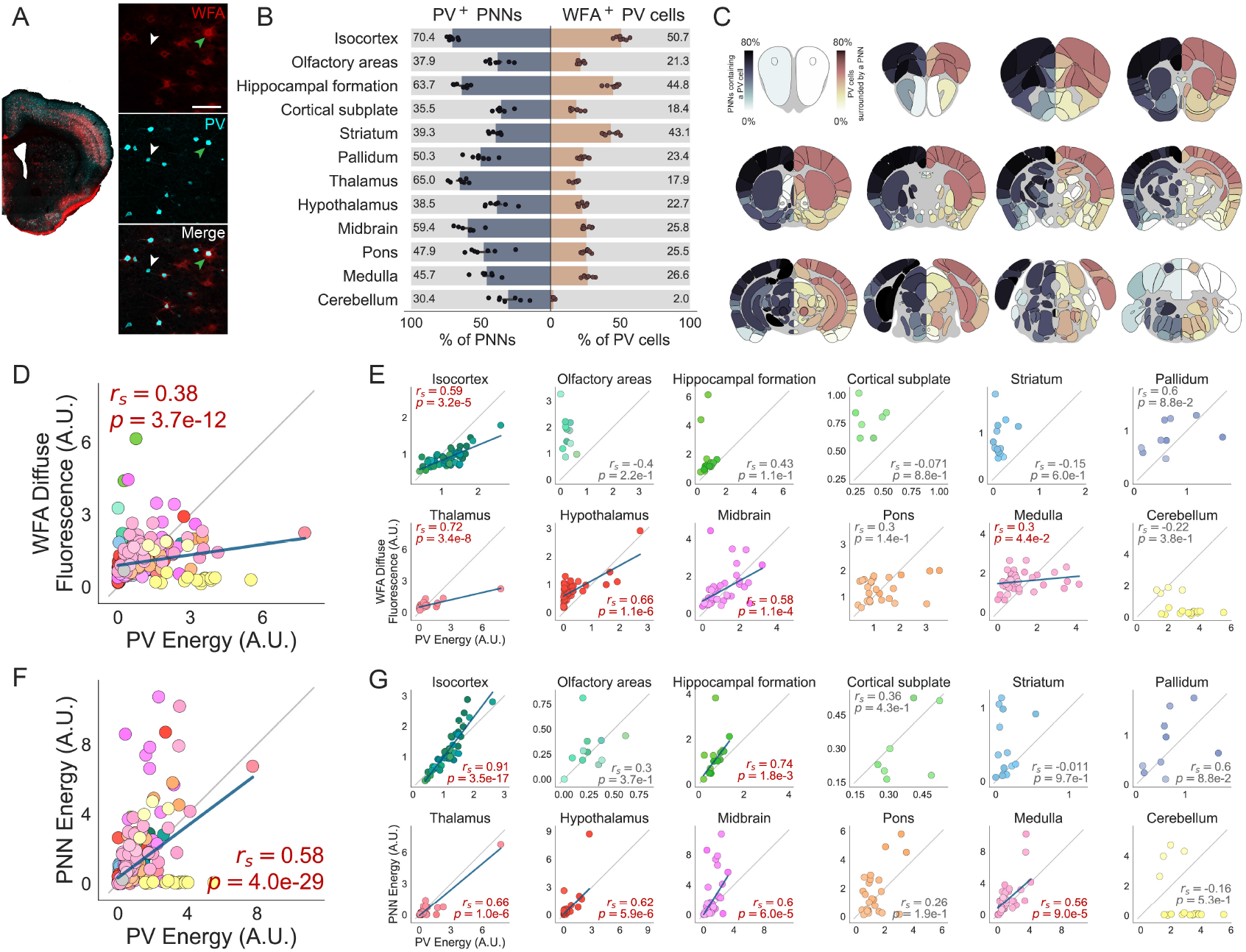
Brain-wide interactions between PNNs and PV cells. (**A**) Representative image of a brain slice stained with WFA (red) and anti-PV (cyan). An inset is magnified on the right, where split channels are also shown. Arrowheads show examples of PV cells without a PNN (white) and colocalized PV-PNNs (green). Scale bar: 100μm. (**B**) Colocalization percentages across 12 major brain subdivisions (on the left, the fraction of PNNs containing a PV cell; on the right, the fraction of PV cells surrounded by a PNN). (**C**) Heatmaps showing coronal sections of the brain, sliced at different anteroposterior locations. On the two hemispheres are represented the percentage of PNNs containing a PV cell (left side) and the percentage of PV cells surrounded by a PNN (right side). (**D**) WFA diffuse fluorescence versus PV energy for all brain areas at a mid-ontology level. (**E**) Same as in D, but areas are split in each major brain subdivision. (**F**) WFA energy versus PV energy for brain areas at a mid-ontology level. (**G**) Same as in F, but areas are split in each major brain subdivision. Error bars in B represent SEM across mice. Dots in B represent mice, while in D, E, F, and G, represent brain areas. Text insets in D, E, F, and G indicate the Spearman correlation coefficient (*r_s_*) and the corresponding p-value, the gray line indicates the X-Y bisector, and, for significant correlations highlighted in red, the blue line shows the best linear fit.

Conversely, analyzing the percentage of PV cells surrounded by a PNN, we observed that in most brain areas, only between 20 and 30% of the PV cells are enwrapped by a WFApositive PNN. A different pattern was present in the isocortex, hippocampal formation, and striatum, where colocalization was much higher (between 40 and 50% of PV cells, reaching for example 71.9±0.4% in VISp4), while in the cerebellum, only few PV-positive cells had a PNN, likely due to the high number of Purkinje cells in the cerebellar cortex that lack PNNs (Baimbridge et al., 1982; Bastianelli, 2003). As before, we also aggregated data in mid-ontology brain regions and measured colocalization metrics in individual areas to reveal patterns with finer granularity (***Figure 3***C, see ***Figure S4*** for data visualization for each region). Colocalization data at three levels of anatomical granularity are available in supplementary data SD3.

Given the high degree of colocalization, we next asked whether PNN and PV staining were correlated across brain regions. To this end, we plotted either WFA diffuse fluorescence (***Figure 3***D) or PNN energy (***Figure 3***F) as a function of PV energy. We found that, throughout all areas of the brain, WFA and PV staining metrics were significantly correlated (***Figure 3***D, F, *r_s_*=0.38 for WFA diffuse vs PV energy, *r_s_*=0.58 for PNN energy vs PV energy). When performing the same analysis at a finer resolution, however, only a subset of brain subdivisions showed a high degree of correlation between WFA and PV (***Figure 3***E, G). The diffuse staining of CSPGs was positively correlated to PV energy in the isocortex, thalamus, hypothalamus, midbrain, and medulla (***Figure 3***E). Interestingly when we compared cellular metrics for both PNNs and PV (PNN energy vs PV energy) correlation coefficients increased, with isocortex showing the most striking trend (***Figure 3***G). Here, PV energy alone was highly predictive of the presence of PNNs (*r_s_*=0.91).

It has been previously reported that two distinct network configurations of PV cells might exist, one more permissive towards plasticity and characterized by weak expression of PV (low-PV), and another that limits plasticity and with strong PV expression (high-PV) (Donato et al., 2013). These two subpopulations likely reflect distinct timing of neurogenesis and connectivity (Donato et al., 2015). Thus, we decided to further explore the relationship between PNNs and PV staining intensity at the level of single cells. First, we looked at the intensity distribution of PNNs and PV cells across our entire dataset. Intriguingly, we found that both PNNs and PV cells had a bimodal intensity distribution (***Figure 4***A, B), suggesting that each could be composed of two subpopulations of high and low expression. Since PNNs are known to inhibit plasticity, we asked if plasticity-inhibiting high-PV cells were more likely to have a PNN. To do this, we grouped all PV cells in four intensity classes of equal width (1:low, 2:intermediate-low, 3:intermediate-high, and 4:high) and measured the probability of being surrounded by a PNN as a function of PV cell intensity. Overall, we found that as PV intensity increased, the probability of having a PNN increased (***Figure 4***C). Repeating the analysis for each brain subdivision, we found that the effect we observed was present in all 12 brain macrostructures except for the hypothalamus, which showed a similar but not statistically significant trend, and the cerebellum (***Figure 4***D). However, the magnitude of such dependency appears to follow three distinct patterns (***Figure 4***E). In isocortex, striatum, and hippocampal formation, PNNs aggregation was strongly and robustly dependent on PV expression. The relationship was inverse in the cerebellum, likely due to the presence of PV-expressing Purkinje cells, and its strength was only moderate for all the other brain areas.

**Figure 4.**
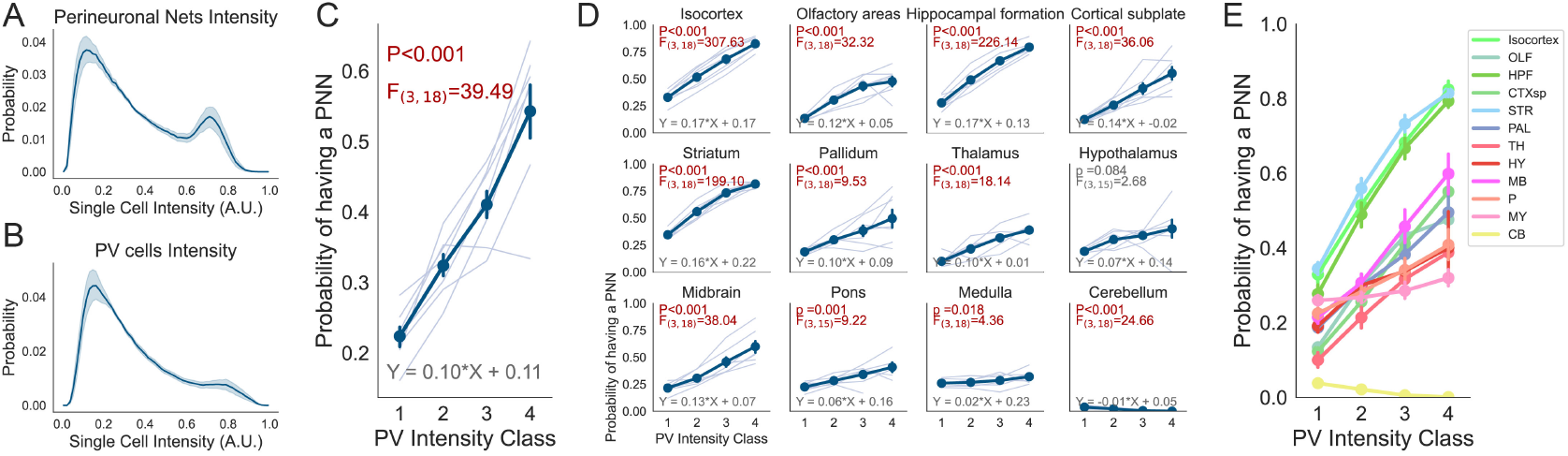
PNN aggregation depends on PV expression levels. (**A**) Probability density function of the intensity of all PNNs. The thick line represents the average, while shading represents SEM across mice (N=7 mice, 69,926±5,235 PNNs per mouse). (**B**) Same as in A but for PV cells (N=7 mice, 13,6479±11,839 PV cells per mouse). (**C**) Probability that a PV cell is surrounded by a PNN as a function of PV intensity class (1: low, 2: intermediate-low, 3: intermediate-high, 4:high) calculated for the whole brain. (**D**) Same as in C, but split in each major brain subdivision. (**E**) Same as in D but all regions are plotted on the same axis. Text insets indicate the result of a one-way RM ANOVA (F statistics and the corresponding p-value), and the estimated parameters of the best first-degree linear fit. Thin lines in C and D represent single mice. Error bars in C, D, and E represent SEM across mice.

Overall these data indicate the existence of a mechanism coupling PV expression with PNN formation. However, the strength of this regulatory mechanism is variable across the brain.

### Primary sensory areas share high levels of PNNs

The precise functional role of PNNs in the cerebral cortex is intensely studied (Fawcett et al., 2019). We reasoned that, by analyzing their expression pattern throughout this anatomical district, we could highlight principles of organization that might explain the spatially inhomogeneous distribution of PNNs. Furthermore, the cerebral cortex is divided into layers with different functional properties and PNN expression. We thus plotted WFA diffuse fluorescence and PNN energy in all cortical regions divided by layer (***Figure 5***A). As previously described, WFA staining was generally more abundant in layers 4 and 5. We noticed that four main groups of regions were characterized by a stronger diffuse WFA staining: somatosensory, visual, and auditory areas, and the retrosplenial cortex (***Figure 5***A). When analyzing aggregated PNNs (PNN energy), this pattern was much sharper and more localized in layer 4 (***Figure 5***A bottom heatmap). Interestingly, PNN energy was particularly high in primary sensory areas (VISp, AUDp, and all SSp areas) while the same enrichment was milder for PV energy (***Figure 5***B). To further investigate this pattern, we isolated each sensory system and aggregated data in primary and associative cortical regions. In the visual cortex, both diffuse fluorescence and PNN energy were lower in associative (VISpor, VISli, VISl, VISpl, VISpm, VISal, VISam, VISrl, VISa) than in primary (VISp) areas (***Figure 5***C) and, splitting data between layers, this effect was present only in layer 4, 5 and 6, and most prominent in layer 4 (***Figure 5***D, ***Figure S5***A). An analogous difference was present in auditory (***Figure 5***E, F, ***Figure S5***B primary (AUDp) versus associative (AUDv, AUDd, AUDpo)) and somatosensory areas (***Figure 5***G, H, ***Figure S5***C, primary (SSp-n, SSp-bfd, SSp-Il, SSp-m, SSp-ul, SSp-tr, SSp-un) versus associative (SSs)) with the exception of diffuse fluorescence in the somatosensory regions of the cortex (***Figure 5***G).

**Figure 5.**
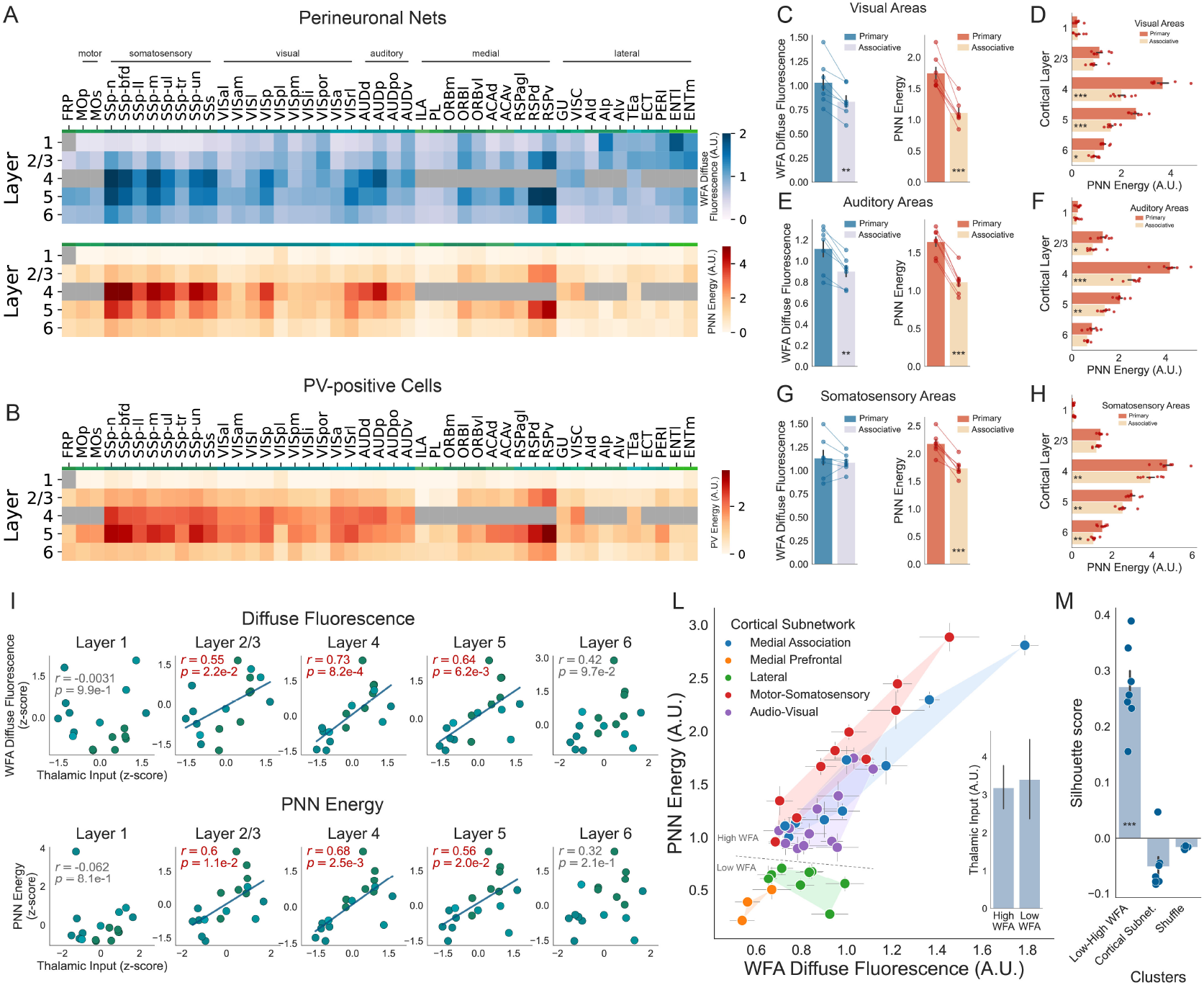
Organization of PNNs in cortical areas. (**A**) Heatmaps representing WFA diffuse fluorescence and PNN energy. Average metrics across mice are shown for each cortical area and layer (area acronyms are available in ***Table ST3***). In brain regions that do not have layer 4, the respective cells are grayed out. (**B**) Same as in (A) but for PV energy. (**C**) WFA diffuse fluorescence and PNN energy in the primary visual cortex versus higher-order associative visual areas. (**D**) PNN energy in primary versus associative visual cortical areas split by layer. (**E**) Same as in (C) but for auditory areas. (**F**) Same as in (D) but for auditory areas. (**G**) Same as in (C) but for somatosensory areas. (**H**) Same as in (D) but for somatosensory areas. (**I**) Correlation between WFA diffuse fluorescence and thalamic input strength in sensory-related areas of the cortex (all somatosensory, visual, and auditory cortices, see Methods & Materials) split by layer. In the bottom part, the same analysis was performed for PNN energy. Text insets indicate the Pearson correlation coefficient (r) and the corresponding p-value. For significant correlations, highlighted in red, the blue line shows the best linear fit. (**L**) Scatterplot of PNN energy vs WFA diffuse fluorescence for all cortical areas colored by their cortical subnetwork. The transparent shading represents the convex hull of all points in a subnetwork. Regions cluster into 2 groups: high-WFA and low-WFA. The inset shows the average thalamic input strength of regions divided into high- and low-WFA groups. (**M**) Silhouette score, representing a metric for clustering quality, calculated for each mouse by grouping cortical areas in: 2 groups (Low-High WFA), 5 groups (cortical Subnet.), or 2 groups but randomly shuffled (shuffle). In C, D, E, F, G, H, and M dots represent mice. In I and L dots represent brain areas. Error bars in C, D, E, F, G, H, L, and M represent SEM across mice. Error bars in L (inset) represent SEM across brain regions. See ***Table 1*** for statistical comparisons.

These results provide the first systematic and layer-specific description of PNNs in all cortical areas and indicate that layers 4-5 of primary cortical regions are privileged sites of PNN expression across multiple sensory systems.

### Determinants of cortical expression of PNNs: role of PV cells and area connectivity

We then investigated the factors responsible for the specific distribution of PNNs in the cerebral cortex. Considering the intimate relationship between PV cells and PNNs in the cortex (***Figure 3***G, ***Figure 4***D), one hypothesis could be that the high expression of PNNs in primary sensory cortices mirrors the distribution of PV cells. However, PV energy was only slightly increased in primary visual and auditory, but not somatosensory areas (***Figure 5***B, ***Figure S6***A, D, G). Accordingly, by splitting data by layers, we detected no differences between primary and associative regions for all the metrics with the exception of PV energy in deep layers of the visual cortex (***Figure S6***B, C, E, F, H, I). Intriguingly, we observed that PV cells in primary sensory cortices were more likely to have PNNs than in secondary areas (***Figure S7***A, C, E). This effect was not due to a higher proportion of high-PV cells in primary versus associative areas (***Figure S7***B, D, F), suggesting that the mechanism by which PNNs are increased in primary regions might be unrelated to PV expression levels.

The high levels of PNNs in layer 4 of primary sensory cortices could be related to the control of feed-forward sensory thalamic inputs that densely innervate layer 4 of primary sensory regions. Indeed, previous work showed that PNNs control plasticity of thalamic connections directly contacting PV cells of the primary visual cortex (Faini et al., 2018). If this hypothesis were true, one should expect PNN energy to scale with thalamic innervation density across sensory areas. To test this, we used published data from the mouse brain connectivity atlas of the Allen Institute (Oh et al., 2014) to measure thalamic input strength for all somatosensory, visual, and auditory areas (total inputs from the sensory-motor cortex related portion of the thalamus, DORsm as indicated in the CCFv3 nomenclature, see Correlation with thalamic afferent connectivity in Methods & Materials). Strikingly, we found that both WFA diffuse fluorescence and PNN energy in cortical layers 2/3, 4, and 5 were highly correlated with thalamic input strength, and this effect was most prominent in layer 4 where thalamic afferents could explain respectively 53% and 46% of the variance in the two PNN metrics (r=0.73 and 0.68) (***Figure 5***I). As a control, we performed the same analysis with connections originating from the associative cortex-related regions of the thalamus (DORpm) and we found no correlation with PNNs in any cortical layer (***Figure S8***).

This data corroborates the possibility that PNNs could be important for the regulation of sensorimotor thalamic inputs across multiple sensory modalities and may provide a basis to investigate the role of PNNs on feed-forward functional signaling in sensory cortices.

If connections represent a determinant factor for PNN abundance, it could be that groups of highly interconnected cortical regions have coregulated levels of PNNs. Recent work clustered the cerebral cortex in five distinct functional subnetworks (Kim et al., 2017; Zingg et al., 2014) based on their intracortical connections. We used this classification to explore whether PNNs were differentially expressed in these subnetworks. To test this hypothesis, we plotted PNN energy vs WFA diffuse fluorescence for each cortical region. We found that cortical subnetworks were clustered in two groups, with no overlap: a “low-WFA” group comprising the lateral and medial prefrontal subnetworks and a “high-WFA” group comprising audiovisual, motor-somatosensory, and medial association networks (***Figure 5***L). To quantify cluster separation, we grouped brain regions using three strategies: the high/low WFA as described above, the original five cortical subnetworks, and high/low WFA regions randomly shuffled. For each grouping, we calculated the silhouette score, a metric representing the separation and quality of data clustering (Zhao et al., 2018). We found that grouping cortical regions in high- and low-WFA resulted in the highest score (***Figure 5***M). The sub-division in high- and low-WFA region groups could not be explained simply by different thalamic input strength, since we did not observe any significant difference in the overall thalamocortical connectivity between these two groups of regions in the Allen Institute dataset (***Figure 5***L, inset). Conversely, we noticed that high-WFA areas also displayed increased PV energy and an increased proportion of high-PV cells (***Figure S9***), suggesting that the different PNN distribution across cortical subnetworks might be instructed by PV cells.

In summary, these results show that each cortical network displays a typical and homogenous PNN aggregation and that PV cells and PV expression level contribute to generating cortical PNN distribution.

### Gene expression correlates of PNNs

Finally, we asked whether PNN abundance could be correlated with gene expression patterns, possibly highlighting molecular principles underlying PNN organization and function. To answer this question, we analyzed data published in the Anatomic Gene Expression Atlas (AGEA) by the Allen Institute (Lein et al., 2007). This dataset describes region-specific expression levels for about 18,000 genes. For each gene, we correlated its expression in all the brain areas with a metric for PNN staining to detect genes whose pattern of expression is predictive of PNN presence. We found about 5,000 genes positively correlated, and about 1,000 negatively correlated with WFA (FDR<0.01, Benjamini-Hochberg, see also Correlation with gene expression and gene set overrepresentation analysis in Methods & Materials, and supplementary data SD4). It is important to note that this analysis reflects gene expression and PNNs at the level of brain areas and not single cells. To validate our approach, we selected a few genes known to be related to PNN structure and function: Aggrecan (*Acan*), a major proteoglycan core protein present in PNNs (Dauth et al., 2016; Fawcett et al., 2019; Ueno et al., 2018; Härtig et al., 2022; Oohashi et al., 2015; Yamada et al., 2017), Hyaluronan and proteoglycan link protein 1 (*Hapln1*), coding for a link protein essential for PNNs structure (Carulli et al., 2010); hyaluronan synthase 3 (*Has3*), a necessary component for PNN aggregation (Kwok et al., 2010); Matrix metallopeptidase 9 (*Mmp9*), an enzyme known to regulate PNN and PV development (Pirbhoy et al., 2020); A disintegrin and metalloproteinase with thrombospondin motifs (*Adamts5* also known as *Adamts11*), an aggrecan-degrading protease (Held-Feindt et al., 2006) that is expressed by PV interneurons with a PNN (Rossier et al., 2015), and parvalbumin (*Pvalb*). All these genes were significantly correlated with both PNN energy and WFA diffuse fluorescence (***Figure 6***A, B). Strikingly, out of 17639 genes, *Acan* was respectively the second and fifth most correlated gene with WFA diffuse fluorescence (*r_s_*=0.58) and PNN energy (*r_s_*=0.57). Consistently, when we repeated this analysis for PV energy we found that the top most correlated gene was *Pvalb* itself (*r_s_*=0.81). Other markers associated with PV neurons were also positively correlated (***Figure 6***C). These included the genes encoding the fast voltage-gated potassium channels Kv3.1 and Kv1.1 (*Kcnc1* and *Kcna1*) (Chow et al., 1999; Lorincz et al., 2008), and the sodium channel Nav1.1 (*Scn1A*) (Ogiwara et al., 2007)); synaptotagmin 2 (*Syt2*), a protein that ensures fast calcium sensing and vesicle release (Bouhours et al., 2017), and *Acan*. These results validated our approach, allowing us to provide lists of positive and negatively correlated genes that might highlight molecular regulators of PNNs. A detailed list of all 17,639 genes and their correlation with PNN and PV staining is available in supplementary data SD4.

**Figure 6.**
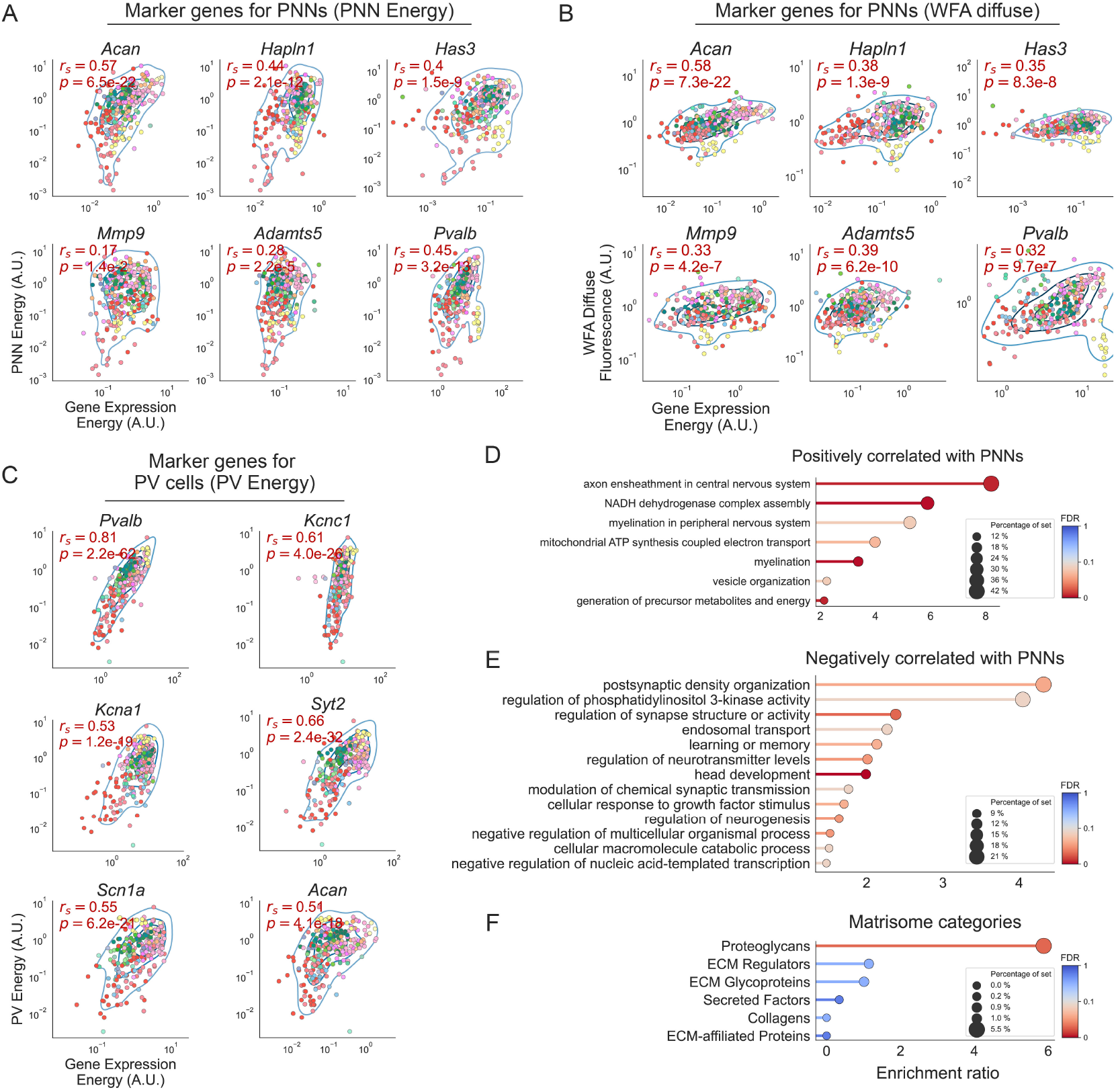
Gene expression correlates of PNNs. (**A**) Correlation between PNN energy and gene expression for six marker genes. *Acan* (aggrecan), *Hapln1* (link protein), *Has3* (Hyaluronan synthase 3), *Mmp9* (Matrix metalloprotease 9), *Adamts5* (an aggrecan-degrading protease), and *Pvalb* (parvalbumin). (**B**) Same as in A but for WFA diffuse fluorescence. (**C**) Correlation between PV energy and gene expression for six marker genes. *Pvalb* (parvalbumin), *Kcnc1* (potassium channel Kv3.1), *Kcna1* (potassium channel Kv1.1), *Syt2* (synaptotagmin 2), *Scn1a* (sodium channel Nav1.1), and *Acan* (aggrecan). (**D**) Biological process terms enriched in genes positively correlated with PNN energy. (**E**) Biological process terms enriched in genes negatively correlated with PNN energy. (**F**) Matrisome categories of genes positively correlated with PNN energy. In A, B, and C text insets indicate the Spearman correlation coefficient (rs), and the corresponding p-value and significant correlations are highlighted in red. Blue lines represent kernel density estimations. Data in D, E, and F are presented in descending order of enrichment ratio, colored based on the q-value with darker red shades corresponding to more significant values (threshold: FDR < 0.1). The dot size represents the percentage of genes of each category, that is present in the experimental gene list.

To obtain insight into the biological processes of the correlated genes, we performed a gene ontology analysis separately on the lists of the top 1,000 most correlated and anticorrelated genes with PNN energy, ranked by their correlation coefficient (***Figure 6***D, E). Genes related to processes of axon ensheathment, myelination, mitochondrial function, and cellular respiration were enriched in the pool of the positively correlated transcripts. Conversely, we found that anticorrelated genes were related to processes involved in synaptic plasticity, including among others, postsynaptic density organization, regulation of synapse structure, and learning and memory. This is consistent with the known inhibitory role of PNNs toward synaptic plasticity (Fawcett et al., 2019). Finally, we performed a similar overrepresentation analysis on a smaller gene set, the “matrisome” (Naba et al., 2016), containing about 1,000 genes related to different categories of ECM structure and function. Only the category proteoglycans was strongly overrepresented in the set of positively correlated genes (***Figure 6***F).

Taken together, these data show that we can reliably identify gene expression correlates of PNN abundance with the approach described above. Moreover, this analysis and the resulting gene lists could prove useful for designing experiments to investigate the molecular biology underlying PNN development and regulation.

## Discussion

In this study, we created and analyzed a whole-brain dataset of PNNs and PV cells in the adult mouse brain. We provide several quantitative measurements of the abundance of PNNs and PV cells and their colocalization in over 600 brain regions. The atlas was built using a shared spatial framework that facilitates replication studies and allows analyzing PNN data together with publicly available connectomics (Oh et al., 2014; Zingg et al., 2014) and gene expression (Lein et al., 2007) datasets, which enabled us to identify potential principles of PNN organization and gene expression profiles that are correlated or anticorrelated with PNN abundance. Previous studies have analyzed PNNs in multiple brain regions (Dauth et al., 2016; Ueno et al., 2018), however, they have been limited by their focus on only a subset of areas, their use of a more qualitative approach, or their use of a non-standard reference volume. In contrast, our atlas addresses all these aspects.

Our public resources (supplementary data SD1-4) will help researchers to generate novel hypotheses and questions, and to design experiments to better understand the function of PNNs and their involvement in pathological conditions.

### A toolset for PNN research: advantages and limitations

One of the challenges in studying PNNs is the difficulty of automatically detecting them due to their high morphological variability. To address this issue, we release two deep-learning models for the detection of PNNs and PV cells, pre-trained on about 0.8 million manually annotated PNNs/cells. The models can also be fine-tuned to specific experimental needs and image qualities with additional training. We have also made all of the raw and processed data from this study freely available (raw dataset: Zenodo link (Lupori et al., 2022), processed data: Supplementary data SD1-4).

In interpreting our results, it is important to note that we used WFA as a marker for PNNs. While WFA is a commonly used method for visualizing PNNs (Fawcett et al., 2019), it does not equally bind to all structures of aggregated CSPGs. Therefore, the use of other antibodies that specifically target different proteoglycans may be necessary to fully reveal the presence of these structures (Galtrey et al., 2008; Ueno et al., 2018; Ariza et al., 2018; Matthews et al., 2002). Our approach can be easily adapted to count these different types of PNNs, creating brain atlases of all the major components of PNNs using this method. Additionally, colocalization with other cell types could also be studied. For example, PV-positive neurons are a heterogeneous population (Tasic et al., 2016) that cannot be distinguished using our immunofluorescence approach. However, specific promoters and enhancers could be used to label PV-cell subtypes in a brain-wide manner, allowing the study of their colocalization with PNNs and a more detailed understanding of PNN expression regulation.

### Diffuse CSPGs and aggregated PNNs distributions

CSPGs are large, complex molecules that are widely distributed throughout the brain, whereas PNNs are aggregated around specific neurons (Fawcett et al., 2019). While most research on PNNs has focused on telencephalic and diencephalic structures, our analysis revealed that PNNs are highly abundant in the midbrain and hindbrain (pons and medulla) compared to other brain regions. These areas are important for vital processes such as heartbeat and breathing control, basic reflexes, motor control, and sleep (Ruder et al., 2021; Saladin et al., 2021). However, the role of PNNs in the neural circuits underlying these functions is largely unknown.

Another finding of our study is that CSPG aggregation in PNNs may be differentially regulated across brain areas. While in most of the brain the amount of non-aggregated CSPGs (as measured by diffuse WFA fluorescence) was a good predictor of the presence of aggregated PNNs (as measured by PNN energy), some areas showed no relationship between the two metrics. For example, all olfactory areas had very intense diffuse staining but contained very few and thin PNNs (***Figure 2A***, C, E)(Hunyadi et al., 2020), indicating that the high amount of CSPGs present in these areas did not aggregate into PNNs. This pattern was also observed in the cortical subplate (***Figure 2***E). The region-specific regulatory mechanisms of CSPG aggregation into PNNs and the functional implications are currently unknown and require further investigation.

### PV levels are associated with the presence of PNNs

A commonly observed property of cortical PNNs is that they preferentially aggregate around GABAergic PV-positive interneurons (Fawcett et al., 2019). We measured that, on average, this was the case for about 60% of PNNs in the entire brain, a much higher percentage than expected from chance. Moreover, across the whole brain, both PNN metrics were correlated with PV energy. Despite this clear association, our study unveils that slightly less than half of the PNNs in the brain do not surround PV neurons, leaving the still unanswered question of whether they might serve to regulate different circuit properties.

The link between PNNs and PV cells also varied between brain subdivisions with the most striking pattern in the isocortex. Here, 70% of all PNNs were around PV cells and half of all PV cells had a net. This intimate association was also evident in the relation between staining metrics. Indeed, cortical areas had a very tight (*r_s_*=0.91) correlation between PNN and PV energy.

Our analysis showed that the probability of being surrounded by a PNN for a PV cell is highly dependent on its PV expression level. Given that PV neurons differentiate before birth (Fishell, 2008) and PNNs aggregate much later during postnatal development (Reichelt et al., 2019), this association suggests that the developmental increase in PV expression enhances the probability to develop a PNN.

The magnitude of the association between PV levels and the probability of having a PNN, however, varies across brain structures suggesting that the mechanism that couples PV expression to PNN aggregation can be fine-tuned. For example, in the isocortex, hippocampal formation, and striatum, PV-PNN coupling was particularly strong. Intriguingly, in all three of these brain regions, PV cells have been previously divided, based on their intensity, into two distinct subpopulations of early-born high-PV cells and late-born low-PV cells with different roles in plasticity and learning (Donato et al., 2013; Donato et al., 2015). Our data are consistent with the interpretation that PNNs might aggregate more onto early-born high-PV neurons contributing to the inhibitory role of this subpopulation toward plasticity. In summary, it is currently unknown how perineuronal nets and parvalbumin are co-regulated. Previous evidence suggests that Otx2 may act as a mediator of this coupling, promoting the maturation of parvalbumin cells and PNNs (Gibel-Russo et al., 2022; Lee et al., 2017). This suggests that Otx2 may play a role in the co-regulation of these two factors, although further research is needed to confirm this hypothesis.

### PNN expression in the cortex is correlated with specific connectivity patterns

Our study demonstrated that strong PNNs are a common feature of layer 4 in all primary sensory cortices. This enrichment was evident also when we directly compared the labeling of primary and associative cortices within each sensory modality. Interestingly, this pattern cannot be explained solely by an increase in the number of PV cells or in the proportion of high-PV expressing cells that are more likely to have a PNN. At a functional level, the high expression of PNNs in primary sensory areas could be related to their action on thalamic afferents. Previous research in the mouse primary visual cortex showed that PNNs can selectively control thalamic excitation onto PV cells (Faini et al., 2018). Our data suggest that the control of feed-forward thalamo-cortical sensory inputs on PV neurons may be one important function across all sensory cortices. This is supported by the observation that the abundance of PNNs correlates with the density of thalamic innervation in all sensory areas. This hypothesis is also in accordance with the findings that plasticity in L4 of the visual cortex is lower (Trachtenberg et al., 2000) and might rely on a separate set of molecular mechanisms (Liu et al., 2008).

The relationship between thalamic inputs and PNN levels raises the possibility that the type of connections may be a determining factor in PNN expression. This idea was further supported by the observation that regions of the cortex with strong PV and PNN expression tend to have similar intracortical connectivity patterns (***Figure 5***L). This finding suggests that circuitry within these areas requires a certain level of stability, which could be achieved through the expression of PNNs. This novel concept merits further investigation to fully understand how this relationship functions.

### Gene expression correlates of PNNs

The search for a gene expression signature of PNN-enwrapped cells is hampered by the fact that PNNs are extracellular multimolecular structures, and that there is currently no means to tag the PNN-positive neurons.

To overcome this problem, we performed a correlational analysis between the AGEA dataset by the Allen Institute (Lein et al., 2007) and PNN expression. This novel approach was validated by the overrepresentation analysis on the matrisome gene set, which showed that PNN-correlated genes are strongly enriched in the proteoglycan category, and by finding key constituents of the PNN ranking in the top positions of the list of genes positively correlated with PNN energy. However, this approach also revealed many other genes with positive and negative correlations with PNNs. A gene ontology analysis strikingly showed that categories related to synaptic function and synaptic plasticity were significantly downregulated in brain areas enriched with PNNs. Furthermore, PNNs were found to be correlated with genes involved in myelination, another plasticity brake (Boghdadi et al., 2018; Bonetto et al., 2021), and genes related to cell metabolism, which may be due to the high energy demands of fast-spiking PV cells (Carter et al., 2009; Kann et al., 2014).

These results not only support the hypothesis that PNNs serve as plasticity brakes in the visual cortex (Fawcett et al., 2019), but also demonstrate that this functional signature emerges from an unbiased comprehensive analysis of all brain regions.

Our work represents a unique approach based on a brain-wide comparison of very large datasets of cellular structures with public resources. This type of analysis has the advantage of being unbiased and data-driven, which is typical of -omics techniques. It can also be applied to the study of many other extracellular matrix components. We envision that the advent of spatial transcriptomics will further enhance this type of approach.

## Methods & Materials

### Mice Handling

All experiments were carried out in accordance with the European Directives (2010/63/EU), and were approved by the Italian Ministry of Health (authorization number 723 / 2020 PR). A total of 7 adult C57BL/6J male and female mice, at approximately postnatal day (P)150 were used in this study. Weaning was performed at P21–23. Animals were maintained at 22°C with a standard 12-h light-dark cycle. During the light phase, a constant illumination below 40 lux from fluorescent lamps was provided. Mice were housed in conventional cages (365 x 207 x 140 mm, 2-3 animals per cage) with nesting material, and had access to food and water ad libitum. During the first 12-14 weeks of life, mice were fed a standard diet (standard diet Mucedola 4RF25). Then, animals were fed a balanced purified diet (Research Diets, Inc., New Brunswick, NJ, USA, cat. no. D12450Ji) for 6 weeks before the sacrifice.

### Immunofluorescence staining

Mice were anesthetized with chloral hydrate (20 ml/Kg BW) and perfused via intracardiac infusion with cold PBS and then 4% paraformaldehyde (PFA, w/vol, dissolved in 0.1M phosphate buffer, pH 7.4). Brains were extracted and post-fixed overnight in PFA 4% at 4°C, then transferred to a 30% (w/vol) sucrose solution for 48 hours. For each brain, 50 μm coronal sections, spanning from the anteriormost part of the cerebral cortex to the cerebellum, were cut on a freezing microtome (Leica). One out of every 3 sections was collected for further processing, leading to a sampling of one slice every 150μm. For a small subset of sections that did not match our quality standards due to deformations during the cutting process (on average 3.7±0.5 slices per animal), an adjacent section was collected instead. For each animal, slices were assigned a unique ID and pooled in 9-10 wells of a 24-well plate for free-floating staining. Each well contained 5-6 sections that sampled the brain at equally spaced points in the anterior-posterior axis.

Slices were blocked for 2h at room temperature (RT) in a solution containing 3% bovine serum albumin (BSA,A7906 Sigma-Aldrich) in PBS. Then, slices were incubated overnight at 4°C with a solution containing biotinylated Wisteria floribunda Lectin (WFA, B-1355-2, Vector Laboratories, 1:200) and 3% BSA in PBS. On the following day, sections were rinsed 3 times in PBS (10 min each) at RT, incubated with a solution of red fluorescent streptavidin (Streptavidin, Alexa Fluor™ 555 conjugate, S21381, Thermo Fisher, 1:400) and 3% BSA in PBS for 2h at RT, and rinsed again 3 times in PBS. On the same day, slices were incubated with a blocking solution for parvalbumin staining containing 10% BSA and 0.3% Triton in PBS for 30 minutes, then washed 3 times (10min each) and finally incubated overnight at 4°C with primary antibody solution containing anti-parvalbumin (Parvalbumin antibody, 195004, Synaptic System 1:1000) 1% BSA and 0.1% Triton in PBS. Then, sections were rinsed 3 times (10 min each) in PBS; incubated with a secondary antibody solution containing secondary antibody (anti-Guinea Pig IgG Alexa Fluor™ 488, A11073, Invitrogen, 1:500), 1% BSA. plus 0.1% Triton for 2h at RT, and washed again 3 times in PBS. Finally, sections were mounted on microscopy slides with a mounting medium (VECTASHIELD^®^ antifade mounting medium, H-100, Vector Laboratories), and stored at 4°C. All sections in each staining well were mounted on the same slide.

### Image acquisition

All images were acquired using the acquisition software ZEN blue with a Zeiss Apotome.2 microscope and a 10x objective and digitized by an AxioCam MR R3 12-bit camera, resulting in a pixel size of 0.645μm. For the WFA channel, excitation light passed through a 538-562nm bandpass filter and a 570nm dichroic mirror, while emitted light was filtered with a 570-640nm bandpass filter. For the PV channel, filters were a 450-490 nm bandpass for excitation, a 495nm dichroic mirror, and a 500-550nm bandpass for emission. All images were acquired with the same intensity of excitation light and with an exposure time of 80ms for the WFA channel and 850ms for the PV channel. For all sections, we acquired 3 apotome images for optical sectioning. Each brain slice was acquired as a tiled multi-image experiment on a single z-plane.

Coronal sections of the entire mouse brain span a relatively large area and even small irregularities in the microscope slide can lead to artifacts in image intensity due to the tissue section not sitting exactly perpendicular to the optical path. To account for this, we acquired each slice with a tilted z-plane linearly interpolated between 4 manually selected focus points at the edges of each section. After the acquisition, multi-image tiles were stitched in ZEN and exported as 8-bit TIFF files for further processing. The resulting dataset consisted of 842 single channel, 8-bit, TIFF images ranging from 7 to 165MB in size and from 2646 to 17631px (width) in resolution.

### Image registration to the Allen Brain Atlas CCF v3

#### Image Preprocessing

For each mouse, all the images were ordered along the anterior-posterior axis according to their unique ID. Images were manually inspected and, based on irregularities in the fixed brain and anatomical landmarks, a minority of them were mirrored vertically to make sure matching hemispheres were always on the same side for the whole image sequence.

All the following steps of preprocessing and image registration were carried out on a down-sampled (20% of the original size) TIFF dataset. For each downsampled experimental image, we created a matching binary mask of the same size, encoding whether each pixel belongs to brain tissue or not. Masks were automatically generated for the entire subsampled dataset by using a machine learning model (random decision forest) interactively trained with Ilastik (Berg et al., 2019) on a subset of 57 image crops (width ranging from 344px to 526px). Masks were used in the quantification steps to restrict fluorescence analysis only to portions of the images that contained biological tissue. All the masks were visually inspected through a custom MATLAB graphical user interface (GUI) and, if necessary, manually adjusted to correct for misclassification of small areas or to exclude parts of the tissue containing experimental artifacts from further analysis.

#### Image Registration

We aligned our dataset to the Allen Mouse Brain Common Coordinate Framework (CCFv3) (Wang et al., 2020) with a multi-step workflow: first, we used the software [QuickNII](https://www.nitrc.org/projects/quicknii) v2.2 (Puchades et al., 2019) to interactively assign each experimental image to a specific plane in the reference atlas based on anatomical landmarks. The software allows the selection of an arbitrary 2D plane out of the CCFv3 volume, thus improving accuracy for samples where sections were not cut on a perfectly coronal plane, but with a slight angle. In the same software, we also performed rigid transformations (i.e., rotations and translations) and uniform horizontal or vertical stretch in order to match the reference plane to each experimental image. In a second step, we used the software VisuAlign v0.9 (RRID: SCR_017978, [VisuAlign](https://www.nitrc.org/projects/visualign/)) to manually apply local, non-rigid transformations to the planes selected in QuickNII in order to match the experimental images.

We then used a custom set of MATLAB functions to load the output file from VisuAlign and to generate a displacement field for each experimental image. Each displacement field defines the local non-rigid transformation as a couple of values (*D_x_, D_y_*) for each pixel, defining the displacement in the image on the X and Y axes. By using the coordinates of the 2D plane defined in QuickNII and the local transformations defined in the displacement field it is possible to match each pixel position in our experimental images (*X_e_, Y_e_*) to a voxel position in the reference atlas (*X_a_, Y_a_, Z_a_*).

### Deep learning models for cell counting

The deep learning models used in this work are based on a novel counting strategy described in Ciampi et al., 2022 specifically designed to account for the variability between experimenters when counting non-trivial, overlapping, or low-contrast objects like PNNs in histological preparations. Briefly, cell counting for both PNNs and PV cells was done through a two-step pipeline. In the first step, we performed cell detection by using the Faster-RCNN network (Ren et al., 2015) with a Feature Pyramid Network module and a ResNet-50 backbone. The goal of this stage is to produce a collection of putative object locations with high recall. The training dataset of this network is large but labeled by a single rater, thus it is assumed to be “weakly labeled”, i.e., it may contain spurious (false positives) and missing annotations (false negatives). In the second step, we scored each detected object to assign it an “objectness” value designed to maximize its correlation with the raters’ agreement. To do this, we trained a small convolutional network to rank samples with increasing agreement values and produce an increasing score for objects with increasing raters’ agreement (***Figure S1***B, C). In this stage, we employed a smaller training dataset labeled by multiple raters for which the agreement between experimenters on each object was computed (see Training Dataset below).

Following this strategy, we employed four different models: a localization model for PNNs and PV cells, and a scoring model for PNNs and PV cells. From now on, we will refer to these models respectively as PNN_*loc*_, PV_*loc*_, PNN_*score*_, and PV_*score*_. We first localized and scored PNNs using PNN_*loc*_ and PNN_*score*_ and PV cells using PV_*loc*_ and PV_*score*_ on separate image channels. Then, we removed PNNs with a score lower than 0.4 and PV cells with a score lower than 0.55.

As a performance metric for this counting pipeline, we computed the mean absolute relative error (MARE) as follows:

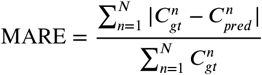

where *N* is the number of test images, and 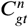 and 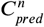 are the ground-truth and the predicted count of the n-th image, respectively. On the test split of our multi-rater dataset, our counting approach achieves a MARE of 0.048 and 0.080 respectively for PNNs and PV cells when considering samples located by at least 3 raters. As a final quality check, we visually inspected all the images and manually removed cases of artefactual cell detection. The source code for training models or making predictions with a pre-trained model can be found at this [link](https://github.com/ciampluca/counting_perineuronal_nets).

#### Training Dataset

Here we describe the training dataset used for each model.

The dataset used for the PNN_*loc*_ model consists of 580 8-bit grayscale TIFF images (width ranging from 2646 to 17631px) dot-annotated with the (x,y) position of each PNN for a total of 678556 PNNs. The dataset used for the PV_*loc*_ model consists of 53 8-bit grayscale TIFF images (width ranging from 5157 to 16389px) dot-annotated with the (x,y) position of each cell for a total of 101348 PV cells. PNNs were annotated by looking for distinctive circular patterns of WFA staining around cell somata and proximal dendrites. Finer PNN-like structures exclusively present in the neuropil, like those found in the olfactory bulbs (Hunyadi et al., 2020), were not annotated in our training dataset due to the magnification factor in our images not allowing for consistent detection of such structures.

The datasets used for the two scoring models both consist of a collection of 25 8-bit grayscale TIFF images (2000 x 2000 px). Seven expert experimenters independently dot-annotated each image for a total of 4727 PNNs and 5833 PV cells that vary in the agreement between raters from 1/7 to 7/7. Pre-trained models, ready for making predictions on new images, are available at this [link](https://github.com/ciampluca/counting_perineuronal_nets/releases).

### Brain structure sets

Throughout the paperwe aggregated data in three sets of brain structures differing by their level of spatial resolution or granularity. The first structure set (`structure_set_id:` 687527670) has a low level of resolution and is composed of 12 coarse-ontology major brain divisions (see ***Table ST2***). The second structure set (`structure_set_id:` 167587189) has a medium level of resolution (e.g., it comprises distinct cortical areas) and is composed of 316 mid-ontology brain regions (see ***Table ST4***).

These two structure sets were defined by the Allen Institute in their [API](https://allensdk.readthedocs.io/en/latest/install.html) and can be accessed using the `StructureTree` object. Lastly, for the analysis of cortical layers, we maintained the finest level of resolution present in the CCFv3, where individual cortical layers are segmented (see ***Table ST3*** for the definition of cortical areas). Please note that, for the visualizations in Fig. 5A-B, we included the lateral and medial parts of the entorhinal cortex (ENTl and ENTm, that actually belong to the hippocampal formation) given their layered structure. For all the analyses in the paper, we dropped data of any structure belonging to, or descending from the fiber tracts (`areaID`:1009) and the ventricular system (`areaID`:73).

### Data analysis

All data analysis was done using custom software written in MATLAB 2021b and Python (3.8). We used the following additional Python libraries for data analysis: NumPy (1.23.5) (Harris et al., 2020), Pandas (1.5.2) (McKinney, 2010), Scikit-learn (1.1.3) (Pedregosa et al., 2011) and SciPy (1.9.3) (Virtanen et al., 2020).

#### Measurement of single-cell staining intensity

Quantification of the staining intensity of individual cells (PNNs or PV cells) was performed on 80×80 pixels image tiles centered on the (x,y) center positions of each PNN/cell. Within each tile, we segmented pixels belonging to the cell or the background, and the intensity of each PNN/cell was defined as the average value of the pixels belonging to that cell. The segmentation was performed by using a random forest pixel classifier implemented with the MATLAB `Treebagger` class with the support of additional custom MATLAB functions (Cicconet et al., 2019). This approach allows the classification of single pixels as background or foreground, based on a collection of features of that pixel. Classifying all the pixels in an image tile results in a binary segmentation mask.

The features considered for pixel classification were the contrast-adjusted pixel intensity (using the `imadjust` MATLAB function), the position of the pixel relative to the center of the tile in the horizontal and vertical axes, and the pixel intensity in 16 versions of the image tile filtered with 16 Gabor filters. The wavelength and orientation of each Gabor represented one of the possible combinations of four different wavelength values (2.8, 5.6, 11.3, 22.6 pixels/cycle) and four different orientations (0°, 45°, 90°, 135°). Wavelengths were sampled in increasing powers of 2 starting from 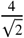 up to the hypotenuse length of the input image tile, while orientations were sampled from 0° to 135° with a step of 45° (Jaini et al., 1991). Each random forest model for segmentation of PNNs and PV cells was trained on 69600 pixels from 1160 tiles (60 pixels randomly chosen for each tile).

#### Staining metrics definitions

We defined four metrics to quantitatively analyze the staining for PNNs and PV cells.

First, *diffuse fluorescence* represents the amount of average fluorescence signal in a brain region. It is defined as the average intensity of all the pixels belonging to that region across all the slices of each mouse. These values were then normalized within each mouse by dividing them by the mean pixel intensity of all the brain. This normalization removes global differences in intensity between mice (due to for example perfusion quality and post-fixation) while highlighting how staining intensity is differentially distributed across brain regions. As a result, a region with diffuse fluorescence of 1 would have a staining intensity equal to the brain average.

Second, *density* represents the number of cells or PNNs per unit of area in a brain region. It was defined as the total number of cells or PNNs belonging to that region across all the slices of each mouse, divided by the total area belonging to that region in mm^2^.

Third, cell *intensity* represents the staining intensity of cells or PNNs in a brain region. Each cell was assigned a value of staining intensity(see section Measurement of single-cell staining intensity). For each region, cell intensity was defined as the average intensity of all the cells belonging to that region. These values were then normalized to the range 0-1 by dividing by 255 (maximum intensity value for 8-bit images).

Last, we defined a combined, more abstract metric, that takes into account both the number and the intensity of cells/PNNs, called *energy*. Cell energy can be thought of as a measure of cell density, weighted on intensity. For each region, energy is defined as the sum of the cell intensity of all the cells in that region, divided by the total surface area. For a region of area *A*, containing *c* cells:

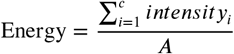

These values were then normalized within each mouse by dividing them by energy calculated on the entire brain. As a result, a region with an energy value of 1 would be equal to the brain’s average energy. This definition of energy is analogous to the one used by the Allen Institute in (Lein et al., 2007) for the analysis of in-situ hybridization data (see the technical paper on the informatics data processing [here](http://help.brain-map.org/display/mousebrain/Documentation)). It is important to note that the brain of each mouse in this study has been sampled in its entire anterior-posterior axis with the same sampling rate (1 every 3 slices) thus ensuring that the normalization step for diffuse fluorescence and energy measurements does not introduce biases due to differential sampling of areas with extreme staining intensity values.

#### Colocalization PNN-PV

PV cells and PNNs were counted with two distinct deep learning models on separate channels. We defined a PNN and a PV cell to be colocalized based on their (x,y) position in the original image using the following criteria. We selected one cell/PNN at a time as a reference object. For each reference object, we selected only objects in the other channel with a distance equal to or smaller than 15 pixels (9.675μm). If multiple objects satisfied this criterion, we picked the closest one as a colocalized object. Otherwise, if no objects were close enough to the reference one, we defined the reference object as non-colocalized (either a PV-negative PNN or a WFA-negative PV cell).

We computed two metrics to describe PNNs and PV colocalization: first, the percentage of PV+ PNNs, that is the fraction of PNNs that are around a PV-positive cell; second, the percentage of WFA+ PV cells, that is the fraction of PV-positive cells that are surrounded by a net. Colocalization metrics at the coarse level of resolution (see section Brain structure sets for definition) were calculated independently for each mouse and the results averaged across mice. For the same analysis at higher levels of resolution (mid-ontology in ***Figure 3***C and ***Figure S4***), we adopted a different strategy. At higher resolutions, brain subdivisions are much smaller and some areas contain a limited number, or even no, of PNN or PV-cells (e.g., layer 1 of cortical areas). As a result, the percentage of colocalization can vary dramatically depending on a few, or even a single cell, thus not providing a robust measure for that area (e.g., an area with 3 PV cells can vary from 0% to 100% depending on the state of PNNs on only 3 neurons). To solve this issue, we calculated colocalization metrics on a dataset of cells pooled from all animals except one, in a manner similar to the leave-one-out cross-validation approach used in machine learning (Wong, 2015). We repeated this process for all mice and considered each repetition an “experimental unit”. We then averaged across experimental units. For the analysis of the colocalization of PNNs and PV cells (***Figure 3***C and ***Figure S4***) we included only brain regions that contained at least 3 PNN and 3 PV cells in at least 4 mice.

#### PV intensity classes

PV cells were divided into four intensity classes of equal width based on their cell intensity levels. The classes were defined as 1: low PV (PV intensity in the range [0, 0.25)); 2: intermediate-low PV (range [0.25, 0.5)); 3: intermediate-high PV (range [0.5, 0.75)); 4: high PV (range [0.75, 1]). The probability of being surrounded by a net was estimated by dividing the total number of PV cells in that class by the number of colocalized PV-PNN cells. This analysis was done independently for each mouse. We fit data to a first-degree linear equation by using the numpy function `np.polyfit`. The estimated first- and zero-order parameters are displayed in the text insets for each plot.

#### Correlation between staining metrics

The analysis of correlations between staining metrics in all the figures (***Figure 2***E, ***Figure 3***D-G, ***Figure S3***E) was done by computing the Spearman’s rank correlation coefficient using the SciPy function `stats.spearmanr`. In each graph, we reported the value of the correlation coefficient (*r_s_*) and the associated p-value. We highlighted in red significant (p <0.05) correlations. For significantly correlated metrics we also reported in blue a linear fit obtained using a Huber regressor robust to outliers (Huber et al., 2009) using the implementation in `sklearn.linear_model.HuberRegressor`.

#### Correlation with thalamic afferent connectivity

To measure thalamic input strength we used connectomics data from the Allen Institute (Oh et al., 2014). In that dataset, we selected the connections that originated from the thalamus and that terminated in sensory-related cortical regions (SSp-n, SSp-bfd, SSp-ll, SSp-m, SSp-ul, SSp-tr, SSp-un, SSs, VISal, VISam, VISl, VISp, VISpl, VISpm, VISli, VISpor, AUDd, AUDp, AUDpo, AUDv). For ***Figure 5***I we selected only thalamic inputs originating from the sensory-motor cortex related part of the thalamus (DORsm, area ID: 864, according to the CCFv3 nomenclature, https://atlas.brain-map.org/). For ***Figure S8*** we selected only thalamic inputs originating from the polymodal-association cortex related part of the thalamus (DORpm, `areaID`: 856). Input strength for each cortical area was measured as the sum of connection strength from all brain regions belonging to either the DORsm or the DORpm to both the ipsilateral and contralateral parts of that cortex. To uniform the scale of PNN measurements and thalamic connectivity, we z-scored each set of data. For the correlation analysis (***Figure 5***I), we computed Pearson’s correlation coefficient and the associated p-values. To estimate connection strength in high-WFA and low-WFA region clusters (***Figure 5***L inset), we averaged thalamic input strength values, obtained in the same way, of all the areas in each cluster.

#### Correlation with gene expression and gene set overrepresentation analysis

We correlated the distribution of PNN energy, WFA diffuse fluorescence and PV energy with the pattern of expression of approximately 18,000 genes, published in the Anatomic Gene Expression Atlas (AGEA) by the Allen Institute (Lein et al., 2007). In this dataset, levels of expression of each gene are derived from the signal intensity of whole-brain in situ hybridization essays and quantified as expression energy, a metric defined in an analogous way to PNN and PV energy. For correlation analysis, both gene expression data and PNN or PV staining parameters were expressed at mid-ontology resolution (see ***Table ST4***). The five areas showing the largest standard deviation in PNN or PV staining metrics were excluded from the analysis. We computed Spearman’s rank correlation coefficient between each of the 3 staining metrics and the pattern of expression of each of the AGEA genes. Correction for multiple testing was performed with Benjamini-Hochberg method. For all the analyses, we considered genes with a q-value<0.01 (Benjamini-Hochberg method) as significantly correlated (if Spearman’s correlation coefficient was positive) or anticorrelated (if Spearman’s correlation coefficient was negative) with the staining metric considered.

For the genes correlated and anticorrelated with PNN energy and WFA fluorescence, we performed gene ontology analysis using WebGestalt platform (Zhang et al., 2005). Overrepresentation of gene ontology terms (biological process domain) was tested separately for the 1,000 genes most correlated (with the largest correlation coefficient) and the 1,000 genes most anticorrelated (with the most negative correlation coefficient) with each of the two metrics. The list of all the genes present in the AGEA was used as the background for all the analyses. Overrepresented gene ontology terms were filtered to ensure a false discovery rate<0.1 (Benjamini-Hochberg method) and clustered via affinity propagation to reduce redundancy.

We then tested for overrepresentation of gene sets related to ECM biology, defined by (Naba et al., 2016) as matrisome categories, in the 200 genes most correlated with PNN energy. As for gene ontology, the entire list of genes of the AGEA was used as the background. To assess statistical significance, we performed hypergeometric test and corrected for multiple testing using Benjamini-Hochberg method. For each matrisome category, the enrichment ratio was calculated as the number of genes observed in both the matrisome category and the 200-gene list divided by the number of genes expected assuming independence of the matrisome set and the gene list.

## Data availability

We strongly and actively support the open availability of all experimental data and analysis code in this manuscript. In this spirit, we made them accessible in the following repositories:

- Raw microscopy [dataset](https://doi.org/10.5281/zenodo.7419283)
- Analysis [code](https://github.com/LeonardoLupori/wholeBrain_PNN_analysis) to reproduce figures
- Deep learning model [code](https://github.com/ciampluca/counting_perineuronal_nets)

## Funding

This work was funded by: AI4Media - A European Excellence Centre for Media, Society and Democracy (EC, H2020 n. 951911); the Tuscany Health Ecosystem (THE) Project (CUP I53C22000780001), funded by the National Recovery and Resilience Plan (NRPP), within the NextGeneration Europe (NGEU) Program;PRIN2017 2017HMH8FAto T.P., R.M. was supported by Fondazione Umberto Veronesi.

## Competing interests

The author declare no competing interests.

## Data visualization

Data visualization for all the figures was done in Python (3.8). Heatmaps, bar plots, and scatterplots were created using the libraries Seaborn (0.12.1) (Waskom, 2021) and Matplotlib (3.4.2) (Hunter, 2007). Rendered heatmaps of coronal brain slices were done by using BrainRender (Claudi et al., 2021) and bg-heatmaps (Claudi et al., 2022).

## Acknowledgment

This work was funded by: AI4Media - A European Excellence Centre for Media, Society and Democracy (EC, H2020 n. 951911); the Tuscany Health Ecosystem (THE) Project (CUP I53C22000780001), funded by the National Recovery and Resilience Plan (NRPP), within the NextGeneration Europe (NGEU) Program; PRIN2017 2017HMH8FA to T.P., R.M. was supported by Fondazione Umberto Veronesi.

We thank Silvia Burchielli, Cecilia Ciampi, Domiziana Terlizzi, and Sara Ciampi, for their support in raising animals in the animal facility.

We also thank our colleagues, Cristiano Ricci, Francesco Calugi, Giulia Sagona, Elsa Ghirardini, Laura Baroncelli, Francesca Damiani, Matteo Alberti, Andrea Tognozzi, Mariagrazia Giuliano, and Matteo Caldarelli for scientific discussions.

## Author contributions

Contributions of each author, based on the CRediT taxonomy (Brand et al., 2015) are shown in ***Figure 7***.

**Figure 7.**
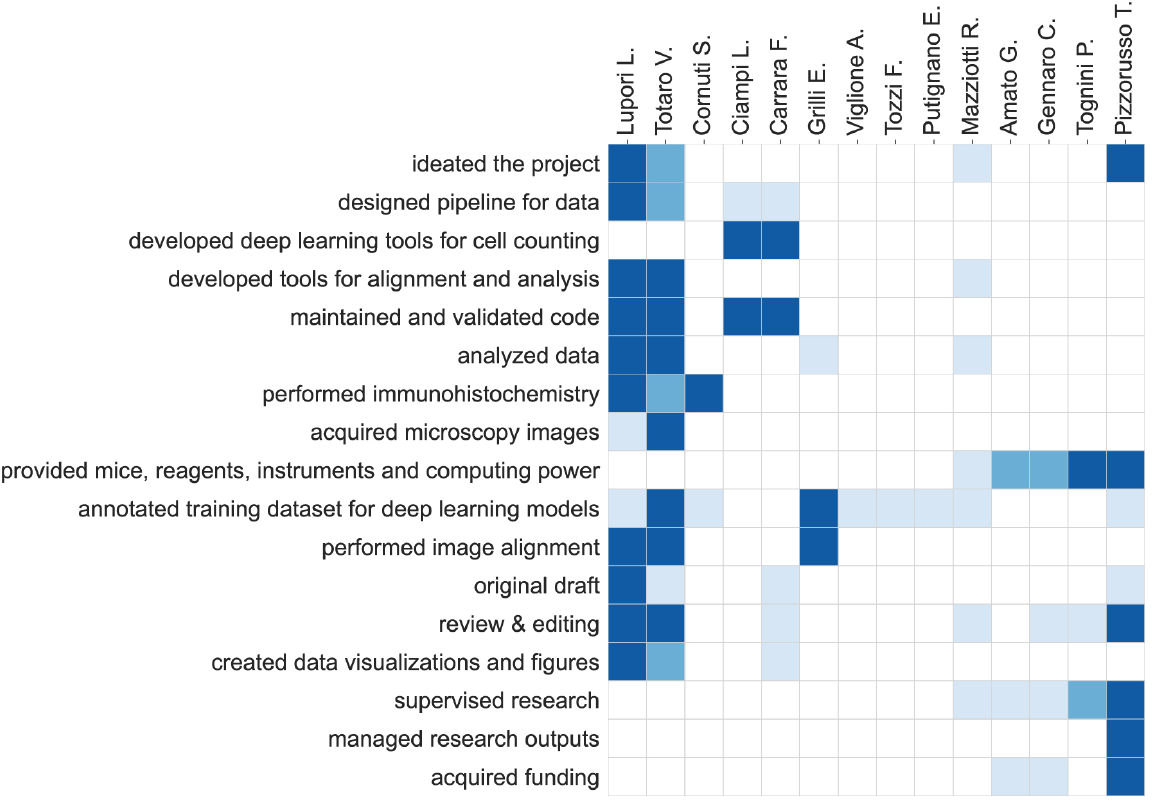
Author contributions. For each type of contribution, there are three levels indicated by color in the diagram: ‘support’ (light), ‘equal’ (medium), and ‘lead’ (dark).

## Supplementary Material

### Supplementary Figures

**Figure S1.**
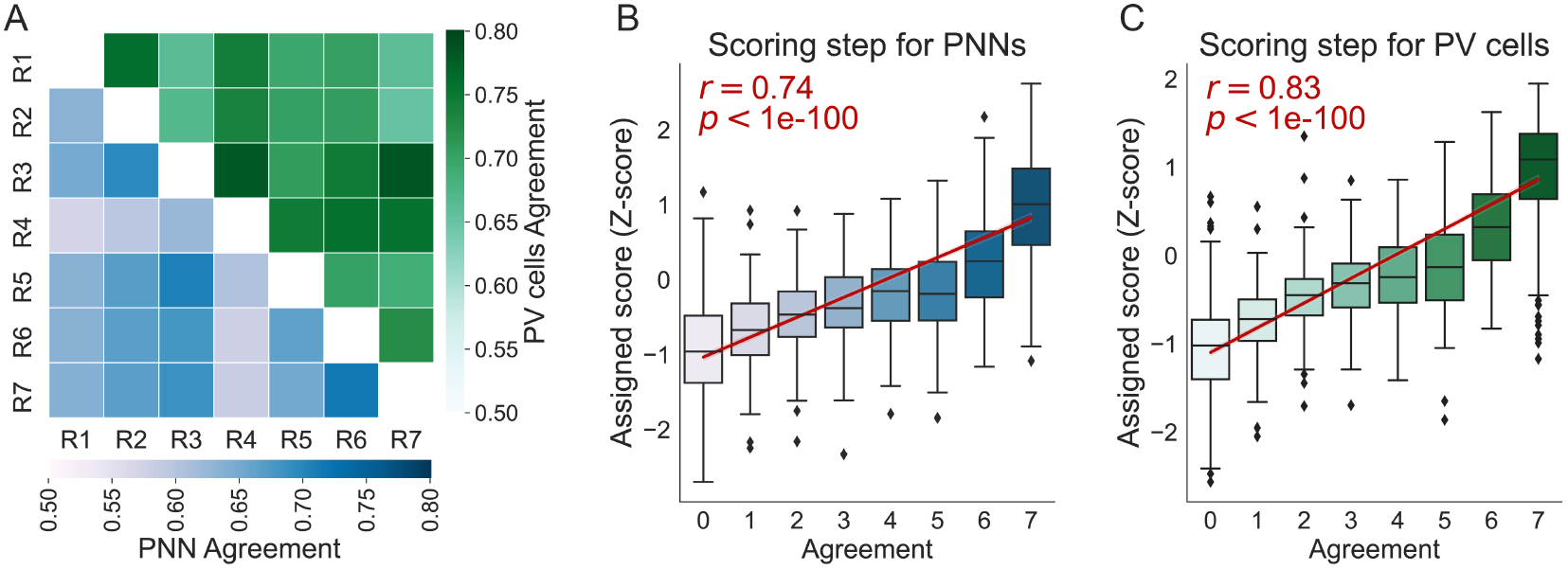
The scores assigned by the scoring models correlate with raters’ agreement. (**A**) Agreement (Jaccard index) between the manual cell annotations of 7 independent raters (R1-R7). The lower part of the matrix (blue shade) represents agreement in PNN counts, while the upper part (green shade) represents agreement in PV counts. (**B**) Performance of the scorer module for PNNs. Individual PNNs are grouped according to their agreement level in the multi-rater dataset and the score assigned to them by the scorer module is shown on the Y-axis. (**C**) Performance of the scorer module for PV cells. Individual PV cells are grouped according to their agreement level in the multi-rater dataset and the score assigned to them by the scorer module is shown on the Y-axis. In B and C, text insets represent Pearson’s correlation coefficient (*r*) and the corresponding p-value. Boxes represent quartiles, whiskers extend to 1.5 IQRs of the lower and upper quartile, and observations that fall outside this range are displayed independently.

**Figure S2.**
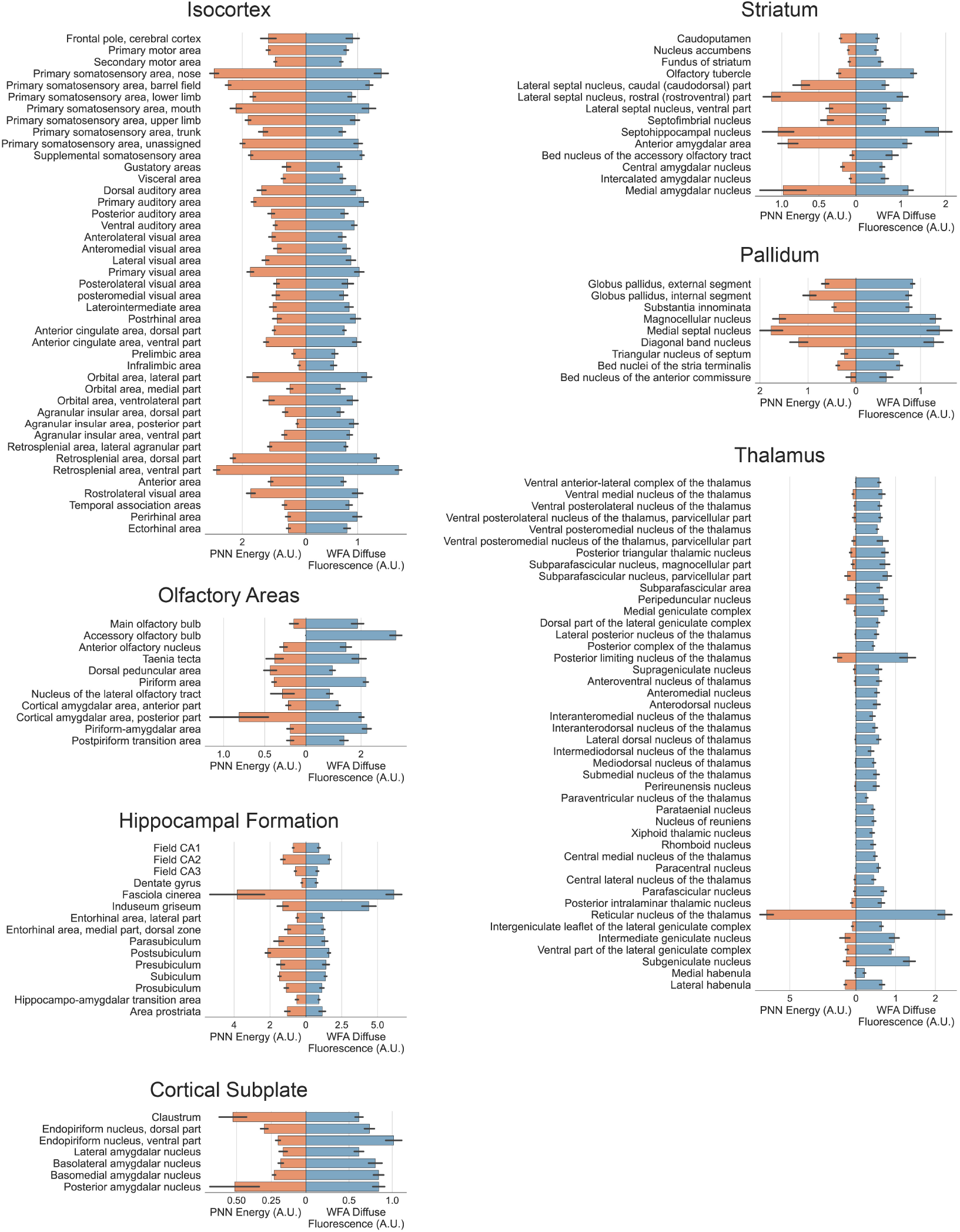
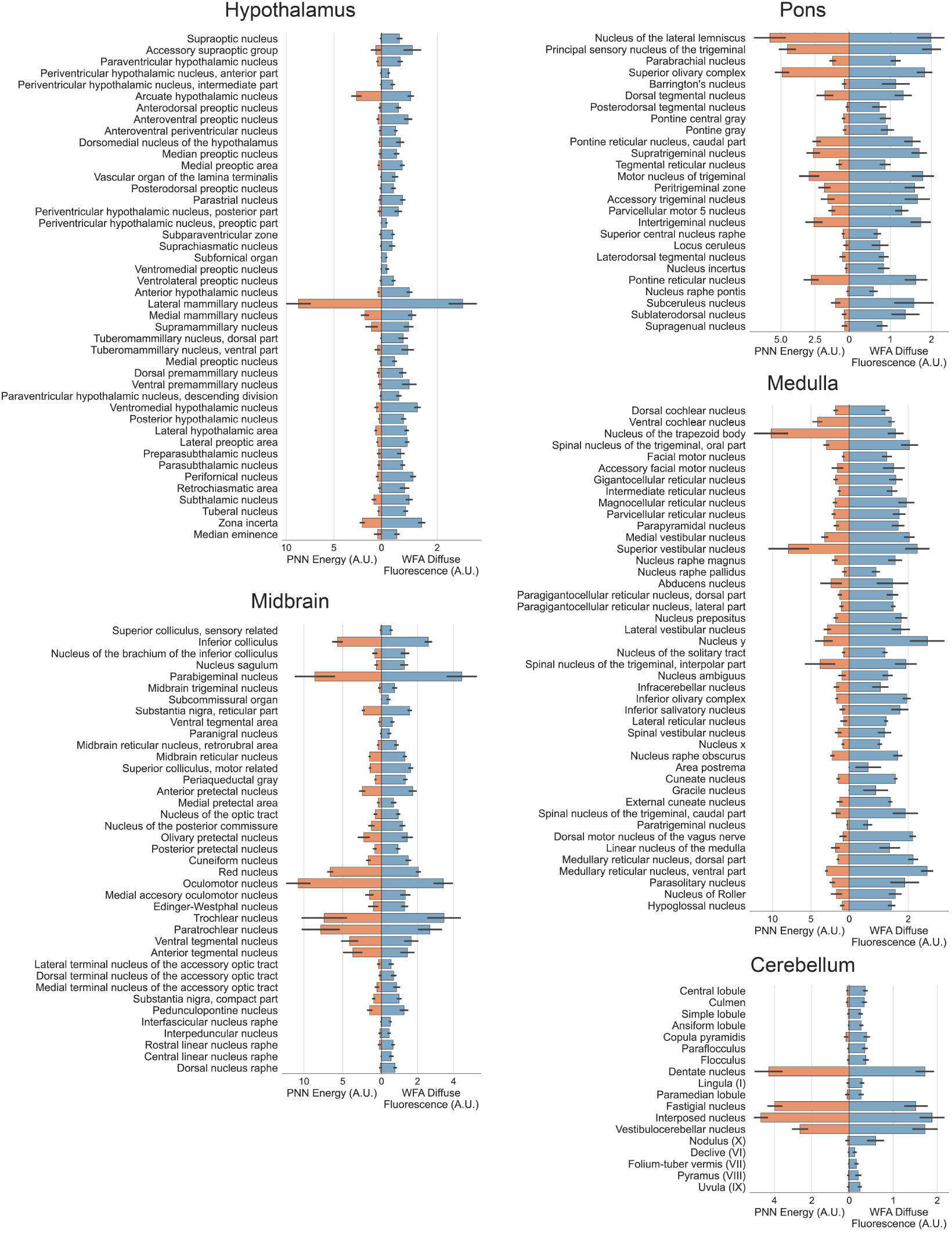
PNN energy and WFA diffuse fluorescence measurements for medium-resolution brain areas grouped by their major subdivision. For each plot, on the left in orange is represented PNN energy, while on the right in blue is represented WFA diffuse fluorescence. Error bars represent SEM across mice.

**Figure S3.**
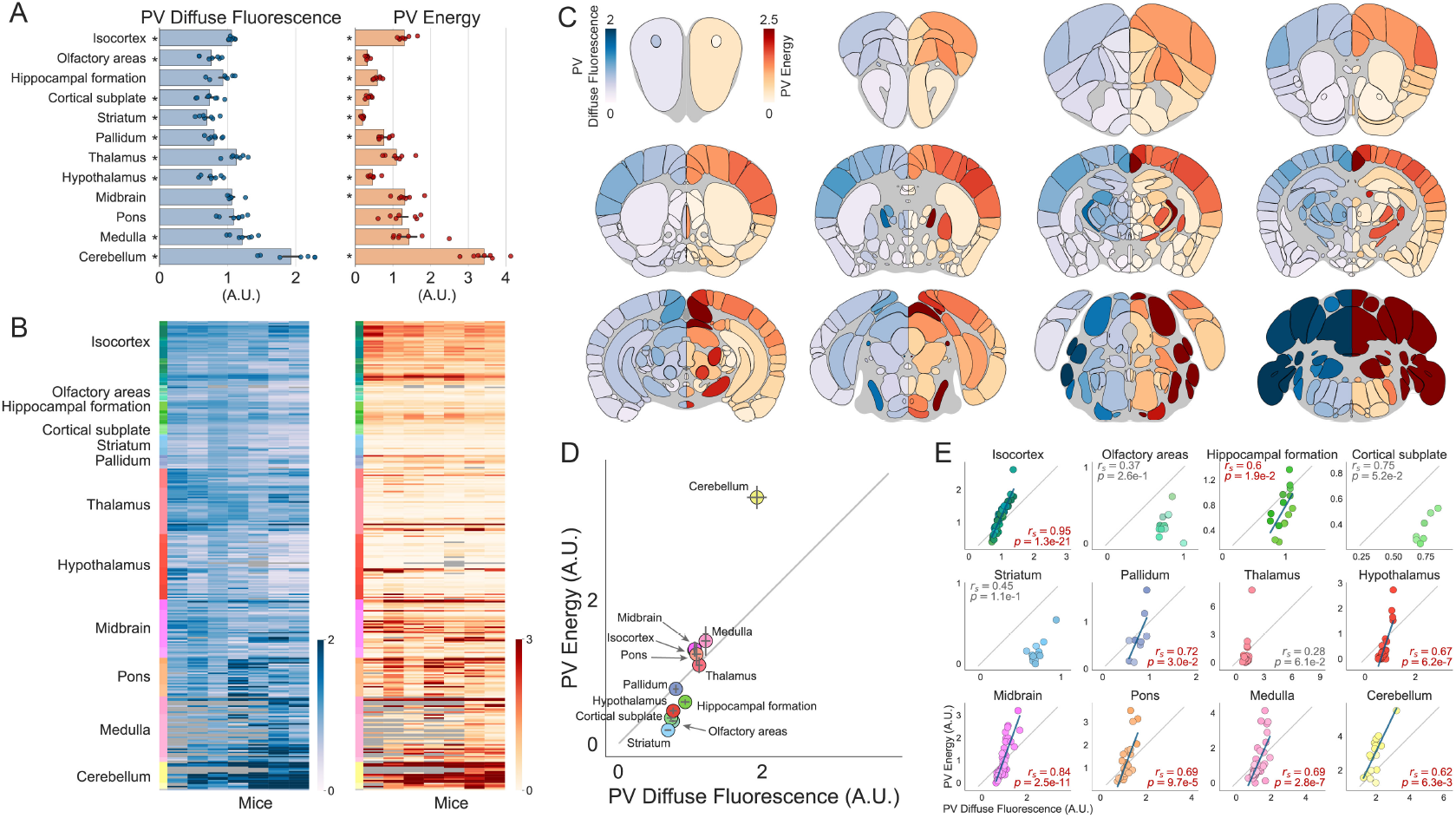
Distribution of PV-positive cells throughout the entire mouse brain. (**A**) Quantification of PV diffuse fluorescence and PV energy for 12 aggregated major brain subdivisions. Dots represent mice. Asterisks indicate brain subdivisions significantly different from the brain average (see ***Table ST1*** for statistical comparisons). (**B**) Heatmaps showing the two quantification metrics for mid-ontology brain regions in individual mice. Grayed-out cells represent brain regions where data is unavailable due to no sampling of that region. (**C**) Heatmaps showing coronal sections of the brain, sliced at different anteroposterior locations. On the left hemisphere (blue colormap) is displayed average diffuse PV fluorescence, while on the right hemisphere (red colormap) is displayed average PV energy for each brain region. (**D**) Plots of PV energy versus PV diffuse fluorescence for each of the 12 major brain subdivisions. (**E**) Same as in D but data is split in each brain region of the 12 major brain subdivisions. Error bars in A and D represent SEM across mice. In D and E, dots represent brain regions. In E, text insets indicate the Spearman correlation coefficient (*r_s_*) and the corresponding p-value, the gray line indicates the X-Y bisector, and, for significant correlations highlighted in red, the blue line shows the best linear fit.

**Figure S4.**
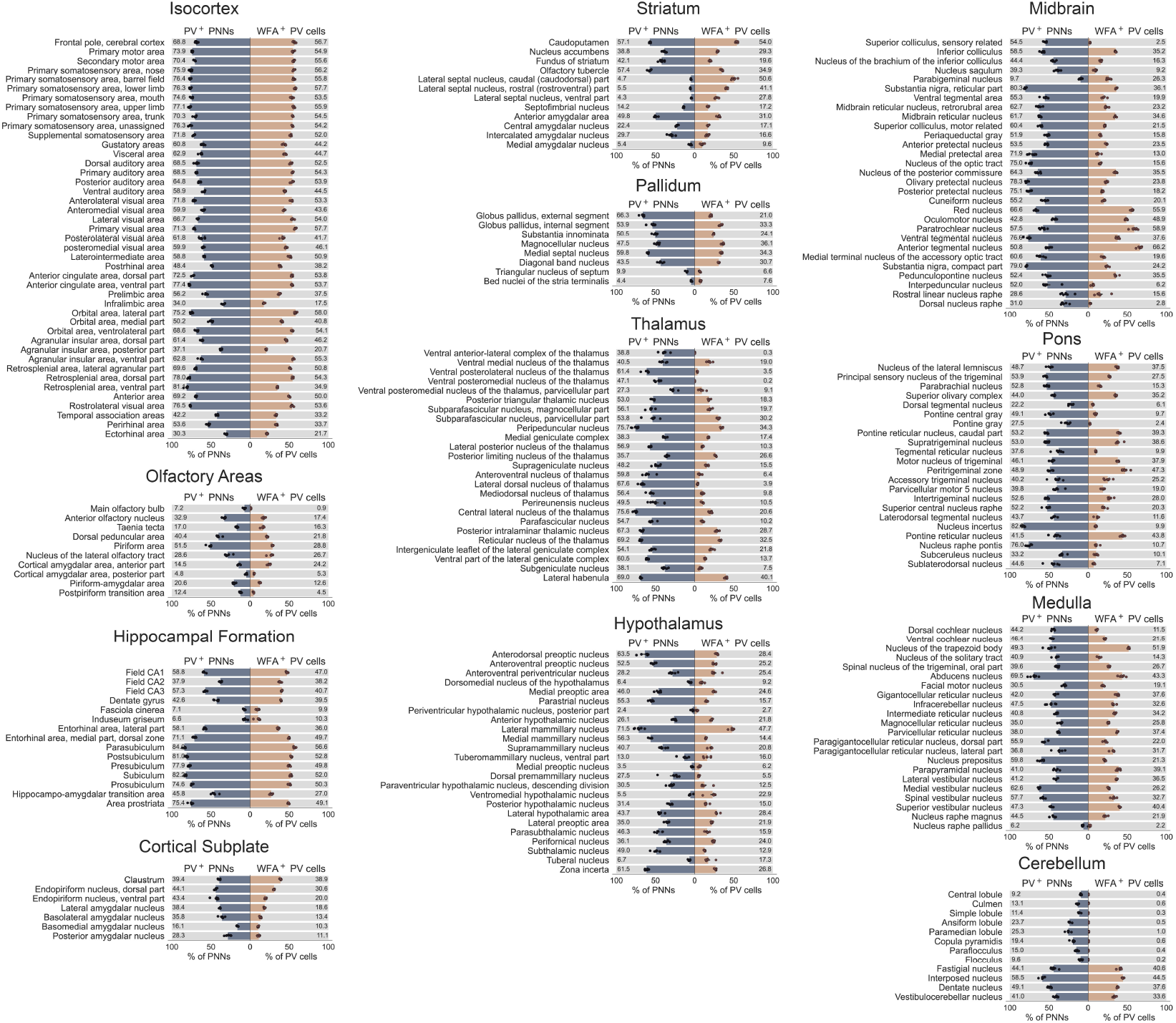
Colocalization of PNNs and PV cells in medium-resolution brain areas grouped by their major subdivision. For each plot, on the left in blue is represented the fraction of PNNs containing a PV cell (PV+ PNNs), while on the right in light orange is represented the fraction of PV cells surrounded by a PNN (WFA+ PV cells). In all the plots, dots represent “experimental units” and not single animals as described in the methods section “colocalization PNN-PV”. Each experimental unit is composed of the aggregated data of all mice in the dataset except one, in a manner similar to the leave-one-out cross-validation approach used in machine learning. This analysis includes only areas that had at least 3 PNNs and 3 PV cells in at least 4 mice. Error bars represent SEM across experimental units.

**Figure S5.**
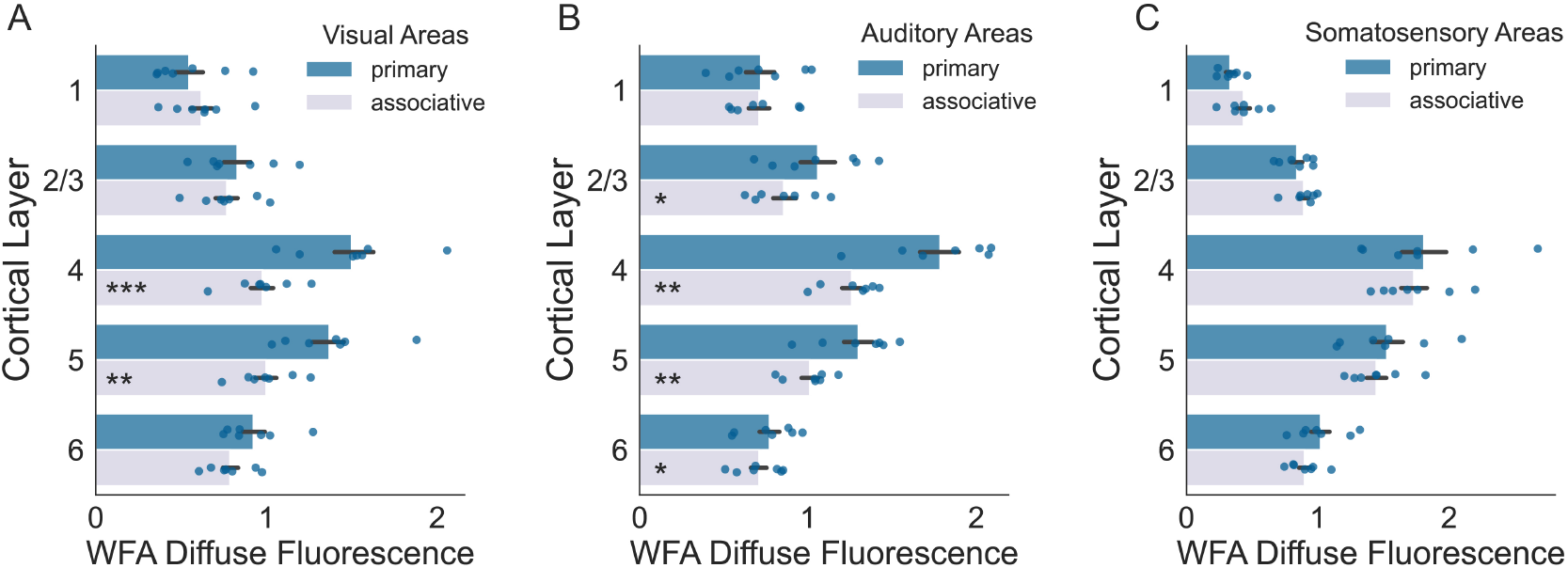
WFA Diffuse Fluorescence in primary vs secondary areas by layers. (**A**) WFA diffuse fluorescence in primary versus associative visual cortical areas split by layer. (**B**) Same as in (A) but for auditory areas. (**C**) Same as in (A) but for somatosensory areas. See ***Table ST1*** for statistical comparisons.

**Figure S6.**
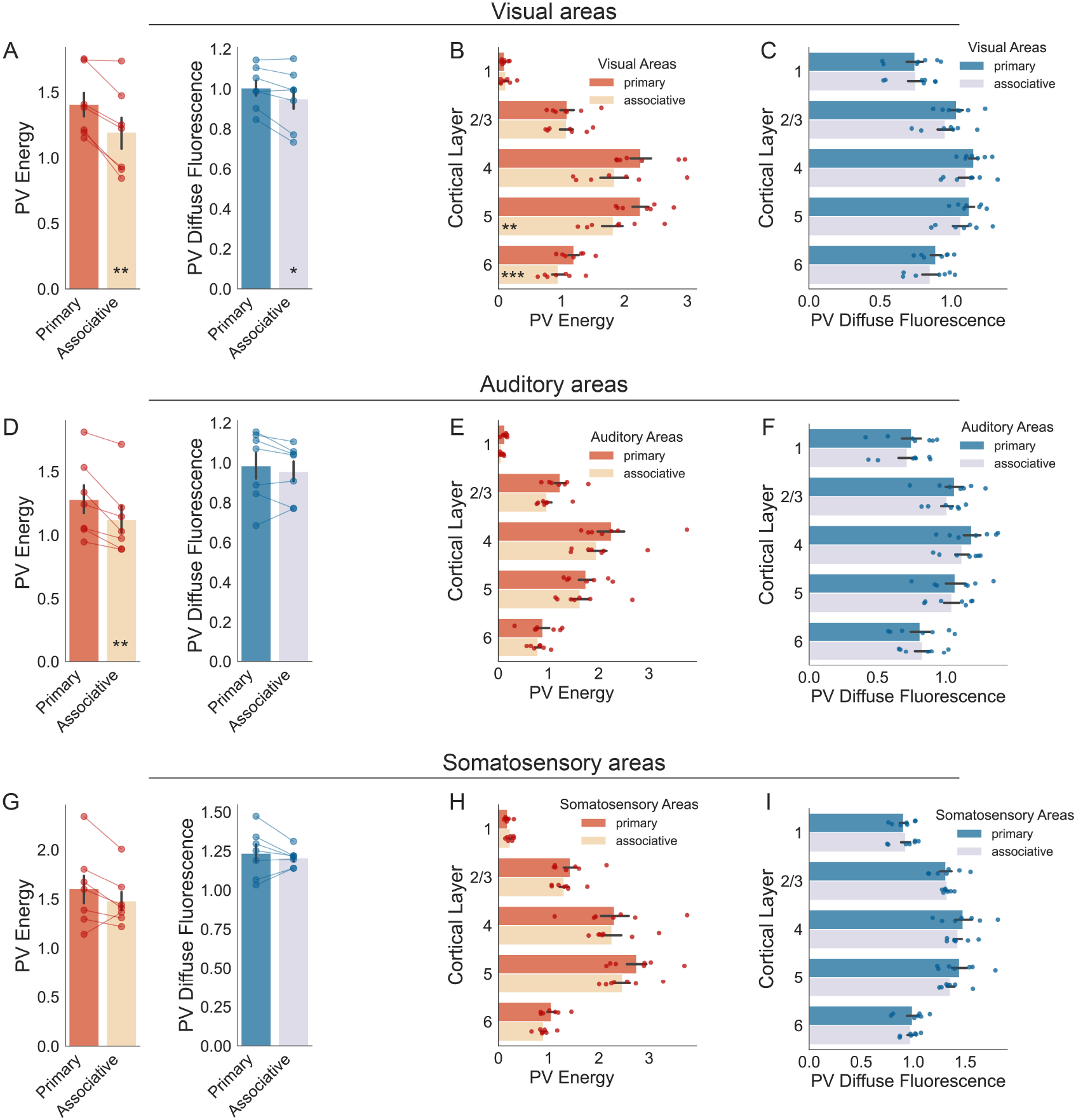
PV cell distribution in sensory cortical areas. (**A**) PV energy and PV diffuse fluorescence in primary versus associative visual areas. (**B**) PV energy and (**C**) PV diffuse fluorescence in primary versus associative visual areas split by layer. (**D**) Same as (A), but for auditory areas. (**E**) Same as (B), but for auditory areas. (**F**) Same as (C), but for auditory areas. (**G**) Same as (A), but for somatosensory areas. (**H**) Same as (B), but for somatosensory areas. (**I**) Same as (C) but for somatosensory areas. See ***Table ST1*** for statistical comparisons.

**Figure S7.**
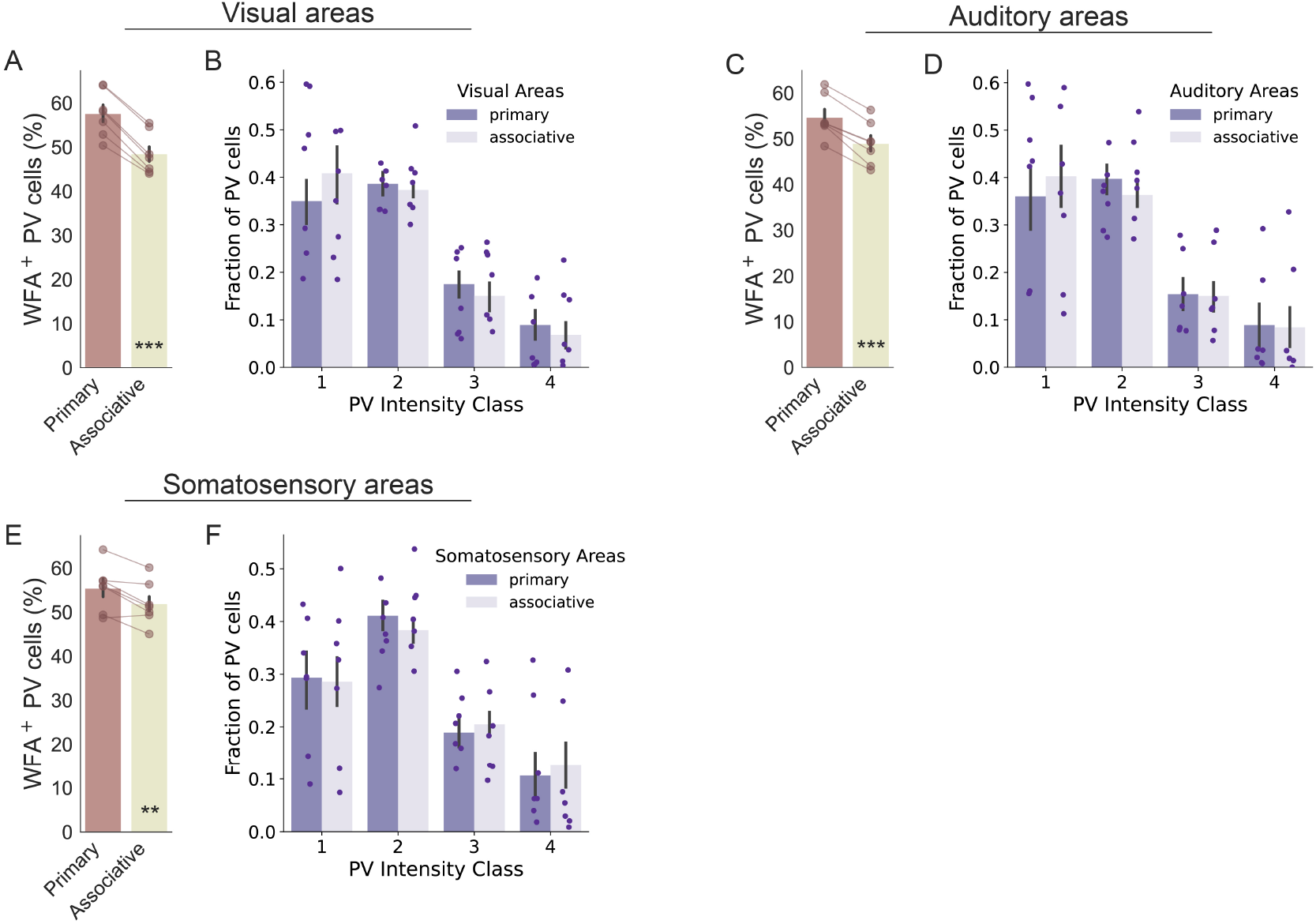
PV cell intensity and colocalization with PNNs in the sensory areas of the cortex. (**A**) Percentage of WFA+ PV cells in primary versus associative visual areas. (**B**) Distribution of PV cells in 4 intensity classes (low PV, intermediate-low PV, intermediate-high PV, and high PV) for primary versus associative visual areas. (**C**) Same as in (A) but for auditory areas. (**D**) Same as in (B) but for auditory areas. (**E**) Same as in (A) but for somatosensory areas. (**F**) Same as in (B) but for somatosensory areas. See ***Table ST1*** for statistical comparisons.

**Figure S8.**
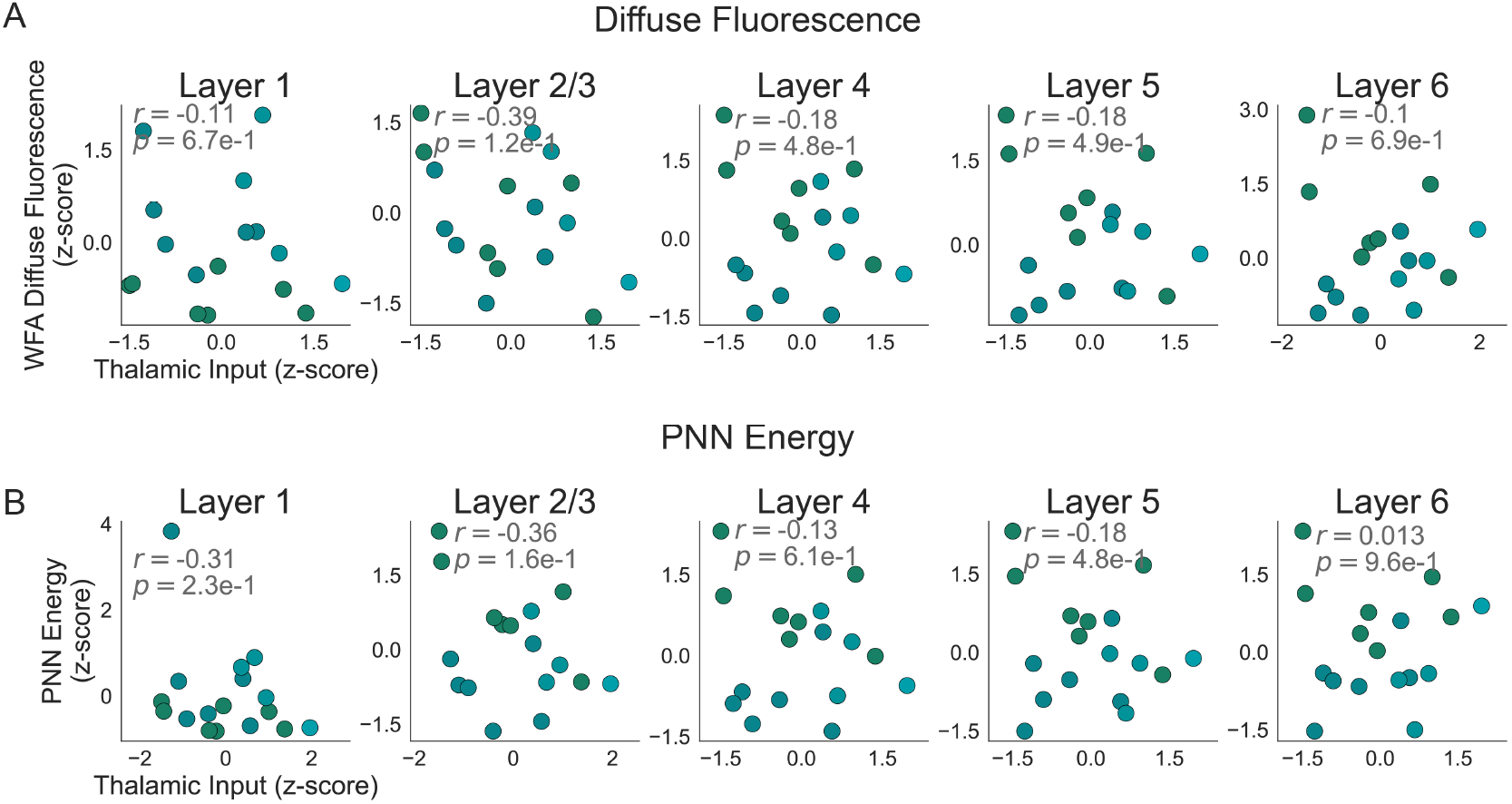
Thalamic inputs from the association-cortex-related portion of the thalamus (DORpm) do not correlate with PNNs in sensory cortices. (**A**) Correlation between WFA diffuse fluorescence and input strength of association-cortex-related thalamic areas (DORpm) in sensory-related cortices (all somatosensory, visual, and auditory cortices, see Correlation with thalamic afferent connectivity in Methods & Materials) split by layer. Text insets indicate the Pearson correlation coefficient (r) and the corresponding p-value. (**B**) Same as in (A) but for PNN energy.

**Figure S9.**
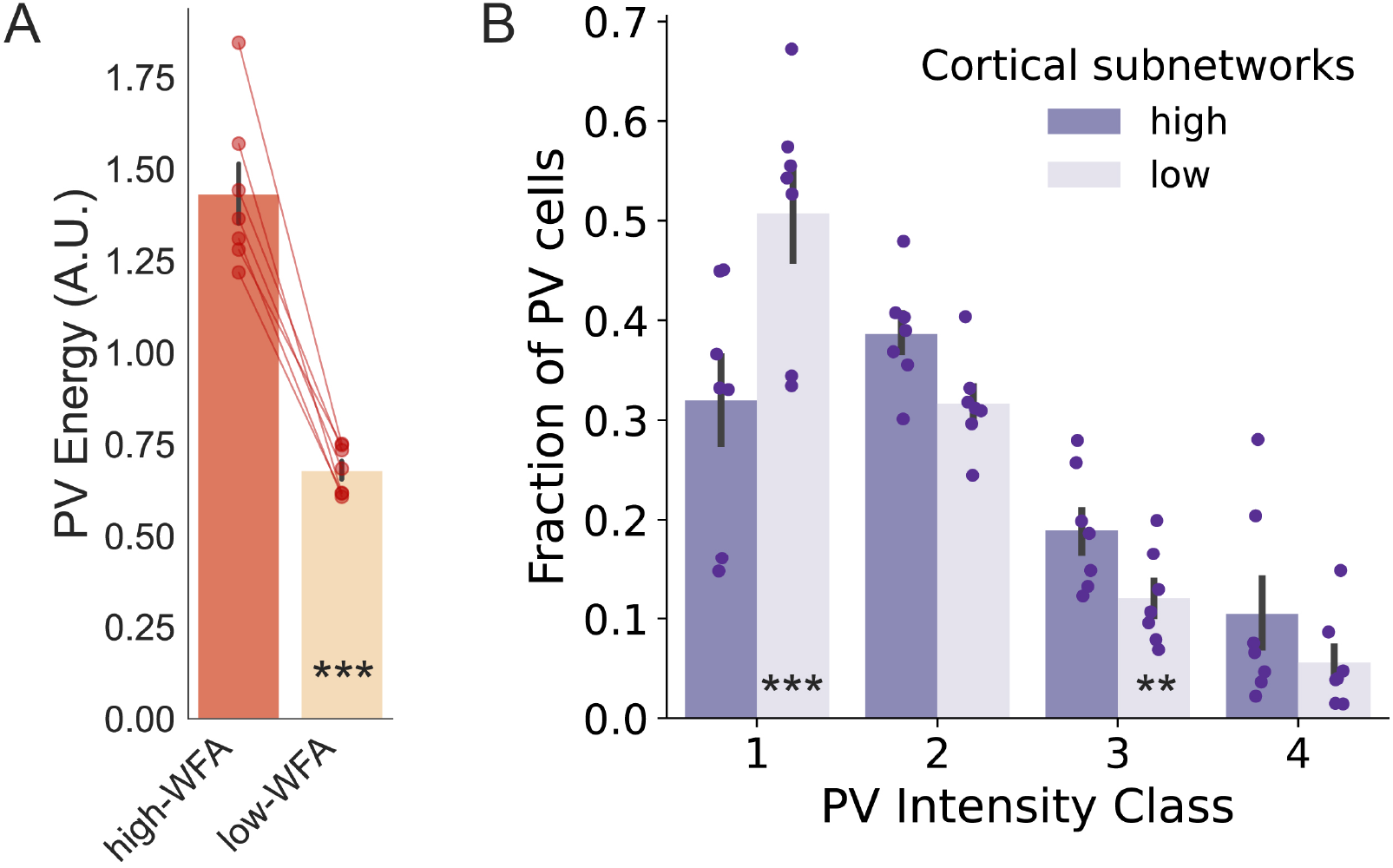
Properties of PV cells in high-WFA and low-WFA cortical subnetworks. (**A**) PV energy and (**B**) Distribution of PV cells in 4 intensity classes (1: low PV, 2: intermediate-low PV, 3: intermediate-high PV, and 4: high PV) for high-WFA and low-WFA cortical subnetworks, as defined in Fig.5. See ***Table ST1*** for statistical comparisons.

### Supplementary Tables

**Table ST1.**
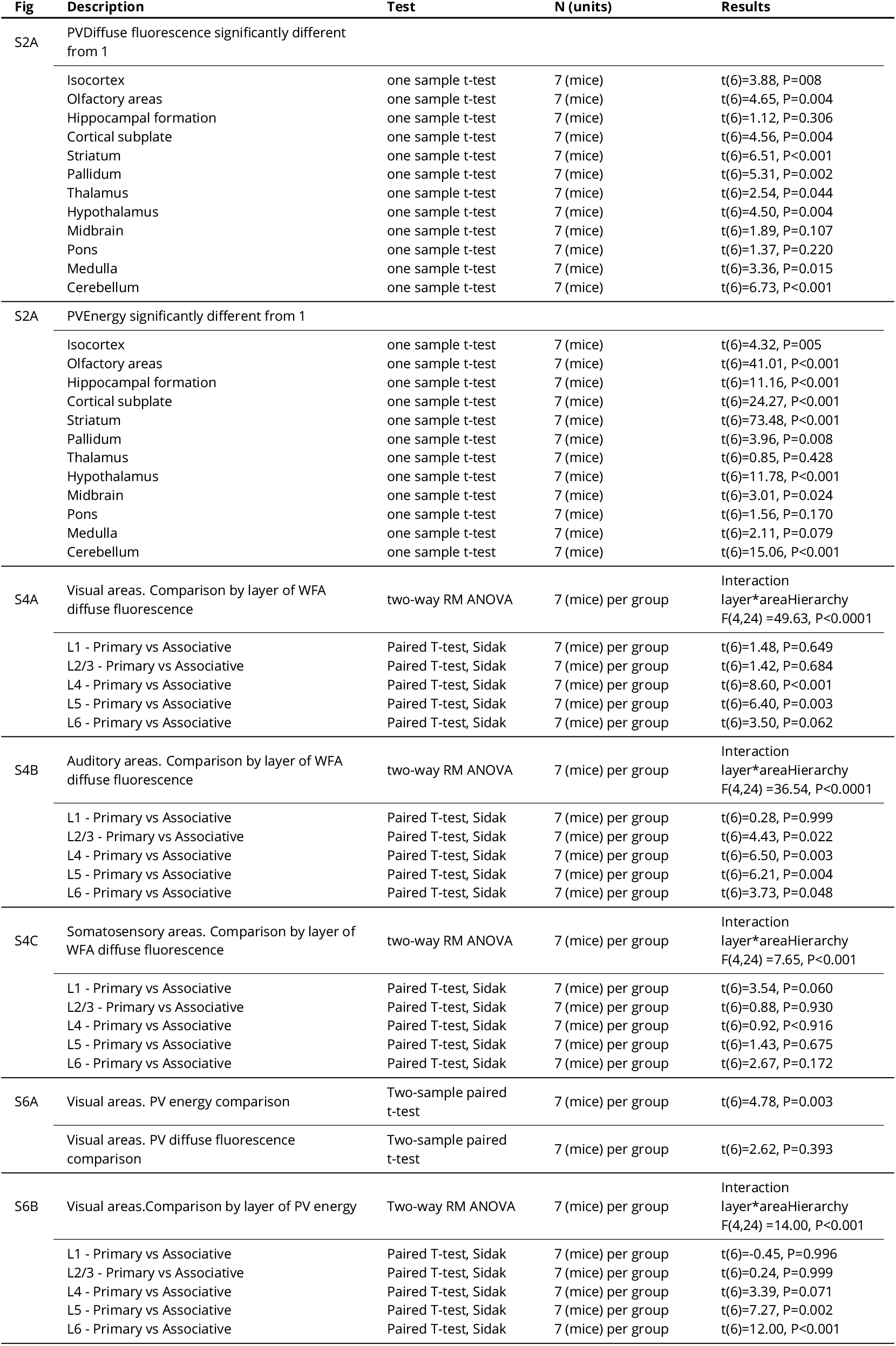
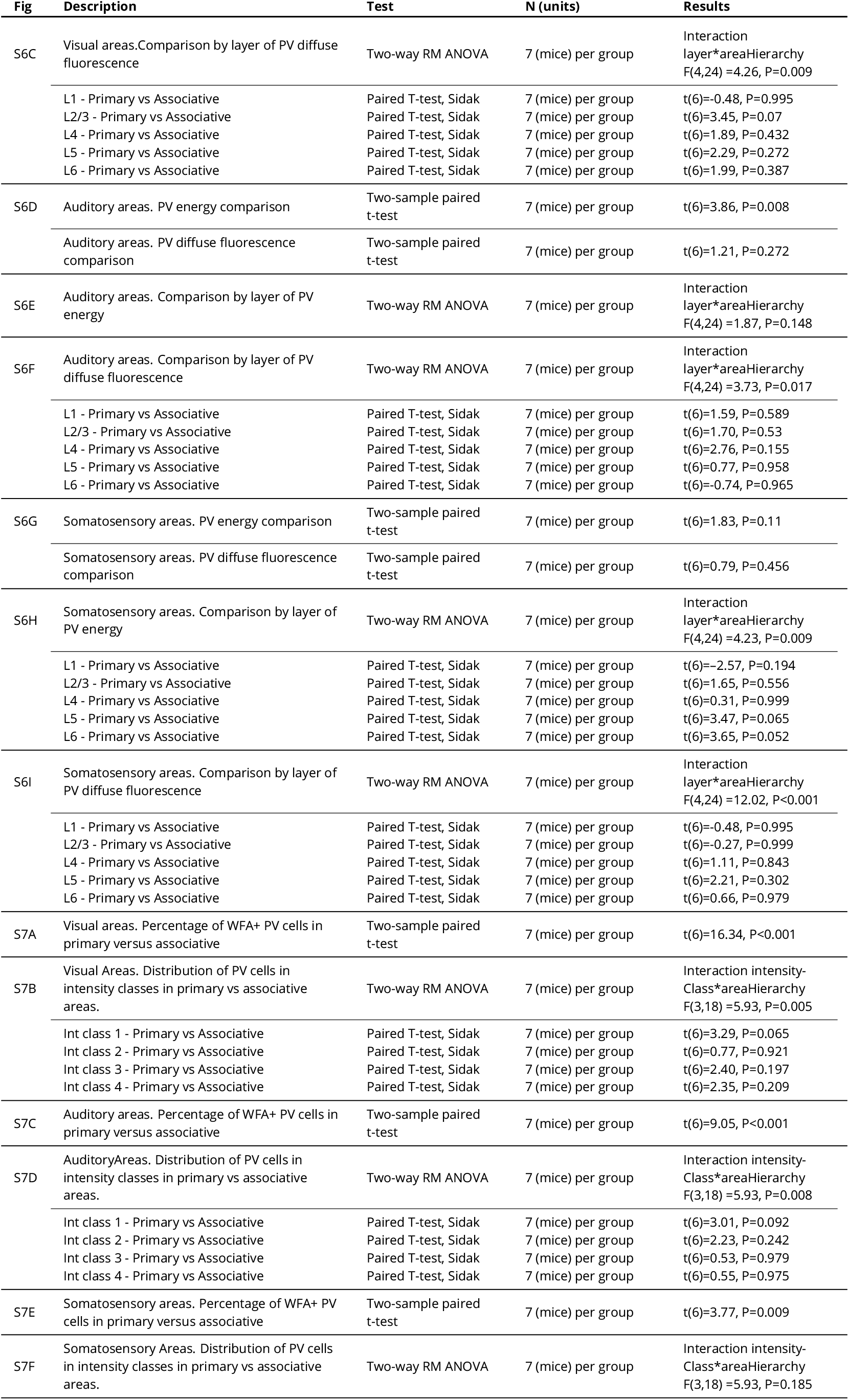
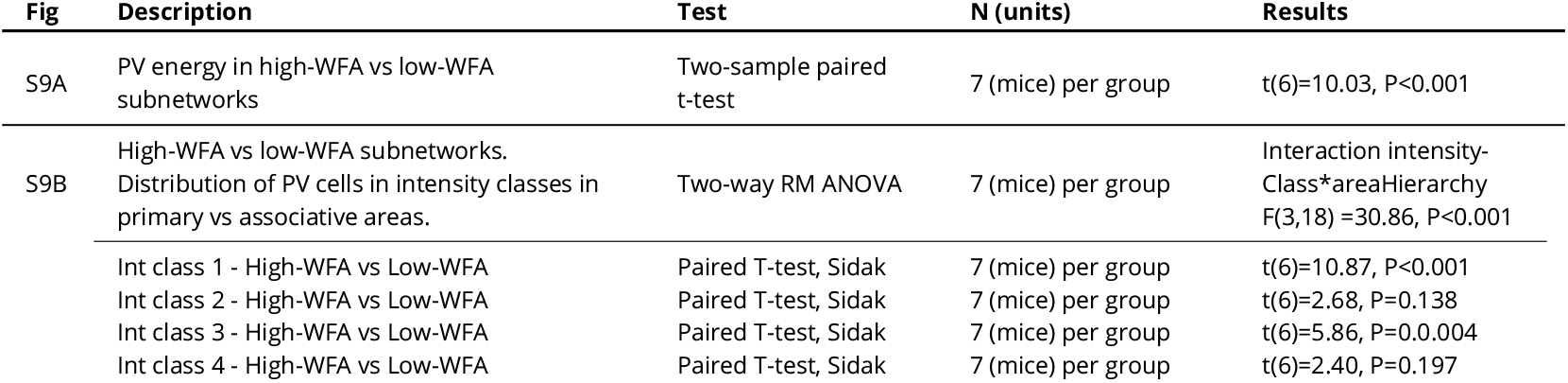
Statistical comparisons.

| Fig | Description | Test | N (units) | Results |
| --- | --- | --- | --- | --- |
| S2A | PV Diffuse fluorescence significantly different from 1 |  |  |  |
|  | Isocortex | one sample t-test | 7 (mice) | t(6)=3.88, P=0.08 |
|  | Olfactory areas | one sample t-test | 7 (mice) | t(6)=4.65, P=0.004 |
|  | Hippocampal formation | one sample t-test | 7 (mice) | t(6)=1.12, P=0.306 |
|  | Cortical subplate | one sample t-test | 7 (mice) | t(6)=4.56, P=0.004 |
|  | Striatum | one sample t-test | 7 (mice) | t(6)=6.51, P<0.001 |
|  | Pallidum | one sample t-test | 7 (mice) | t(6)=5.31, P=0.002 |
|  | Thalamus | one sample t-test | 7 (mice) | t(6)=2.54, P=0.044 |
|  | Hypothalamus | one sample t-test | 7 (mice) | t(6)=4.50, P=0.004 |
|  | Midbrain | one sample t-test | 7 (mice) | t(6)=1.89, P=0.107 |
|  | Pons | one sample t-test | 7 (mice) | t(6)=1.37, P=0.220 |
|  | Medulla | one sample t-test | 7 (mice) | t(6)=3.36, P=0.015 |
|  | Cerebellum | one sample t-test | 7 (mice) | t(6)=6.73, P<0.001 |
| S2A | PV Energy significantly different from 1 |  |  |  |
|  | Isocortex | one sample t-test | 7 (mice) | t(6)=4.32, P=0.05 |
|  | Olfactory areas | one sample t-test | 7 (mice) | t(6)=41.01, P<0.001 |
|  | Hippocampal formation | one sample t-test | 7 (mice) | t(6)=11.16, P<0.001 |
|  | Cortical subplate | one sample t-test | 7 (mice) | t(6)=24.27, P<0.001 |
|  | Striatum | one sample t-test | 7 (mice) | t(6)=73.48, P<0.001 |
|  | Pallidum | one sample t-test | 7 (mice) | t(6)=3.96, P=0.008 |
|  | Thalamus | one sample t-test | 7 (mice) | t(6)=0.85, P=0.428 |
|  | Hypothalamus | one sample t-test | 7 (mice) | t(6)=11.78, P<0.001 |
|  | Midbrain | one sample t-test | 7 (mice) | t(6)=3.01, P=0.024 |
|  | Pons | one sample t-test | 7 (mice) | t(6)=1.56, P=0.170 |
|  | Medulla | one sample t-test | 7 (mice) | t(6)=2.11, P=0.079 |
|  | Cerebellum | one sample t-test | 7 (mice) | t(6)=15.06, P<0.001 |
| S4A | Visual areas. Comparison by layer of WFA diffuse fluorescence | two-way RM ANOVA | 7 (mice) per group | Interaction layer*areaHierarchy<br>F(4,24) =49.63, P<0.0001 |
|  | L1 - Primary vs Associative | Paired T-test, Sidak | 7 (mice) per group | t(6)=1.48, P=0.649 |
|  | L2/3 - Primary vs Associative | Paired T-test, Sidak | 7 (mice) per group | t(6)=1.42, P=0.684 |
|  | L4 - Primary vs Associative | Paired T-test, Sidak | 7 (mice) per group | t(6)=8.60, P<0.001 |
|  | L5 - Primary vs Associative | Paired T-test, Sidak | 7 (mice) per group | t(6)=6.40, P=0.003 |
|  | L6 - Primary vs Associative | Paired T-test, Sidak | 7 (mice) per group | t(6)=3.50, P=0.062 |
| S4B | Auditory areas. Comparison by layer of WFA diffuse fluorescence | two-way RM ANOVA | 7 (mice) per group | Interaction layer*areaHierarchy<br>F(4,24) =36.54, P<0.0001 |
|  | L1 - Primary vs Associative | Paired T-test, Sidak | 7 (mice) per group | t(6)=0.28, P=0.999 |
|  | L2/3 - Primary vs Associative | Paired T-test, Sidak | 7 (mice) per group | t(6)=4.43, P=0.022 |
|  | L4 - Primary vs Associative | Paired T-test, Sidak | 7 (mice) per group | t(6)=6.50, P=0.003 |
|  | L5 - Primary vs Associative | Paired T-test, Sidak | 7 (mice) per group | t(6)=6.21, P=0.004 |
|  | L6 - Primary vs Associative | Paired T-test, Sidak | 7 (mice) per group | t(6)=3.73, P=0.048 |
| S4C | Somatosensory areas. Comparison by layer of WFA diffuse fluorescence | two-way RM ANOVA | 7 (mice) per group | Interaction layer*areaHierarchy<br>F(4,24) =7.65, P<0.001 |
|  | L1 - Primary vs Associative | Paired T-test, Sidak | 7 (mice) per group | t(6)=3.54, P=0.060 |
|  | L2/3 - Primary vs Associative | Paired T-test, Sidak | 7 (mice) per group | t(6)=0.88, P=0.930 |
|  | L4 - Primary vs Associative | Paired T-test, Sidak | 7 (mice) per group | t(6)=0.92, P<0.916 |
|  | L5 - Primary vs Associative | Paired T-test, Sidak | 7 (mice) per group | t(6)=1.43, P=0.675 |
|  | L6 - Primary vs Associative | Paired T-test, Sidak | 7 (mice) per group | t(6)=2.67, P=0.172 |
| S6A | Visual areas. PV energy comparison | Two-sample paired t-test | 7 (mice) per group | t(6)=4.78, P=0.003 |
|  | Visual areas. PV diffuse fluorescence comparison | Two-sample paired t-test | 7 (mice) per group | t(6)=2.62, P=0.393 |
| S6B | Visual areas. Comparison by layer of PV energy | Two-way RM ANOVA | 7 (mice) per group | Interaction layer*areaHierarchy<br>F(4,24) =14.00, P<0.001 |
|  | L1 - Primary vs Associative | Paired T-test, Sidak | 7 (mice) per group | t(6)= -0.45, P=0.996 |
|  | L2/3 - Primary vs Associative | Paired T-test, Sidak | 7 (mice) per group | t(6)=0.24, P=0.999 |
|  | L4 - Primary vs Associative | Paired T-test, Sidak | 7 (mice) per group | t(6)=3.39, P=0.071 |
|  | L5 - Primary vs Associative | Paired T-test, Sidak | 7 (mice) per group | t(6)=7.27, P=0.002 |
|  | L6 - Primary vs Associative | Paired T-test, Sidak | 7 (mice) per group | t(6)=12.00, P<0.001 |
| S6C | Visual areas. Comparison by layer of PV diffuse fluorescence | Two-way RM ANOVA | 7 (mice) per group | Interaction layer*areaHierarchy<br>F(4,24) =4.26, P=0.009 |
|  | L1 - Primary vs Associative | Paired T-test, Sidak | 7 (mice) per group | t(6)=-0.48, P=0.995 |
|  | L2/3 - Primary vs Associative | Paired T-test, Sidak | 7 (mice) per group | t(6)=3.45, P=0.07 |
|  | L4 - Primary vs Associative | Paired T-test, Sidak | 7 (mice) per group | t(6)=1.89, P=0.432 |
|  | L5 - Primary vs Associative | Paired T-test, Sidak | 7 (mice) per group | t(6)=2.29, P=0.272 |
|  | L6 - Primary vs Associative | Paired T-test, Sidak | 7 (mice) per group | t(6)=1.99, P=0.387 |
| S6D | Auditory areas. PV energy comparison | Two-sample paired t-test | 7 (mice) per group | t(6)=3.86, P=0.008 |
|  | Auditory areas. PV diffuse fluorescence comparison | Two-sample paired t-test | 7 (mice) per group | t(6)=1.21, P=0.272 |
| S6E | Auditory areas. Comparison by layer of PV energy | Two-way RM ANOVA | 7 (mice) per group | Interaction layer*areaHierarchy<br>F(4,24) =1.87, P=0.148 |
| S6F | Auditory areas. Comparison by layer of PV diffuse fluorescence | Two-way RM ANOVA | 7 (mice) per group | Interaction layer*areaHierarchy<br>F(4,24) =3.73, P=0.017 |
|  | L1 - Primary vs Associative | Paired T-test, Sidak | 7 (mice) per group | t(6)=1.59, P=0.589 |
|  | L2/3 - Primary vs Associative | Paired T-test, Sidak | 7 (mice) per group | t(6)=1.70, P=0.53 |
|  | L4 - Primary vs Associative | Paired T-test, Sidak | 7 (mice) per group | t(6)=2.76, P=0.155 |
|  | L5 - Primary vs Associative | Paired T-test, Sidak | 7 (mice) per group | t(6)=0.77, P=0.958 |
|  | L6 - Primary vs Associative | Paired T-test, Sidak | 7 (mice) per group | t(6)=-0.74, P=0.965 |
| S6G | Somatosensory areas. PV energy comparison | Two-sample paired t-test | 7 (mice) per group | t(6)=1.83, P=0.11 |
|  | Somatosensory areas. PV diffuse fluorescence comparison | Two-sample paired t-test | 7 (mice) per group | t(6)=0.79, P=0.456 |
| S6H | Somatosensory areas. Comparison by layer of PV energy | Two-way RM ANOVA | 7 (mice) per group | Interaction layer*areaHierarchy<br>F(4,24) =4.23, P=0.009 |
|  | L1 - Primary vs Associative | Paired T-test, Sidak | 7 (mice) per group | t(6)=-2.57, P=0.194 |
|  | L2/3 - Primary vs Associative | Paired T-test, Sidak | 7 (mice) per group | t(6)=1.65, P=0.556 |
|  | L4 - Primary vs Associative | Paired T-test, Sidak | 7 (mice) per group | t(6)=0.31, P=0.999 |
|  | L5 - Primary vs Associative | Paired T-test, Sidak | 7 (mice) per group | t(6)=3.47, P=0.065 |
|  | L6 - Primary vs Associative | Paired T-test, Sidak | 7 (mice) per group | t(6)=3.65, P=0.052 |
| S6I | Somatosensory areas. Comparison by layer of PV diffuse fluorescence | Two-way RM ANOVA | 7 (mice) per group | Interaction layer*areaHierarchy<br>F(4,24) =12.02, P<0.001 |
|  | L1 - Primary vs Associative | Paired T-test, Sidak | 7 (mice) per group | t(6)=-0.48, P=0.995 |
|  | L2/3 - Primary vs Associative | Paired T-test, Sidak | 7 (mice) per group | t(6)=-0.27, P=0.999 |
|  | L4 - Primary vs Associative | Paired T-test, Sidak | 7 (mice) per group | t(6)=1.11, P=0.843 |
|  | L5 - Primary vs Associative | Paired T-test, Sidak | 7 (mice) per group | t(6)=2.21, P=0.302 |
|  | L6 - Primary vs Associative | Paired T-test, Sidak | 7 (mice) per group | t(6)=0.66, P=0.979 |
| S7A | Visual areas. Percentage of WFA+ PV cells in primary versus associative | Two-sample paired t-test | 7 (mice) per group | t(6)=16.34, P<0.001 |
| S7B | Visual Areas. Distribution of PV cells in intensity classes in primary vs associative areas. | Two-way RM ANOVA | 7 (mice) per group | Interaction intensity-Class*areaHierarchy<br>F(3,18) =5.93, P=0.005 |
|  | Int class 1 - Primary vs Associative | Paired T-test, Sidak | 7 (mice) per group | t(6)=3.29, P=0.065 |
|  | Int class 2 - Primary vs Associative | Paired T-test, Sidak | 7 (mice) per group | t(6)=0.77, P=0.921 |
|  | Int class 3 - Primary vs Associative | Paired T-test, Sidak | 7 (mice) per group | t(6)=2.40, P=0.197 |
|  | Int class 4 - Primary vs Associative | Paired T-test, Sidak | 7 (mice) per group | t(6)=2.35, P=0.209 |
| S7C | Auditory areas. Percentage of WFA+ PV cells in primary versus associative | Two-sample paired t-test | 7 (mice) per group | t(6)=9.05, P<0.001 |
| S7D | Auditory Areas. Distribution of PV cells in intensity classes in primary vs associative areas. | Two-way RM ANOVA | 7 (mice) per group | Interaction intensity-Class*areaHierarchy<br>F(3,18) =5.93, P=0.008 |
|  | Int class 1 - Primary vs Associative | Paired T-test, Sidak | 7 (mice) per group | t(6)=3.01, P=0.092 |
|  | Int class 2 - Primary vs Associative | Paired T-test, Sidak | 7 (mice) per group | t(6)=2.23, P=0.242 |
|  | Int class 3 - Primary vs Associative | Paired T-test, Sidak | 7 (mice) per group | t(6)=0.53, P=0.979 |
|  | Int class 4 - Primary vs Associative | Paired T-test, Sidak | 7 (mice) per group | t(6)=0.55, P=0.975 |
| S7E | Somatosensory areas. Percentage of WFA+ PV cells in primary versus associative | Two-sample paired t-test | 7 (mice) per group | t(6)=3.77, P=0.009 |
| S7F | Somatosensory Areas. Distribution of PV cells in intensity classes in primary vs associative areas. | Two-way RM ANOVA | 7 (mice) per group | Interaction intensity-Class*areaHierarchy<br>F(3,18) =5.93, P=0.185 |
| S9A | PV energy in high-WFA vs low-WFA subnetworks | Two-sample paired t-test | 7 (mice) per group | t(6)=10.03, P<0.001 |
| S9B | High-WFA vs low-WFA subnetworks. Distribution of PV cells in intensity classes in primary vs associative areas. | Two-way RM ANOVA | 7 (mice) per group | Interaction intensity-Class*areaHierarchy F(3,18) =30.86, P<0.001 |
|  | Int class 1 - High-WFA vs Low-WFA | Paired T-test, Sidak | 7 (mice) per group | t(6)=10.87, P<0.001 |
|  | Int class 2 - High-WFA vs Low-WFA | Paired T-test, Sidak | 7 (mice) per group | t(6)=2.68, P=0.138 |
|  | Int class 3 - High-WFA vs Low-WFA | Paired T-test, Sidak | 7 (mice) per group | t(6)=5.86, P=0.0004 |
|  | Int class 4 - High-WFA vs Low-WFA | Paired T-test, Sidak | 7 (mice) per group | t(6)=2.40, P=0.197 |

**Table ST2.** Coarse-ontology brain regions.

| Name | Acronym | regionID |
| --- | --- | --- |
| Isocortex | Isocortex | 315 |
| Olfactory areas | OLF | 698 |
| Hippocampal formation | HPF | 1089 |
| Cortical subplate | CTXsp | 703 |
| Striatum | STR | 477 |
| Pallidum | PAL | 803 |
| Thalamus | TH | 549 |
| Hypothalamus | HY | 1097 |
| Midbrain | MB | 313 |
| Pons | P | 771 |
| Medulla | MY | 354 |
| Cerebellum | CB | 512 |

**Table ST3.** Cortical regions.

| Name | Acronym | regionID |
| --- | --- | --- |
| Frontal pole, cerebral cortex | FRP | 184 |
| Primary motor area | MOp | 985 |
| Secondary motor area | MOs | 993 |
| Primary somatosensory area, nose | SSp-n | 353 |
| Primary somatosensory area, barrel field | SSp-bfd | 329 |
| Primary somatosensory area, lower limb | SSp-lI | 337 |
| Primary somatosensory area, mouth | SSp-m | 345 |
| Primary somatosensory area, upper limb | SSp-ul | 369 |
| Primary somatosensory area, trunk | SSp-tr | 361 |
| Primary somatosensory area, unassigned | SSp-un | 182305689 |
| Supplemental somatosensory area | SSs | 378 |
| Anterolateral visual area | VISal | 402 |
| Anteromedial visual area | VISam | 394 |
| Lateral visual area | VISl | 409 |
| Primary visual area | VISp | 385 |
| Posterolateral visual area | VISpl | 425 |
| posteromedial visual area | VISpm | 533 |
| Laterointermediate area | VISli | 312782574 |
| Postrhinal area | VISpor | 312782628 |
| Anterior area | VISa | 312782546 |
| Rostrolateral visual area | VISrl | 417 |
| Dorsal auditory area | AUDd | 1011 |
| Primary auditory area | AUDp | 1002 |
| Posterior auditory area | AUDpo | 1027 |
| Ventral auditory area | AUDv | 1018 |
| Infralimbic area | ILA | 44 |
| Prelimbic area | PL | 972 |
| Orbital area, medial part | ORBm | 731 |
| Orbital area, lateral part | ORBI | 723 |
| Orbital area, ventrolateral part | ORBvl | 746 |
| Anterior cingulate area, dorsal part | ACAd | 39 |
| Anterior cingulate area, ventral part | ACAv | 48 |
| Retrosplenial area, lateral agranular part | RSPagl | 894 |
| Retrosplenial area, dorsal part | RSPd | 879 |
| Retrosplenial area, ventral part | RSPv | 886 |
| Gustatory areas | GU | 1057 |
| Visceral area | VISC | 677 |
| Agranular insular area, dorsal part | Ald | 104 |
| Agranular insular area, posterior part | Alp | 111 |
| Agranular insular area, ventral part | Alv | 119 |
| Temporal association areas | TEa | 541 |
| Ectorhinal area | ECT | 895 |
| Perirhinal area | PERI | 922 |
| Entorhinal area, lateral part | ENTI | 918 |
| Entorhinal area, medial part, dorsal zone | ENTm | 926 |

**Table ST4.**
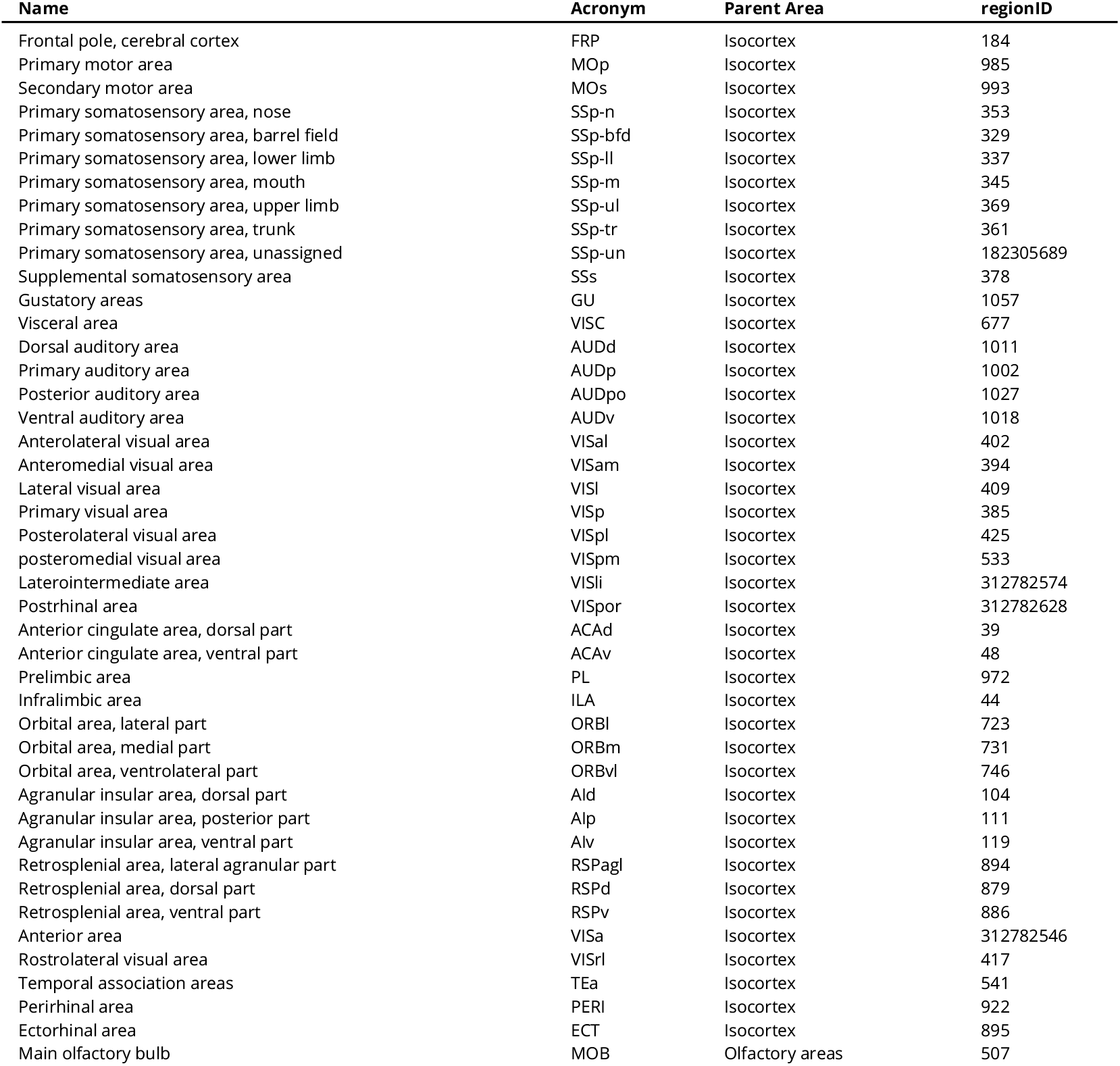
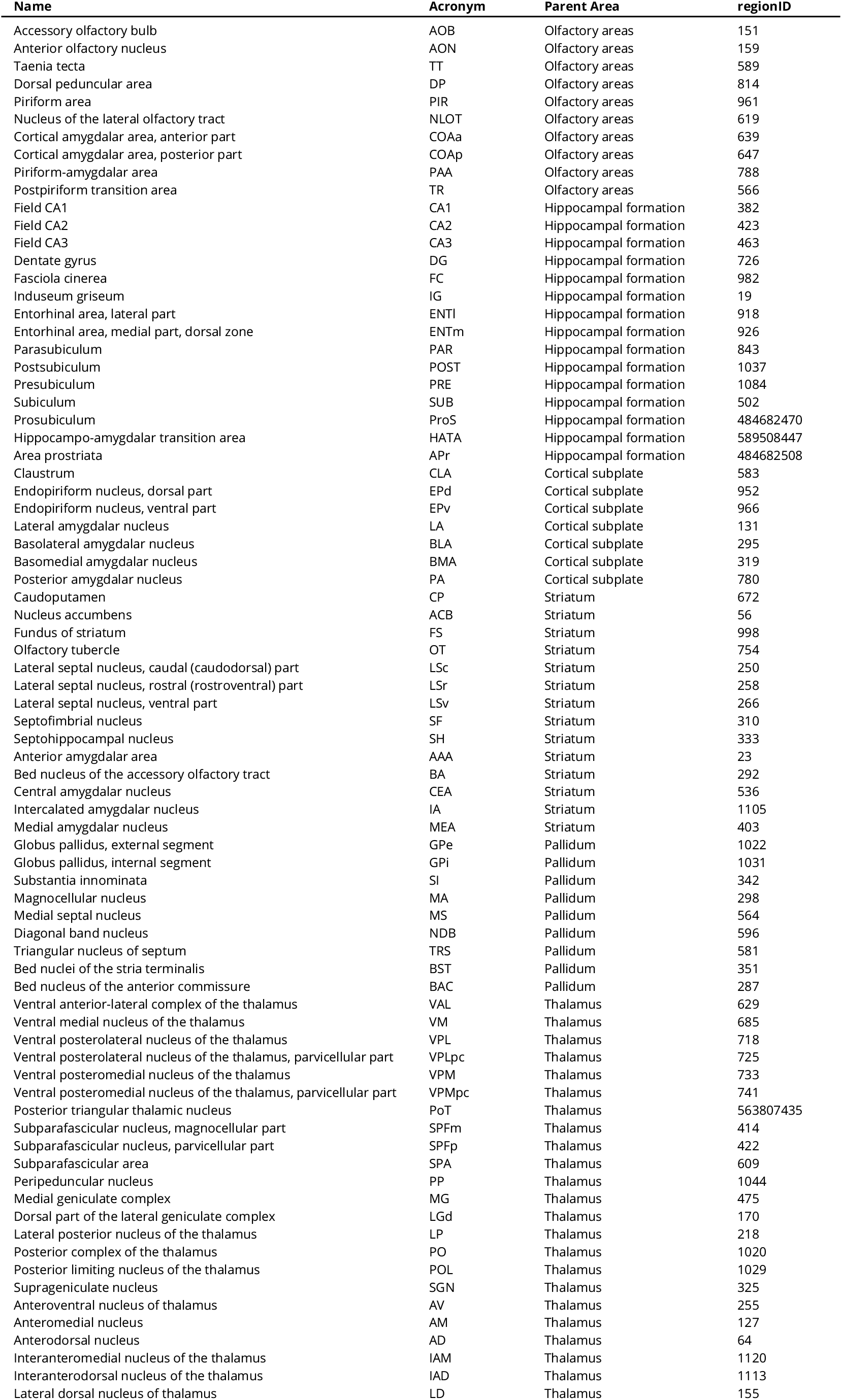
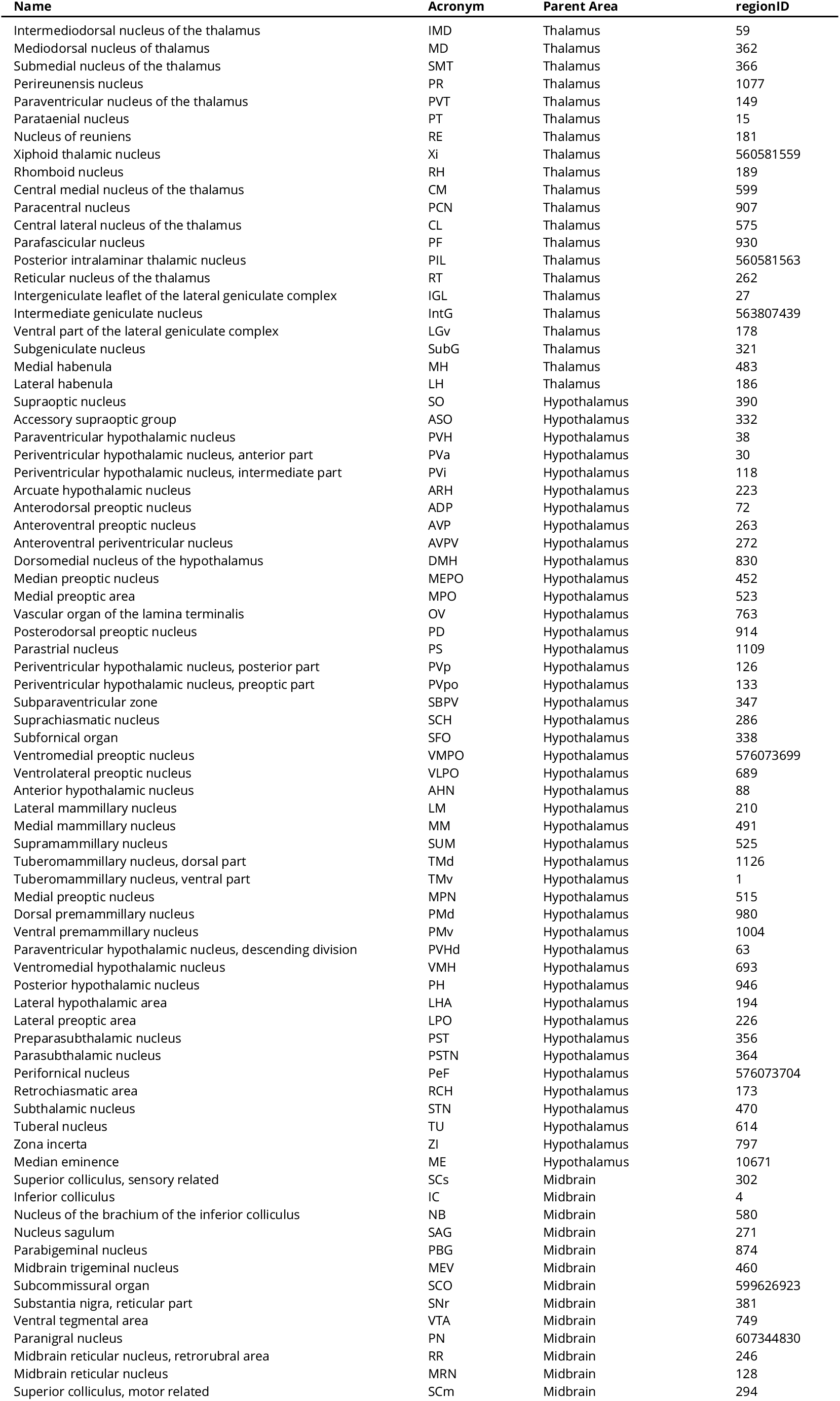
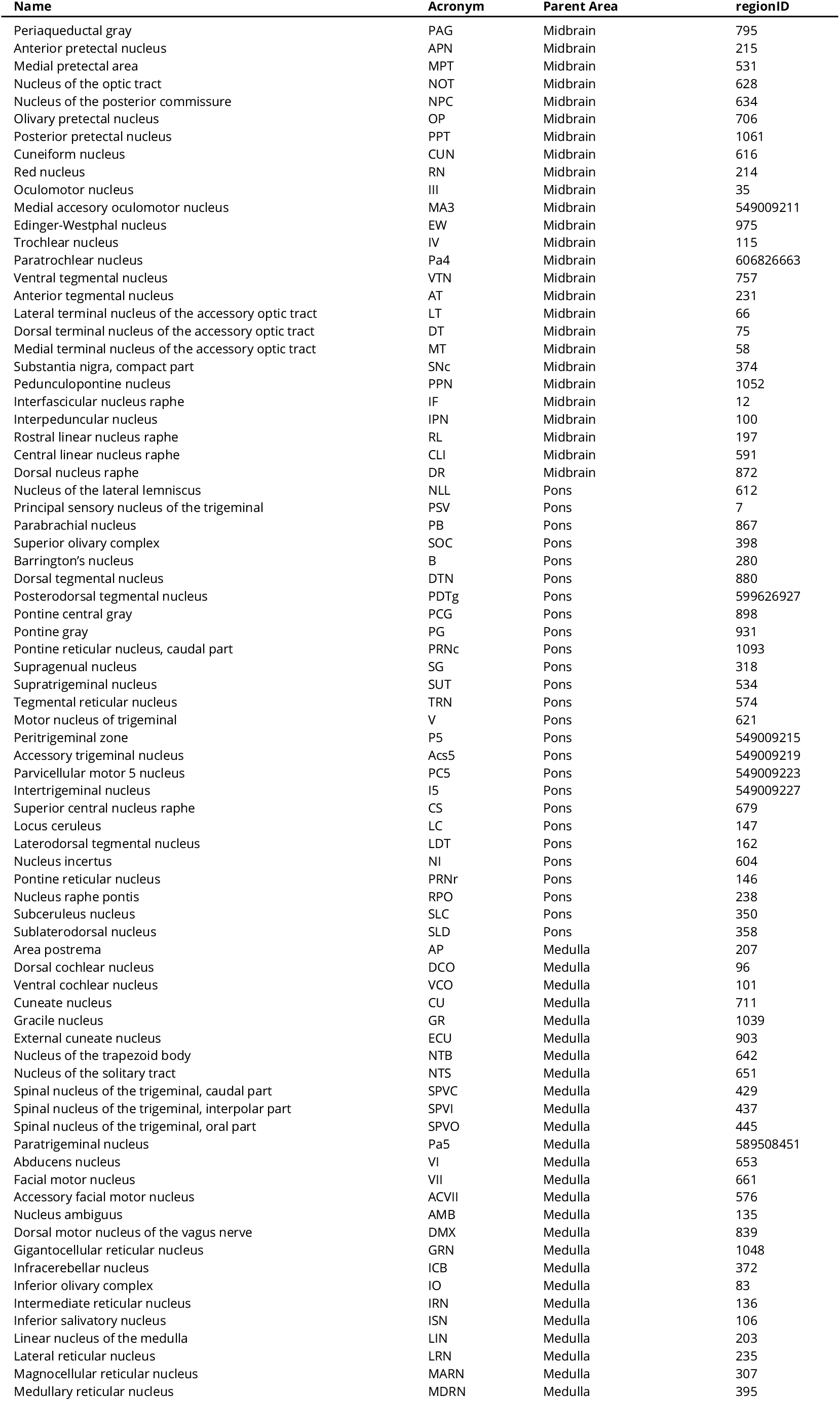
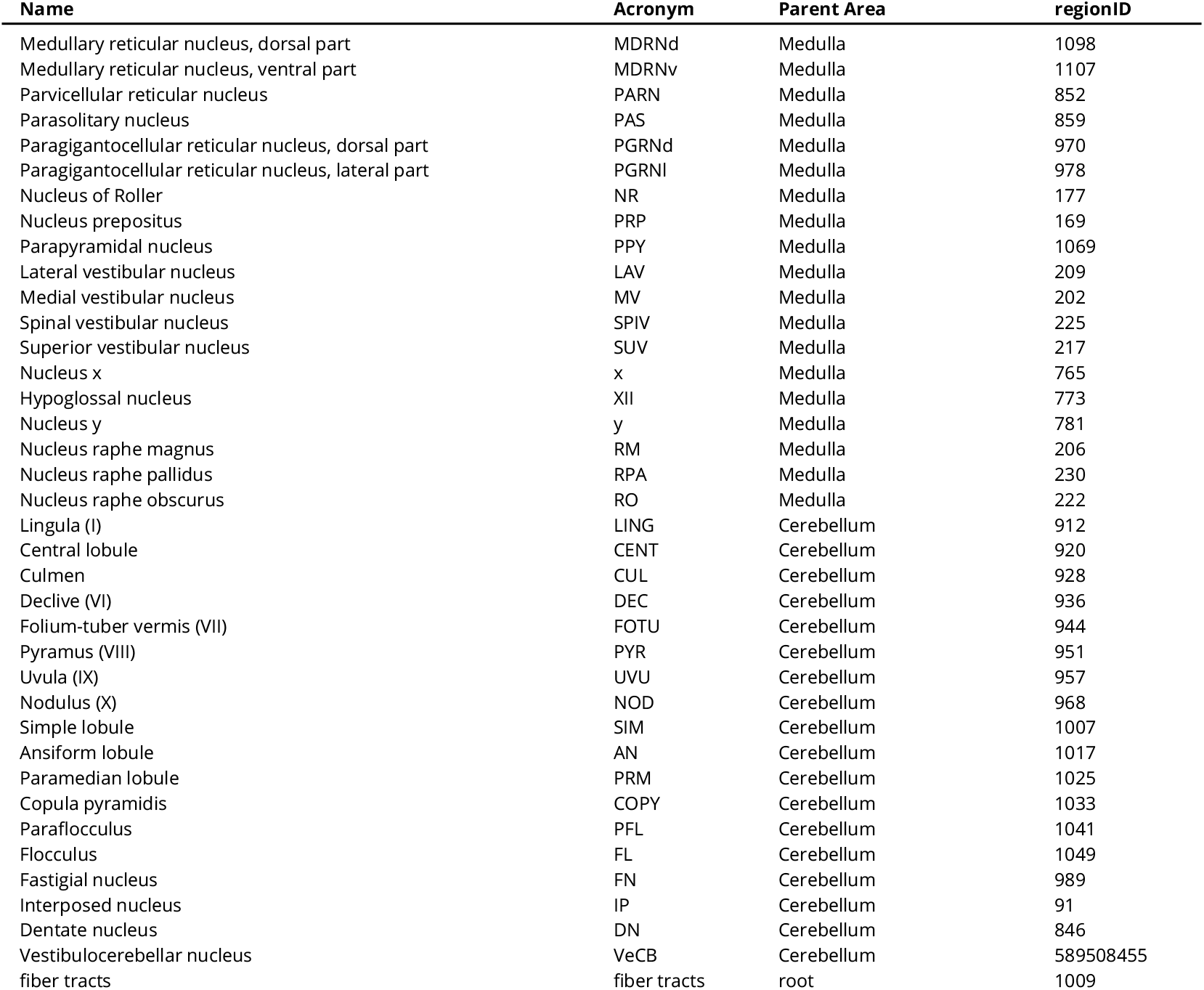
Mid-ontology brain regions.

### Supplementary Data

#### Supplementary data SD1

##### Whole-brain PNNs metrics

This .xlsx file contains tables with quantitative measurements for PNN density, WFA diffuse fluorescence, PNN intensity, and PNN energy for all brain areas. Data are presented at three levels of resolution: coarse, medium, and fine. For each resolution level, we report data from each mouse and the mean values across all mice in separate sheets.

#### Supplementary data SD2

##### Whole-brain PV-positive cells metrics

This .xlsx file contains tables with quantitative measurements for PV cell density, PV diffuse fluorescence, PV cell intensity, and PV energy for all brain areas. Data are presented at three levels of resolution: coarse, medium, and fine. For each level, we report data from each mouse and the mean values across all mice in separate sheets.

#### Supplementary data SD3

##### Whole-brain PNN-PV colocalization metrics

This .xlsx file contains tables with the percentage of PNNs ensheathing a PV cell (`pvPositive_pnn`) and the percentage of PV cells surrounded by a PNN (`wfaPositive_pv`), for all brain areas. Data are presented at three levels of resolution: coarse, medium, and fine, in separate sheets. For the coarse resolution level, we report data from each mouse and the mean values across all mice. For medium and fine and resolution levels, we report data from each experimental unit (indicated with the identifier of the animal that was excluded, see section Colocalization PNN-PV in Methods & Materials for details) as well as the mean values across all experimental units. Only areas with at least 3 PNNs and 3 PV cells in at least 4 animals are included.

#### Supplementary data SD4

##### Correlation of staining metrics with gene expression

This .xlsx file contains tables with the results of the correlation analysis of the staining metrics in our dataset with the gene expression data published in the Allen Institute Anatomic Gene Expression Atlas (AGEA). Correlations with PNN energy, PV energy, and WFA diffuse fluorescence are reported in separate sheets. Each gene is referred to with the acronym, the ID in the AGEA, the Entrez ID, and the full name. For each gene, we report the Spearman correlation coefficient, the correspondent p-value, the false discovery rate (FDR), and the Bonferroni-adjusted p-value.

## Notes

https://zenodo.org/record/7419283

https://github.com/LeonardoLupori/wholeBrain_PNN_analysis

### Competing Interest Statement

The authors have declared no competing interest.

## References

Ariza, Jeanelle, Haille Rogers, Ezzat Hashemi, Stephen C Noctor, and Verónica Martínez-Cerdeño (2018). “The Number of Chandelier and Basket Cells Are Differentially Decreased in Prefrontal Cortex in Autism”. In: Cerebral Cortex 28.2, pp. 411–420. DOI: 10.1093/cercor/bhw349 (cit. on p.13).

Baimbridge, K. G. and J. J. Miller (1982). “Immunohistochemical localization of calcium-binding protein in the cerebellum, hippocampal formation and olfactory bulb of the rat”. In: Brain Research 245.2, pp. 223–229. DOI: 10.1016/0006-8993(82)90804-6 (cit. on p. 6).

Bastianelli, Enrico (2003). “Distribution of calcium-binding proteins in the cerebellum”. In: The Cerebellum 2.4, pp. 242–262. DOI: 10.1080/14734220310022289 (cit. on p. 6).

Berg, Stuart, Dominik Kutra, Thorben Kroeger, Christoph N. Straehle, Bernhard X. Kausler, Carsten Haubold, Martin Schiegg, Janez Ales, Thorsten Beier, Markus Rudy, Kemal Eren, Jaime I. Cervantes, Buote Xu, Fynn Beuttenmueller, Adrian Wolny, Chong Zhang, Ullrich Koethe, Fred A. Hamprecht, and Anna Kreshuk (2019). “ilastik: interactive machine learning for (bio)image analysis”. In: Nature Methods 16.12, pp. 1226–1232. DOI: 10.1038/s41592-019-0582-9 (cit. on p. 17).

Beurdeley, Marine, Julien Spatazza, Henry H. C. Lee, Sayaka Sugiyama, Clémence Bernard, Ariel A. Di Nardo, Takao K. Hensch, and Alain Prochiantz (2012). “Otx2 Binding to Perineuronal Nets Persistently Regulates Plasticity in the Mature Visual Cortex”. In: Journal of Neuroscience 32.27, pp. 9429–9437. DOI: 10.1523/JNEUROSCI.0394-12.2012 (cit. on p. 2).

Bjerke, Ingvild E., Sharon C. Yates, Arthur Laja, Menno P. Witter, Maja A. Puchades, Jan G. Bjaalie, and Trygve B. Leergaard (2021). “Densities and numbers of calbindin and parvalbumin positive neurons across the rat and mouse brain”. In: iScience 24.1, p. 101906. DOI: 10.1016/j.isci.2020.101906 (cit. on p. 5).

Boggio, Elena Maria, Erich M. Ehlert, Leonardo Lupori, Elizabeth B. Moloney, Fred De Winter, Craig W. Vander Kooi, Laura Baroncelli, Vasilis Mecollari, Bas Blits, James W. Fawcett, Joost Verhaagen, and Tommaso Pizzorusso (2019). “Inhibition of Semaphorin3A Promotes Ocular Dominance Plasticity in the Adult Rat Visual Cortex”. In: Molecular Neurobiology 56.9, pp. 5987–5997. DOI: 10.1007/s12035-019-1499-0 (cit. on p. 1).

Boghdadi, Anthony G., Leon Teo, and James A. Bourne (2018). “The Involvement of the Myelin-Associated Inhibitors and Their Receptors in CNS Plasticity and Injury”. In: Molecular Neurobiology 55.3, pp. 1831–1846. DOI: 10.1007/s12035-017-0433-6 (cit. on p. 15).

Bonetto, Giulia, David Belin, and Ragnhildur Thóra Káradóttir (2021). “Myelin: A gatekeeper of activity-dependent circuit plasticity?” In: Science 374.6569, eaba6905. DOI: 10.1126/science.aba6905 (cit. on p. 15).

Bouhours, Brice, Enida Gjoni, Olexiy Kochubey, and Ralf Schneggenburger (2017). “Synaptotagmin2 (Syt2) Drives Fast Release Redundantly with Syt1 at the Output Synapses of Parvalbumin-Expressing Inhibitory Neurons”. In: The Journal of Neuroscience: The Official Journal of the Society for Neuroscience 37.17, pp. 4604–4617. DOI: 10.1523/JNEUROSCI.3736-16.2017 (cit. on p. 10).

Bradbury, Elizabeth J., Lawrence D. F. Moon, Reena J. Popat, Von R. King, Gavin S. Bennett, Preena N. Patel, James W. Fawcett, and Stephen B. McMahon (2002). “Chondroitinase ABC promotes functional recovery after spinal cord injury”. In: Nature 416.6881, pp. 636–640. DOI: 10.1038/416636a (cit. on p. 2).

Brand, Amy, Liz Allen, Micah Altman, Marjorie Hlava, and Jo Scott (2015). “Beyond authorship: attribution, contribution, collaboration, and credit”. In: Learned Publishing 28.2, pp. 151–155. DOI: 10.1087/20150211 (cit. on p. 22).

Bukalo, O., M. Schachner, and A. Dityatev (2007). “Hippocampal Metaplasticity Induced by Deficiency in the Extracellular Matrix Glycoprotein Tenascin-R”. In: Journal of Neuroscience 27.22, pp. 6019–6028. DOI: 10.1523/JNEUROSCI.1022-07.2007 (cit. on p. 2).

Cabungcal, Jan-Harry, Pascal Steullet, Hirofumi Morishita, Rudolf Kraftsik, Michel Cuenod, Takao K. Hensch, and Kim Q. Do (2013). “Perineuronal nets protect fast-spiking interneurons against oxidative stress”. In: Proceedings of the National Academy of Sciences 110.22, pp. 9130–9135. DOI: 10.1073/pnas.1300454110 (cit. on p. 2).

Carstens, Kelly E., Mary L. Phillips, Lucas Pozzo-Miller, Richard J. Weinberg, and Serena M. Dudek (2016). “Perineuronal Nets Suppress Plasticity of Excitatory Synapses on CA2 Pyramidal Neurons”. In: Journal of Neuroscience 36.23, pp. 6312–6320. DOI: 10.1523/JNEUROSCI.0245-16.2016 (cit. on p. 2).

Carter, Brett C. and Bruce P. Bean (2009). “Sodium Entry during Action Potentials of Mammalian Neurons: Incomplete Inactivation and Reduced Metabolic Efficiency in Fast-Spiking Neurons”. In: Neuron 64.6, pp. 898–909. DOI: 10.1016/j.neuron.2009.12.011 (cit. on p. 15).

Carulli, Daniela, Robin Broersen, Fred de Winter, Elizabeth M. Muir, Maja Mešković, Matthijs de Waal, Sharon de Vries, Henk-Jan Boele, Cathrin B. Canto, Chris I. De Zeeuw, and Joost Verhaagen (2020). “Cerebellar plasticity and associative memories are controlled by perineuronal nets”. In: Proceedings of the National Academy of Sciences of the United States of America 117.12, pp. 6855–6865. DOI: 10.1073/pnas.1916163117 (cit. on p. 2).

Carulli, Daniela, Tommaso Pizzorusso, Jessica C. F. Kwok, Elena Putignano, Andrea Poli, Serhiy Forostyak, Melissa R. Andrews, Sathyaseelan S. Deepa, Tibor T. Glant, and James W. Fawcett (2010). “Animals lacking link protein have attenuated perineuronal nets and persistent plasticity”. In: Brain 133.8, pp. 2331–2347. DOI: 10.1093/brain/awq145 (cit. on pp. 2,10).

Chow, A., A. Erisir, C. Farb, M. S. Nadal, A. Ozaita, D. Lau, E. Welker, and B. Rudy (1999). “K+ Channel Expression Distinguishes Subpopulations of Parvalbumin-and Somatostatin-Containing Neocortical Interneurons”. In: The Journal of Neuroscience 19.21, pp. 9332–9345. DOI: 10.1523/JNEUROSCI.19-21-09332.1999 (cit. on p. 10).

Christensen, Ane Charlotte, Kristian Kinden Lensjø, Mikkel Elle Lepperød, Svenn-Arne Dragly, Halvard Sutterud, Jan Sigurd Blackstad, Marianne Fyhn, and Torkel Hafting (2021). “Perineuronal nets stabilize the grid cell network”. In: Nature Communications 12.1, p. 253. DOI: 10.1038/s41467-020-20241-w (cit. on p. 2).

Ciampi, Luca, Fabio Carrara, Valentino Totaro, Raffaele Mazziotti, Leonardo Lupori, Carlos Santiago, Giuseppe Amato, Tommaso Pizzorusso, and Claudio Gennaro (2022). “Learning to count biological structures with raters’ uncertainty”. In: Medical Image Analysis 80, p. 102500. DOI: 10.1016/j.media.2022.102500 (cit. on pp. 3, 17).

Cicconet, Marcelo and Daniel R. Hochbaum (2019). A Supervised, Symmetry-Driven, GUI Toolkit for Mouse Brain Stack Registration and Plane Assignment. preprint. Bioinformatics. DOI: 10.1101/781880 (cit. on p. 19).

Claudi, Federico and Luigi Petrucco (2022). brainglobe/bg-heatmaps: DOI: 10.5281/zenodo.5891814 (cit. on p. 21).

Claudi, Federico, Adam LTyson, Luigi Petrucco, Troy W Margrie, Ruben Portugues, and Tiago Branco (2021). “Visualizing anatomically registered data with brainrender”. In: eLife 10. Ed. by Mackenzie W Mathis, Kate M Wassum, and Juan Nunez-Iglesias, e65751. DOI: 10.7554/eLife.65751 (cit. on p.21).

Cope, Elise C., Anna D. Zych, Nicole J. Katchur, Renée C. Waters, Blake J. Laham, Emma J. Diethorn, Christin Y. Park, William R. Meara, and Elizabeth Gould (2021). “Atypical perineuronal nets in the CA2 region interfere with social memory in a mouse model of social dysfunction”. In: Molecular Psychiatry, pp. 1–12. DOI: 10.1038/s41380-021-01174-2 (cit. on p. 2).

Dauth, Stephanie, Thomas Grevesse, Harry Pantazopoulos, Patrick H. Campbell, Ben M. Maoz, Sabina Berretta, and Kevin Kit Parker (2016). “Extracellular matrix protein expression is brain region dependent”. In: Journal of Comparative Neurology 524.7, pp. 1309–1336. DOI: 10.1002/cne.23965 (cit. on pp. 2, 10, 13).

Domínguez, Soledad, Christophe Clément Rey, Ludivine Therreau, Aurélien Fanton, Dominique Massotte, Laure Verret, Rebecca Ann Piskorowski, and Vivien Chevaleyre (2019). “Maturation of PNN and ErbB4 Signaling in Area CA2 during Adolescence Underlies the Emergence of PV Interneuron Plasticity and Social Memory”. In: Cell Reports 29.5, 1099–1112.e4. DOI: 10.1016/j.celrep.2019.09.044 (cit. on p. 2).

Donato, Flavio, Ananya Chowdhury, Maria Lahr, and Pico Caroni (2015). “Early-and Late-Born Parvalbumin Basket Cell Subpopulations Exhibiting Distinct Regulation and Roles in Learning”. In: Neuron 85.4, pp. 770–786. DOI: 10.1016/j.neuron.2015.01.011 (cit. on pp. 7, 14).

Donato, Flavio, Santiago Belluco Rompani, and Pico Caroni (2013). “Parvalbumin-expressing basket-cell network plasticity induced by experience regulates adult learning”. In: Nature 504.7479, pp. 272–276. DOI: 10.1038/nature12866 (cit. on pp. 7, 14).

Faini, Giulia, Andrea Aguirre, Silvia Landi, Didi Lamers, Tommaso Pizzorusso, Gian Michele Ratto, Charlotte Deleuze, and Alberto Bacci (2018). “Perineuronal nets control visual input via thalamic recruitment of cortical PV interneurons”. In: eLife 7. Ed. by Marlene Bartos and Gary L Westbrook, e41520. DOI: 10.7554/eLife.41520 (cit. on pp. 9, 15).

Fawcett, James W., Marianne Fyhn, Pavla Jendelova, Jessica C. F. Kwok, Jiri Ruzicka, and Barbara A. Sorg (2022). “The extracellular matrix and perineuronal nets in memory”. In: Molecular Psychiatry 27.8, pp. 3192–3203. DOI: 10.1038/s41380-022-01634-3 (cit. on p. 2).

Fawcett, James W., Toshitaka Oohashi, and Tommaso Pizzorusso (2019). “The roles of perineuronal nets and the perinodal extracellular matrix in neuronal function”. In: Nature Reviews. Neuro-science 20.8, pp. 451–465. DOI: 10.1038/s41583-019-0196-3 (cit. on pp. 2, 7, 10, 11, 13-15).

Fishell, Gordon (2008). “Perspectives on the Developmental Origins of Cortical Interneuron Diversity”. In: Cortical Development: Genes and Genetic Abnormalities. John Wiley & Sons, Ltd, pp. 21–44. DOI: 10.1002/9780470994030.ch3 (cit. on p. 14).

Galtrey, Clare M., Jessica C. F. Kwok, Daniela Carulli, Kate E. Rhodes, and James W. Fawcett (2008). “Distribution and synthesis of extracellular matrix proteoglycans, hyaluronan, link proteins and tenascin-R in the rat spinal cord”. In: European Journal of Neuroscience 27.6, pp. 1373–1390. DOI: 10.1111/j.1460-9568.2008.06108.x (cit. on pp. 1, 13).

Gibel-Russo, Rachel, David Benacom, and Ariel A. Di Nardo (2022). “Non-Cell-Autonomous Factors Implicated in Parvalbumin Interneuron Maturation and Critical Periods”. In: Frontiers in Neural Circuits 16 (cit. on p. 14).

Gogolla, Nadine, Pico Caroni, Andreas Lüthi, and Cyril Herry (2009). “Perineuronal Nets Protect Fear Memories from Erasure”. In: Science 325.5945, pp. 1258–1261. DOI: 10.1126/science.1174146 (cit. on p. 2).

Harris, Charles R. et al. (2020). “Array programming with NumPy”. In: Nature 585.7825, pp. 357–362. DOI: 10.1038/s41586-020-2649-2 (cit. on p. 19).

Härtig, Wolfgang, Amin Derouiche, Klaus Welt, Kurt Brauer, Jens Grosche, Michael Mäder, Andreas Reichenbach, and Gert Brückner (1999). “Cortical neurons immunoreactive for the potassium channel Kv3.1b subunit are predominantly surrounded by perineuronal nets presumed as a buffering system for cations”. In: Brain Research 842.1, pp. 15–29. DOI: 10.1016/S0006-8993(99)01784-9 (cit. on p. 2).

Härtig, Wolfgang, Anton Meinicke, Dominik Michalski, Stefan Schob, and Carsten Jäger (2022). “Update on Perineuronal Net Staining With Wisteria floribunda Agglutinin (WFA)”. In: Frontiers in Integrative Neuroscience 16, p. 851988. DOI: 10.3389/fnint.2022.851988 (cit. on p. 10).

Held-Feindt, Janka, Elke Bernedo Paredes, Ulrike Blömer, Constanze Seidenbecher, Andreas M. Stark, H. Maximilian Mehdorn, and Rolf Mentlein (2006). “Matrix-degrading proteases ADAMTS4 and ADAMTS5 (disintegrins and metalloproteinases with thrombospondin motifs 4 and 5) are expressed in human glioblastomas”. In: International Journal of Cancer 118.1, pp. 55–61. DOI: 10.1002/ijc.21258 (cit. on p. 10).

Hendry, S. H., E. G. Jones, S. Hockfield, and R. D. McKay (1988). “Neuronal populations stained with the monoclonal antibody Cat-301 in the mammalian cerebral cortex and thalamus”. In:Journal of Neuroscience 8.2, pp. 518–542. DOI: 10.1523/JNEUROSCI.08-02-00518.1988 (cit. on p. 1).

Huber, Peter J. and Elvezio M. Ronchetti (2009). “Robust Statistics, Concomitant scale estimates”. In: Robust Statistics. Second Edition. John Wiley & Sons (cit. on p. 20).

Hunter, John D. (2007). “Matplotlib: A 2D Graphics Environment”. In: Computing in Science & Engineering 9.3, pp. 90–95. DOI: 10.1109/MCSE.2007.55 (cit. on p. 21).

Hunyadi, Andrea, Botond Gaál, Clara Matesz, Zoltan Meszar, Markus Morawski, Katja Reimann, David Lendvai, Alan Alpar, Ildikó Wéber, and Éva Rácz (2020). “Distribution and classification of the extracellular matrix in the olfactory bulb”. In: Brain Structure and Function 225.1, pp. 321–344. DOI: 10.1007/s00429-019-02010-8 (cit. on pp. 14, 18).

Jaini, K. Anil and Farshid Farrokhnia (1991). “Unsupervised texture segmentation using Gabor fil-ters”. In: 24.12, pp. 1167–1186. DOI: https://doi.org/10.1016/0031-3203(91)90143-S (cit. on p.19).

Kann, Oliver, Ismini E Papageorgiou, and Andreas Draguhn (2014). “Highly Energized Inhibitory Interneurons are a Central Element for Information Processing in Cortical Networks”. In: Journal of Cerebral Blood Flow & Metabolism 34.8, pp. 1270–1282. DOI: 10.1038/jcbfm.2014.104 (cit. on p.15).

Kim, Yongsoo, Guangyu Robert Yang, Kith Pradhan, Kannan Umadevi Venkataraju, Mihail Bota, Luis Carlos García del Molino, Greg Fitzgerald, Keerthi Ram, Miao He, Jesse Maurica Levine, Partha Mitra, Z. Josh Huang, Xiao-Jing Wang, and Pavel Osten (2017). “Brain-wide Maps Reveal Stereo-typed Cell-Type-Based Cortical Architecture and Subcortical Sexual Dimorphism”. In: Cell 171.2, 456–469.e22. DOI: 10.1016/j.cell.2017.09.020 (cit. on pp. 5, 9).

Köppe, Gerlinde, Gert Brückner, Kurt Brauer, Wolfgang Härtig, and Volker Bigl (1997). “Developmental patterns of proteoglycan-containing extracellular matrix in perineuronal nets and neuropil of the postnatal rat brain”. In: Cell and Tissue Research 288.1, pp. 33–41. DOI: 10.1007/s004410050790 (cit. on p. 1).

Kwok, Jessica C. F., Daniela Carulli, and James W. Fawcett (2010). “In vitro modeling of perineuronal nets: hyaluronan synthase and link protein are necessary for their formation and integrity”. In: Journal of Neurochemistry 114.5, pp. 1447–1459. DOI: 10.1111/j.1471-4159.2010.06878.x (cit. on pp.2, 10).

Lee, H. H. C., C. Bernard, Z. Ye, D. Acampora, A. Simeone, A. Prochiantz, A. A. Di Nardo, and T. K. Hensch (2017). “Genetic Otx2 mis-localization delays critical period plasticity across brain regions”. In: Molecular Psychiatry 22.5, pp. 680–688. DOI: 10.1038/mp.2017.1 (cit. on p. 14).

Lein, Ed S. et al. (2007). “Genome-wide atlas of gene expression in the adult mouse brain”. In: Nature 445.7124, pp. 168–176. DOI: 10.1038/nature05453 (cit. on pp. 4, 10, 13, 15, 20, 21).

Liu, Cheng-Hang, Arnold J. Heynen, Marshall G. Hussain Shuler, and Mark F. Bear (2008). “Cannabinoid Receptor Blockade Reveals Parallel Plasticity Mechanisms in Different Layers of Mouse Visual Cortex”. In: Neuron 58.3, pp. 340–345. DOI: 10.1016/j.neuron.2008.02.020 (cit. on p. 15).

Lorincz, Andrea and Zoltan Nusser (2008). “Cell-Type-Dependent Molecular Composition of the Axon Initial Segment”. In: Journal of Neuroscience 28.53, pp. 14329–14340. DOI: 10.1523/JNEUROSCI.4833-08.2008 (cit. on p. 10).

Lupori, Leonardo, Valentino Totaro, Sara Cornuti, Luca Ciampi, Fabio Carrara, Edda Grilli, Aurelia Viglione, Francesca Tozzi, Elena Putignano, Raffaele Mazziotti, Giuseppe Amato, Claudio Gennaro, Paola Toglini, and Tommaso Pizzorusso (2022). A brain-wide, annotated dataset of WFA-positive perineuronal nets and parvalbumin neurons in the adult mouse brain. DOI: 10.5281/zenodo.7419283 (cit. on p. 13).

Matthews, Russell T., Gail M. Kelly, Cynthia A. Zerillo, Grace Gray, Michael Tiemeyer, and Susan Hockfield (2002). “Aggrecan glycoforms contribute to the molecular heterogeneity of perineuronal nets”. In: The Journal of Neuroscience: The Official Journal of the Society for Neuroscience 22.17, pp. 7536–7547 (cit. on p. 13).

McKinney, Wes (2010). “Data Structures for Statistical Computing in Python”. In: Python in Science Conference. Austin, Texas, pp. 56–61. DOI: 10.25080/Majora-92bf1922-00a (cit. on p. 19).

Miyata, Shinji, Yukio Komatsu, Yumiko Yoshimura, Choji Taya, and Hiroshi Kitagawa (2012). “Persistent cortical plasticity by upregulation of chondroitin 6-sulfation”. In: Nature Neuroscience 15.3, pp. 414–422. DOI: 10.1038/nn.3023 (cit. on p. 2).

Naba, Alexandra, Karl R. Clauser, Huiming Ding, Charles A. Whittaker, Steven A. Carr, and Richard O. Hynes (2016). “The extracellular matrix: Tools and insights for the “omics” era”. In: Matrix Biology 49, pp. 10–24. DOI: 10.1016/j.matbio.2015.06.003 (cit. on pp. 11, 21).

Nabel, Elisa and Hirofumi Morishita (2013). “Regulating Critical Period Plasticity: Insight from the Visual System to FearCircuitryforTherapeutic Interventions”. In: Frontiers in Psychiatry 4 (cit. on p.2).

Ogiwara, Ikuo, Hiroyuki Miyamoto, Noriyuki Morita, Nafiseh Atapour, Emi Mazaki, Ikuyo Inoue, Tamaki Takeuchi, Shigeyoshi Itohara, Yuchio Yanagawa, Kunihiko Obata, Teiichi Furuichi, Takao K. Hensch, and Kazuhiro Yamakawa (2007). “Nav1.1 Localizes to Axons of Parvalbumin-Positive Inhibitory Interneurons: A Circuit Basis for Epileptic Seizures in Mice Carrying an Scn1a Gene Mutation”. In: The Journal of Neuroscience 27.22, pp. 5903–5914. DOI: 10.1523/JNEUROSCI.5270-06.2007 (cit. on p. 10).

Oh, Seung Wook et al. (2014). “A mesoscale connectome of the mouse brain”. In: Nature 508.7495, pp. 207–214. DOI: 10.1038/nature13186 (cit. on pp. 9, 13, 20).

Oohashi, Toshitaka, Midori Edamatsu, Yoko Bekku, and Daniela Carulli (2015). “The hyaluronan and proteoglycan link proteins: Organizers of the brain extracellular matrix and key molecules for neuronal function and plasticity”. In: Experimental Neurology. Deciphering sugar chain-based signals regulating integrative neuronal functions 274, pp. 134–144. DOI: 10.1016/j.expneurol.2015.09.010 (cit. on p. 10).

Pantazopoulos, Harry, Tsung-Ung W. Woo, Maribel P. Lim, Nicholas Lange, and Sabina Berretta (2010). “Extracellular matrix-glial abnormalities in the amygdala and entorhinal cortex of subjects diagnosed with schizophrenia”. In: Archives of General Psychiatry 67.2, pp. 155–166. DOI: 10.1001/archgenpsychiatry.2009.196 (cit. on p. 2).

Pedregosa, Fabian, Gaël Varoquaux, Alexandre Gramfort, Vincent Michel, Bertrand Thirion, Olivier Grisel, Mathieu Blondel, Peter Prettenhofer, Ron Weiss, Vincent Dubourg, Jake Vanderplas, Alexandre Passos, David Cournapeau, Matthieu Brucher, Matthieu Perrot, and Édouard Duchesnay (2011). “Scikit-learn: Machine Learning in Python”. In: Journal of Machine Learning Research 12.85, pp. 2825–2830 (cit. on p. 19).

Pirbhoy, Patricia S., Maham Rais, Jonathan W. Lovelace, Walker Woodard, Khaleel A. Razak, Devin K. Binder, and Iryna M. Ethell (2020). “Acute pharmacological inhibition of matrix metalloproteinase-9 activity during development restores perineuronal net formation and normalizes auditory processing in Fmr1 KO mice”. In: Journal of Neurochemistry 155.5, pp. 538–558. DOI: 10.1111/jnc.15037 (cit. on p. 10).

Pizzorusso, Tommaso, Paolo Medini, Nicoletta Berardi, Sabrina Chierzi, James W. Fawcett, and Lamberto Maffei (2002). “Reactivation of Ocular Dominance Plasticity in the Adult Visual Cortex”. In: Science 298.5596, pp. 1248–1251. DOI: 10.1126/science.1072699 (cit. on pp. 1, 2).

Puchades, Maja A., Gergely Csucs, Debora Ledergerber, Trygve B. Leergaard, and Jan G. Bjaalie (2019). “Spatial registration of serial microscopic brain images to three-dimensional reference atlases with the QuickNII tool”. In: PLOS ONE 14.5, e0216796. DOI: 10.1371/journal.pone.0216796 (cit. on p. 17).

Reichelt, Amy C., Dominic J. Hare, Timothy J. Bussey, and Lisa M. Saksida (2019). “Perineuronal Nets: Plasticity, Protection, and Therapeutic Potential”. In: Trends in Neurosciences 42.7, pp. 458–470. DOI: 10.1016/j.tins.2019.04.003 (cit. on pp. 1,14).

Ren, Shaoqing, Kaiming He, Ross Girshick, and Jian Sun (2015). “Faster R-CNN: Towards Real-Time Object Detection with Region Proposal Networks”. In: Advances in Neural Information Processing Systems. Vol. 28. Curran Associates, Inc. (cit. on p. 17).

Romberg, C., S. Yang, R. Melani, M. R. Andrews, A. E. Horner, M. G. Spillantini, T. J. Bussey, J. W. Fawcett, T. Pizzorusso, and L. M. Saksida (2013). “Depletion of Perineuronal Nets Enhances Recognition Memory and Long-Term Depression in the Perirhinal Cortex”. In: Journal of Neuroscience 33.16, pp. 7057–7065. DOI: 10.1523/JNEUROSCI.6267-11.2013 (cit. on p. 2).

Rossier, J., A. Bernard, J.-H. Cabungcal, Q. Perrenoud, A. Savoye, T. Gallopin, M. Hawrylycz, M. Cuénod, K. Do, A. Urban, and Ed S. Lein (2015). “Cortical fast-spiking parvalbumin interneurons enwrapped in the perineuronal net express the metallopeptidases Adamts8, Adamts15 and Neprilysin”. In: Molecular Psychiatry 20.2, pp. 154–161. DOI: 10.1038/mp.2014.162 (cit. on p. 10).

Rowlands, Daire, Kristian K. Lensjø, Tovy Dinh, Sujeong Yang, Melissa R. Andrews, Torkel Hafting, Marianne Fyhn, James W. Fawcett, and Gunnar Dick (2018). “Aggrecan Directs Extracellular Matrix-Mediated Neuronal Plasticity”. In: Journal of Neuroscience 38.47, pp. 10102–10113. DOI: 10.1523/JNEUROSCI.1122-18.2018 (cit. on p. 2).

Ruder, Ludwig, Riccardo Schina, Harsh Kanodia, Sara Valencia-Garcia, Chiara Pivetta, and Silvia Arber (2021). “Afunctional map for diverse forelimb actions within brainstem circuitry”. In: Nature 590.7846, pp. 445–450. DOI: 10.1038/s41586-020-03080-z (cit. on p. 14).

Saladin, Kenneth S., Christina A. Gan, and Heather N. Cushman (2021). Anatomy & Physiology: The Unity of Form and Function. McGraw-Hill Education. Dimensions 23.1×29.0×4.3 cm, book (cit. on p.14).

Seeger, G., K. Brauer, W. Härtig, and G. Brückner (1994). “Mapping of perineuronal nets in the rat brain stained by colloidal iron hydroxide histochemistry and lectin cytochemistry”. In: Neuroscience 58.2, pp. 371–388. DOI: 10.1016/0306-4522(94)90044-2 (cit. on p. 1).

Tasic, Bosiljka et al. (2016). “Adult mouse cortical cell taxonomy revealed by single cell transcriptomics”. In: Nature Neuroscience 19.2, pp. 335–346. DOI: 10.1038/nn.4216 (cit. on p. 13).

Trachtenberg, Joshua T., Christopher Trepel, and Michael P. Stryker (2000). “Rapid Extragranular Plasticity in the Absence of Thalamocortical Plasticity in the Developing Primary Visual Cortex”. In: Science 287.5460, pp. 2029–2032. DOI: 10.1126/science.287.5460.2029 (cit. on p. 15).

Ueno, Hiroshi, Kazuki Fujii, Shunsuke Suemitsu, Shinji Murakami, Naoya Kitamura, Kenta Wani, Shozo Aoki, Motoi Okamoto, Takeshi Ishihara, and Keizo Takao (2018). “Expression of aggrecan components in perineuronal nets in the mouse cerebral cortex”. In: IBRO Reports 4, pp. 22–37. DOI: 10.1016/j.ibror.2018.01.002 (cit. on pp. 2, 10, 13).

Virtanen, Pauli et al. (2020). “SciPy 1.0: fundamental algorithms for scientific computing in Python”. In: Nature Methods 17.3, pp. 261–272. DOI: 10.1038/s41592-019-0686-2 (cit. on p. 19).

Wang, Quanxin et al. (2020). “The Allen Mouse Brain Common Coordinate Framework: A 3D Reference Atlas”. In: Cell 181.4, 936–953.e20. DOI: 10.1016/j.cell.2020.04.007 (cit. on p. 17).

Waskom, Michael L. (2021). “seaborn: statistical data visualization”. In: Journal of Open Source Software 6.60, p. 3021. DOI: 10.21105/joss.03021 (cit. on p. 21).

Wingert, Jereme C. and Barbara A. Sorg (2021). “Impact of Perineuronal Nets on Electrophysiology of Parvalbumin Interneurons, Principal Neurons, and Brain Oscillations: A Review”. In: Frontiers in Synaptic Neuroscience 13, p. 673210. DOI: 10.3389/fnsyn.2021.673210 (cit. on p. 2).

Wong, Tzu-Tsung (2015). “Performance evaluation of classification algorithms by k-fold and leave-one-out cross validation”. In: Pattern Recognition 48.9, pp. 2839–2846. DOI: 10.1016/j.patcog.2015.03.009 (cit. on p. 20).

Yamada, Jun and Shozo Jinno (2017). “Molecular heterogeneity of aggrecan-based perineuronal nets around five subclasses of parvalbumin-expressing neurons in the mouse hippocampus”. In: Journal of Comparative Neurology 525.5, pp. 1234–1249. DOI: 10.1002/cne.24132 (cit. on p. 10).

Ye, Qian and Qing-long Miao (2013). “Experience-dependent development of perineuronal nets and chondroitin sulfate proteoglycan receptors in mouse visual cortex”. In: Matrix Biology 32.6, pp. 352–363. DOI: 10.1016/j.matbio.2013.04.001 (cit. on p. 1).

Zhang, Bing, Stefan Kirov, and Jay Snoddy (2005). “WebGestalt: an integrated system for exploring gene sets in various biological contexts”. In: Nucleic Acids Research 33 (suppl_2), W741–W748. DOI: 10.1093/nar/gki475 (cit. on p. 21).

Zhao, Shitao, Jianqiang Sun, Kentaro Shimizu, and Koji Kadota (2018). “Silhouette Scores for Arbitrary Defined Groups in Gene Expression Data and Insights into Differential Expression Results”. In: Biological Procedures Online 20.1, p. 5. DOI: 10.1186/s12575-018-0067-8 (cit. on p. 10).

Zingg, Brian, Houri Hintiryan, Lin Gou, Monica Y. Song, Maxwell Bay, Michael S. Bienkowski, Nicholas N. Foster, Seita Yamashita, Ian Bowman, Arthur W. Toga, and Hong-Wei Dong (2014). “Neural Networks of the Mouse Neocortex”. In: Cell 156.5, pp. 1096–1111. DOI: 10.1016/j.cell.2014.02.023 (cit. on pp. 9, 13).

